# The Foundational data initiative for Parkinson’s disease (FOUNDIN-PD): enabling efficient translation from genetic maps to mechanism

**DOI:** 10.1101/2021.06.03.446785

**Authors:** Elisangela Bressan, Xylena Reed, Vikas Bansal, Elizabeth Hutchins, Melanie M. Cobb, Michelle G Webb, Eric Alsop, Francis P. Grenn, Anastasia Illarionova, Natalia Savytska, Ivo Violich, Stefanie Broeer, Noémia Fernandes, Ramiyapriya Sivakumar, Alexandra Beilina, Kimberley Billingsley, Joos Berghausen, Caroline B. Pantazis, Vanessa Pitz, Dhairya Patel, Kensuke Daida, Bessie Meechoovet, Rebecca Reiman, Amanda Courtright-Lim, Amber Logemann, Jerry Antone, Mariya Barch, Robert Kitchen, Yan Li, Clifton L. Dalgard, The American Genome Center, Patrizia Rizzu, Dena G Hernandez, Brooke E. Hjelm, Mike Nalls, J. Raphael Gibbs, Steven Finkbeiner, Mark R Cookson, Kendall Van Keuren-Jensen, David W Craig, Andrew B Singleton, Peter Heutink, Cornelis Blauwendraat

## Abstract

The FOUNdational Data INitiative for Parkinson’s Disease (FOUNDIN-PD) is an international collaboration producing fundamental resources for Parkinson’s disease (PD). FOUNDIN-PD generated a multi-layered molecular dataset in a cohort of induced pluripotent stem cell (iPSC) lines differentiated to dopaminergic (DA) neurons, a major affected cell type in PD. The lines were derived from the Parkinson’s Progression Markers Initiative study including participants with PD carrying monogenic PD (*SNCA*) variants, variants with intermediate effects and variants identified by genome-wide association studies and unaffected individuals. We generated genetic, epigenetic, regulatory, transcriptomic, and longitudinal cellular imaging data from iPSC-derived DA neurons to understand molecular relationships between disease associated genetic variation and proximate molecular events. These data reveal that iPSC-derived DA neurons provide a valuable cellular context and foundational atlas for modelling PD genetic risk. We have integrated these data into a FOUNDIN-PD data browser (https://www.foundinpd.org) as a resource for understanding the molecular pathogenesis of PD.

## Introduction

Our understanding of the genetic architecture of Parkinson’s disease (PD) has expanded considerably over the last decade. Investigations of rare monogenic forms of PD and parkinsonism have revealed multiple genes that contain disease-causing mutations ^1^. Additionally, iterative application of genome wide association studies (GWAS) in increasingly larger sample sizes have identified 90 independent risk variants for PD, which cumulatively contribute to 16-36% of the heritable risk for the disease ^2^. One of the main pathological hallmarks of PD is the progressive degeneration of dopaminergic (DA) neurons in the substantia nigra and the accumulation of alpha-synuclein protein aggregates, known as Lewy bodies and Lewy neurites ^3^. Additionally, previous work has highlighted that genetic risk in PD is likely to play a significant role in DA) neurons ^2,4^.

On a clinical level, there is large variability in age at onset and progression across both monogenic and idiopathic PD patients, even in those carrying the same damaging variant. This variation is likely caused by a combination of environmental and genetic factors, and while some environmental risk factors have been identified, such as smoking and exposure to pesticides, studying the environment remains complex ^5^. For this reason, within this study we have chosen to focus on genetic risk factors in the context of PD. Interestingly, several genetic risk factors for PD identified by GWAS also influence the overall risk in carriers of *LRRK2* or *GBA1* mutations ^6,7^, which are the most common genetic causes of PD. In addition, multiple GWAS-nominated loci include genes implicated in monogenic forms of PD (e.g. *SNCA* and *LRRK2*), highlighting a clear etiologic link between monogenic and sporadic disease. Thus, understanding the molecular mechanisms underlying known genetic risk factors and mutations would provide actionable insights into the biology of disease risk, onset, progression, and modifiers of disease.

While the pace of genetic discovery has increased dramatically in recent years, our ability to characterise the associated function and dysfunction of nominated genes and risk loci has not matched this progress. Research centred on the biology of genes that contain rare disease-causing mutations has revealed important insights into the molecular mechanisms leading to disease; however, it is challenging to demonstrate how risk loci identified by GWAS may lead to disease. This is largely due to the complexity of these risk signals and the lack of large-scale reference data to interpret the molecular outcomes at these risk loci. A significant issue arises when unravelling GWAS loci due to the complex architecture of the human genome, meaning that modifier and risk loci identified by GWAS generally nominate genomic regions and not specific genes. Adding to this complexity, disease effect sizes are modest, the cellular context is often unknown, and the genetic mediator is generally unlikely to be protein coding. Extensive experimental work has provided clear insights into the molecular consequences of these variants but has not yet shown the influence of additional risk factors on these molecular disturbances, which is essential to understand why some carriers of these risk factors develop disease and others do not.

Studying the biology of GWAS loci in traditional cellular and animal models is extremely challenging due to large linkage disequilibrium (LD) blocks resulting in many highly correlated variants. Additionally, variants identified by GWAS are generally non-coding and correlating these variants to a causative gene is difficult. Low effect sizes and uncertainty in the resulting phenotype further confounds the identification of adequate models further. Therefore, the large number of known and to-be-discovered risk loci require an alternative strategy to understand underlying biology. The development of human induced pluripotent stem cell (iPSC)-based cellular models provides a unique opportunity to address the collective impact of genetic risk factors and define the relevant cellular context for modelling these variants at scale. It is important to note that iPSC models are unlikely to be able to model fulminant disease processes that likely take decades to develop in the context of organismal ageing. However, they may still be useful in identifying proximate molecular signatures that can be captured in cells containing specific risk factors or mutations. The collection of Parkinson’s Progression Markers Initiative ^8^ (PPMI, https://www.ppmi-info.org/) iPSC lines carrying different mutations and combinations of genetic risk factors allows research into the molecular consequences of the burden of genetic risk factors in a single patient. While the PPMI iPSC resource is not yet large enough to investigate all possible combinations of genetic risk and modifying factors, it can shed light on the molecular consequences caused by different combinations of the major genetic risk factors in PD.Molecular, cellular and genomic methods that can quantify epigenetic, regulatory, transcriptomic, proteomic and cellular alterations have the potential to provide us with an atlas that describes coordinated molecular and cellular changes. When such maps are generated in cells from varied genetic backgrounds, they can reveal the consequences of genetic variation on complex processes and how these consequences are interrelated. Combining iPSC approaches with quantitative molecular assays provides the capacity to assess genes of interest and risk loci at scale within a disease-relevant cellular context and an unprecedented opportunity for insights into the pathogenesis of PD.

In order to create this atlas we formed the FOUNdational Data INitiative for Parkinson’s Disease (FOUNDIN-PD; https://www.foundinpd.org/). Here we focused on the production of a large series of iPSC lines, driven to a dopaminergic neuronal cell type using standardized methods, from which a host of genetic, epigenetic, regulatory, transcriptomic, and cellular data were collected (Figure 1, blue icons represent assays included in the initial data release and light blue icons represent ongoing assays and will be released at a later stage). All iPSC lines are derived from subjects within PPMI. We describe here the production and characterization of the first release of the FOUNDIN-PD data. We recognize that while this first phase is larger than any other systematic iPSC study performed to date in PD, it represents only a pilot. This phase of data will most immediately be useful in examining high risk effects. As a part of this resource, we have also created a portal for data access and analysis and provide evidence that this system represents a relevant cellular context to investigate PD-related risk alleles. This represents a large multi-omic iPSC-derived DA neuron dataset which will serve the community as a unique resource. Lastly, we discuss the opportunities and challenges that these data have revealed for the next stages of FOUNDIN-PD.

**Figure 1.**
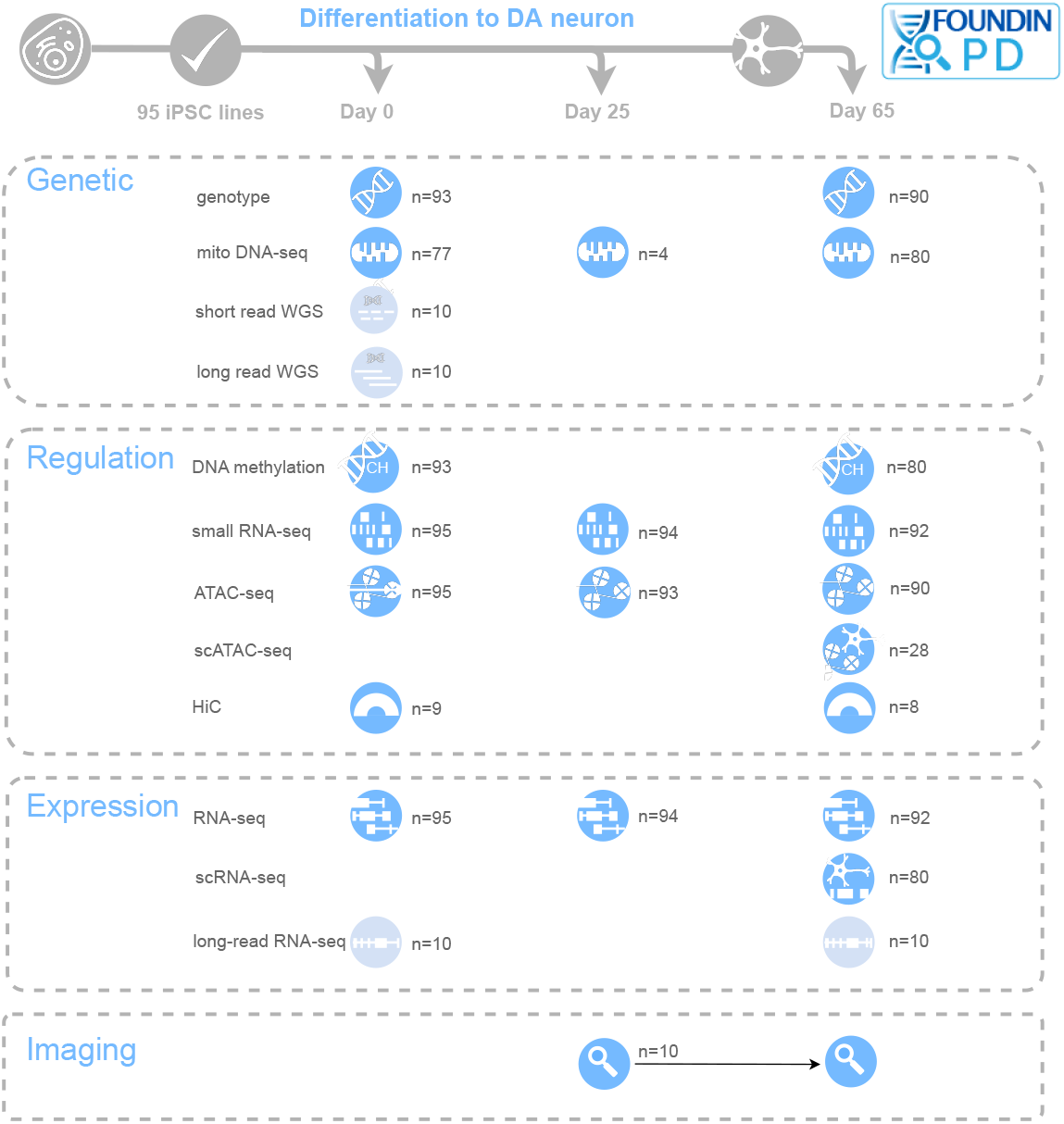
Graphical overview of the FOUNdational Data INitiative for Parkinson’s Disease (FOUNDIN-PD). Classes of assays, timepoints and number (n) of samples included in each assay are shown. Blue icons represent assays that are included in the initial data release and light blue icons represent assays that are ongoing and will be released at a later stage.

## Results

### FOUNDIN-PD overview

The basis of FOUNDIN-PD is the generation of molecular readouts from 95 iPSC lines driven to a DA neuronal state using consistent methods for all lines (Figure 1, Supplemental Table 1). These lines were available as a part of PPMI, a landmark longitudinal study that has collected data from more than 1,400 individuals at 33 sites in 11 countries and contains a wealth of clinical, imaging, and biomarker data (https://www.ppmi-info.org/). From the PPMI iPSC collection, we included lines derived from healthy controls (HC), idiopathic PD (iPD) patients, and individuals carrying known disease-linked mutations (Monogenic).

Genome sequence data were available for all donors, thus we were able to not only identify subjects with damaging mutations in *LRRK2* (p.G2019S, n=25 and p.R1441G, n=1), *GBA1* (p.N370S, n=20), and *SNCA* (p.A53T, n=4) (hereafter refer to these variants as *LRRK2*+, *GBA1*+, and *SNCA*+, respectively) (Table 1), but also those with high and low polygenic risk scores (Supplemental Figure 1, Supplemental Table 1). These 95 iPSC lines were differentiated into DA neurons using a well-established protocol ^9^, with minor modifications (Figure 2A; protocols.io, dx.doi.org/10.17504/protocols.io.bfpzjmp6)^10^, and an automated robotic cell culture system ^11^. The differentiation protocol was previously established and validated with five in-house lines where three independent differentiations produced neuron-enriched cultures with averages of 90% TUJ1 (range: 73-99%) and 80% *MAP2* (range: 72-98%) positive cells identified by immunocytochemistry (ICC). More than 60% (range: 56-89%) of the differentiated cells were also positive for the DA marker tyrosine hydroxylase (*TH*) (Supplemental Figure 3). This was considered a satisfactory differentiation efficiency and, therefore, the same protocol was applied to differentiate the 95 PPMI iPSC lines.

**Figure 2.**
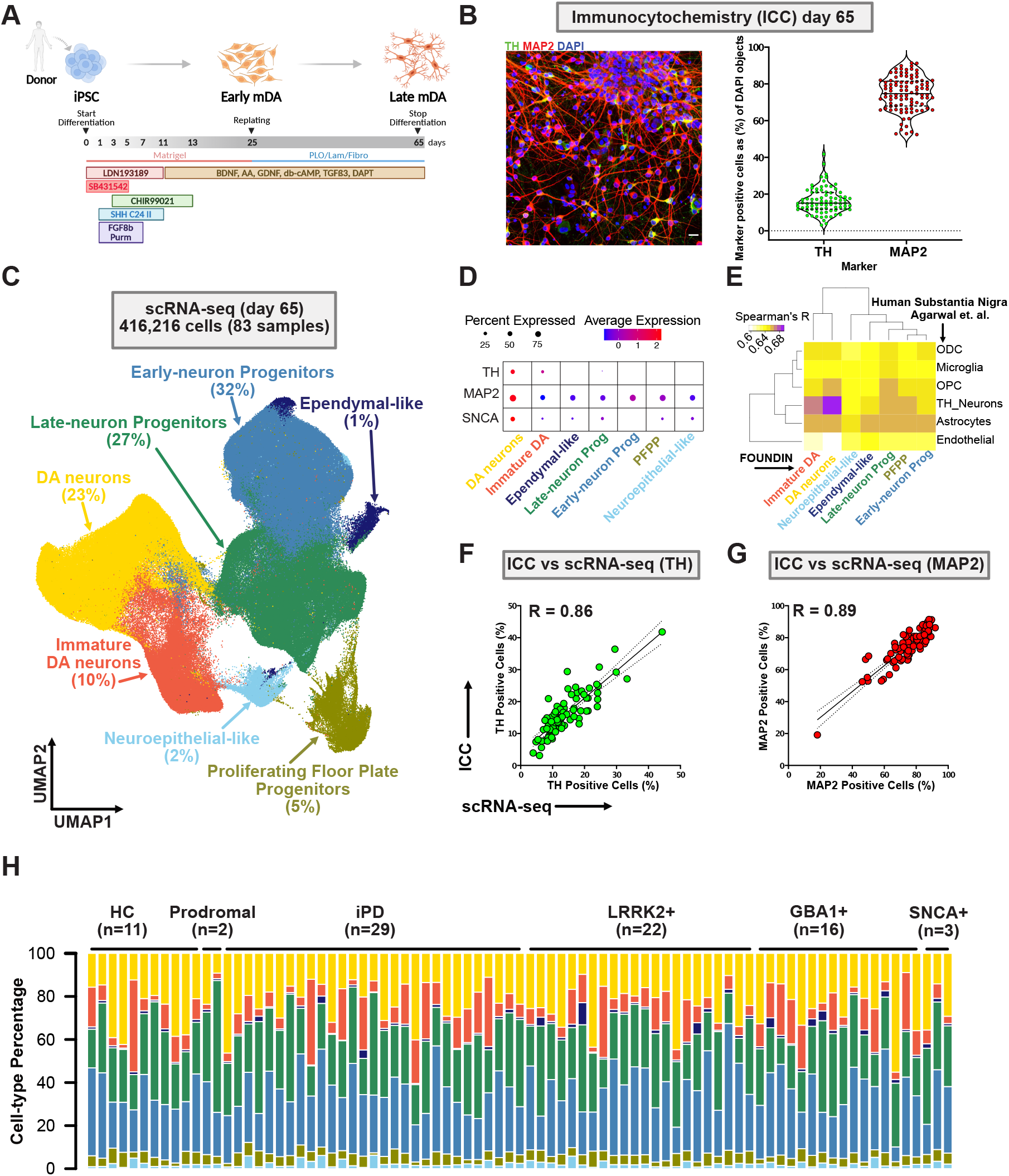
Quality control and scRNA-seq on day 65. A, Schematic overview of the differentiation protocol to dopaminergic neurons. B, Left: Representative ICC image showing *TH* (DA neurons) and *MAP2* (neuron) positive cells co-stained with DAPI (nuclei). Scale bar: 50 µm; Right: Percentage of *TH* (DA neuron) and *MAP2* positive cells detected by ICC and normalized to the total number of nuclei. Data is represented as the percentage of positive cells per 30 imaged fields. Each dot represents one cell line (n=95). C, Uniform Manifold Approximation and Projection (UMAP) illustrates cell clusters identified at day 65 (n=416,216 single cells, n=79+4 control replicate cell lines). Cell types with their respective percentages are indicated. D, Percentage of cells and average expression level of *TH, MAP2* and *SNCA* for each cell type. The dot color scale from blue to red, corresponds to lower and higher expression, respectively. The size of the dot is directly proportional to the percentage of cells expressing the gene in a given cell type. PFPP, Proliferating floor plate progenitors; Prog, Progenitors) E, Spearman’s correlation test showing high correlation of gene expression across FOUNDIN-PD DA neuronal types and post mortem substantia nigra human brain. ODCs, oligodendrocytes; OPCs, oligodendrocyte precursor cells. See Supplemental Figure 6A for UMAP of cell types identified by using Agarwal and collaborators data. F-G, Correlation between percentages of *TH* (Pel-Freez) and *MAP2* positive cells in ICC and scRNA-seq (R, Pearson correlation coefficient; p<0.0001). Each dot represents one cell line (n=83). H, Cell type percentage by cell line showing variability in differentiation efficiency across the iPSC lines. Each color represents the cell types annotated in scRNA-seq UMAP and each bar represents a different cell line. In total, 83 cell lines were included in the scRNA-seq. HC, Healthy Control (n=8 plus 3 replicates of the control line); Prodromal (n=2); idiopathic PD (iPD;n=29), monogenic PD (*LRRK2*+, *GBA1*+ or *SNCA*+; n=41). Colors refer back to clusters in panel C: yellow = DA neurons, Orange = Immature DA neurons, light blue = Neuroepithelial-like cells, olive = PFPP, green = Late-neuron progenitors, blue = Early-neuron progenitors, indigo = ependymal-like cells.

PPMI lines were differentiated in five batches (ranging from 10-30 cell lines per batch) until day 25 or 65, followed by harvesting the cells for ICC and molecular assays. Quantification of *MAP2*- and *TH*-positive cells revealed that on average 80% (range 52-93%) of the cells were converted to neurons, and 20% of the cells (range 4-42%) expressed *TH* (Figure 2B; Supplemental Figure 2), showing a higher variability in differentiation efficiency than the in-house iPSC lines used in protocol optimization. The average proportion of *TH*-positive cells in the iPSC lines, relative to all cells in the culture, was similar when assessed by ICC with two independent *TH* antibodies and the estimate of the proportion of *MAP2* positive cells, relative to all cells, was also independent of the *TH* antibody used (Supplemental Figure 4A). To measure how robust and reproducible the differentiation protocol was using our automated system, we included a control line in each batch as a technical replicate (n=5). The percentage of *MAP2*- and *TH*-positive cells obtained from the control cell line using ICC across all five differentiation batches was consistent (Supplemental Figure 4B), and no significant differences in the percentage of neurons or *MAP2*- and *TH*-positive neurons between batches were identified (p>0.2 for both).

### Quantifying gene expression in FOUNDIN-PD data using RNA sequencing

To further characterise the iPSC-derived neurons, we generated a wealth of data types including genetic, epigenetic, regulatory, transcriptomic, and cellular imaging data (Figure 1). To fully characterise the identity of the cell types generated by the iPSC differentiation protocol used in the present study, we performed single-cell RNA sequencing (scRNA-seq) on the majority of the day 65 cell lines (n=79 with 4 control replicates, 84% of total included iPSC). In total, 416,216 high quality cells were retained, with an average of 5,015 cells per sample ^12^(range 584 to 9,640). Cells were first clustered using an unsupervised method (Louvain algorithm), and then annotated based on canonical cell type markers found in the differentially expressed genes of the cluster (Supplemental Table 2, Supplemental Figure 5). Seven distinct broad-cell types were identified across all samples and are defined as: early neuron progenitors expressing RFX4, HES1 and SLIT2 ^12^ (131,251 cells, 32% of total), late neuron progenitors expressing DLK1, LGALS1 and VCAN ^13^ (113,425 total, 27% of total), DA neurons expressing *TH*, ATP1A3, ZCCHC12, *MAP2*, SYT1 and SNAP25 ^14^ (96,623 total, 23% of total), immature DA neurons expressing TPH1, SLC18A1, SLC18A2 and SNAP25 ^12^ (41,267 total, 10% of total), proliferating floor plate progenitors (PFPP) expressing HMGB2, TOP2A and MKI67 ^13,15^ (18,984 total, 5% of total), neuroepithelial-like cells expressing KRT19, KRT8 and COL17A1 ^16^ (8,979 total, 2% of total) and ependymal-like cells expressing MLF1, STOML3 and FOXJ1 ^15^ (5,687 total, 1% of total) (Figure 2C). Overall, expression of *TH, MAP2* and *SNCA* was clearly enriched in the neuronal cell types (Figure 2D).

Next, we assessed how similar the expression signatures are of the cultured DA neurons vs human tissue DA neurons and also how our cultured DA neurons compare to previously published DA neuron datasets. We compared our identified cell type populations to public datasets from human post mortem substantia nigra ^14^ and human iPSC-derived DA neuron subtypes using a slightly modified DA neuron differentiation protocol and distinct set of iPSC cell lines ^12^. The DA neuron population identified in our data showed the highest correlation (Spearman’s R=0.69) with the *TH*-positive neuron cluster found in human post mortem substantia nigra (Figure 2E). This correlation was also identified using dendrogram clustering of DA neurons from this study and the *TH*-positive neurons from human post mortem substantia nigra (Supplemental Figure 6A-B). The second highest correlation was observed between our immature DA neurons and the *TH*-positive neuronal cluster from Agarwal et al (Spearman’s R=0.67, Supplemental Figure 6A-B). Additionally, both immature and DA neurons were highly correlated with the iPSC-derived DA neuron subtypes (DAn1-4) identified by Fernandes and collaborators ^12^ (Supplemental Figure 6C), which were produced using a similar iPSC to DA neuron differentiation protocol. Another similarity detected between both iPSC-derived neuron datasets was the expression of serotoninergic markers in our immature DA neurons (FOUNDIN-PD, Supplemental Table 2) and the previously published DAn2 ^12^.

To validate the identified (neuronal) cell types using scRNA we compared ICC-based estimates of DA neurons (*TH*-positive cells) and overall neurons (*MAP2*-positive cells) with the percentage of positive cells obtained from scRNA-seq data showed high correlations (Pearson correlation of R=0.8562, P<0.0001 and R=0.8916, P<0.0001 for *TH* (Pel-Freeze) and *MAP2*, respectively (Figure 2F-G). Similar results were obtained with a second *TH* (Millipore) antibody (Supplemental Figure 7). Although the differentiation efficiency (percentage of each cell type) varied between cell lines (Figure 2H), no consistent cell type enrichment could be identified based on batch, phenotype, recruitment category, genetic sex or PD-linked genotype (*GBA1*+, *LRRK2*+, *SNCA*+) (Supplemental Figure 8). Additionally, a very high correlation was observed (R>0.9) between technical replicates (n=4) using gene-level scRNA-seq data of the identified DA neuron cluster (Supplemental Figure 9) and total *TH* and *MAP2* levels (Supplemental Figure 4), suggesting that, while there is variability in differentiation efficiency across the lines, this is likely not caused by the robustness of the differentiation protocol but may be due to inherent characteristics of each individual line.

To assess gene expression differences across multiple time-points during differentiation we generated bulk RNA-seq at day 0 (n=99), day 25 (n=98) and day 65 (n=96) with each timepoint including five technical replicates of the control line. A principal component analysis (PCA) of bulk RNA-seq showed clear clustering by timepoint (Figure 3A). In accordance with the scRNA-seq data, we also observed a very high correlation (R>0.9) between the technical replicates in the gene-level expression of each timepoint in bulk RNA-seq data (Supplemental Figure 10A-C). Further evaluation of the bulk RNA-seq data across all timepoints showed clear transcriptional enrichment signatures that correlate with neuron-like features, including synapse assembly, neurotransmitter transport and action potential (Supplemental Table 4) at timepoints day 25 and day 65. Additionally, specific genes of interest, such as *MAP2* and *TH* (Figure 3B-C), and *GBA1, SNCA, LRRK2* and SYN1 (Supplemental Figure 11A-D) showed increased expression levels as cells were differentiated. Concurrently, iPSC-associated genes such as POU5F1 (Figure 3D), NANOG and TDGF1 (Supplemental Figure 11E-F) showed significantly reduced expression at later timepoints relative to day 0, which correlated with a decrease in pathway signatures of iPSC differentiation and growth, including somatic stem cell population maintenance and positive regulation of cell population proliferation (Supplemental Table 5).

**Figure 3.**
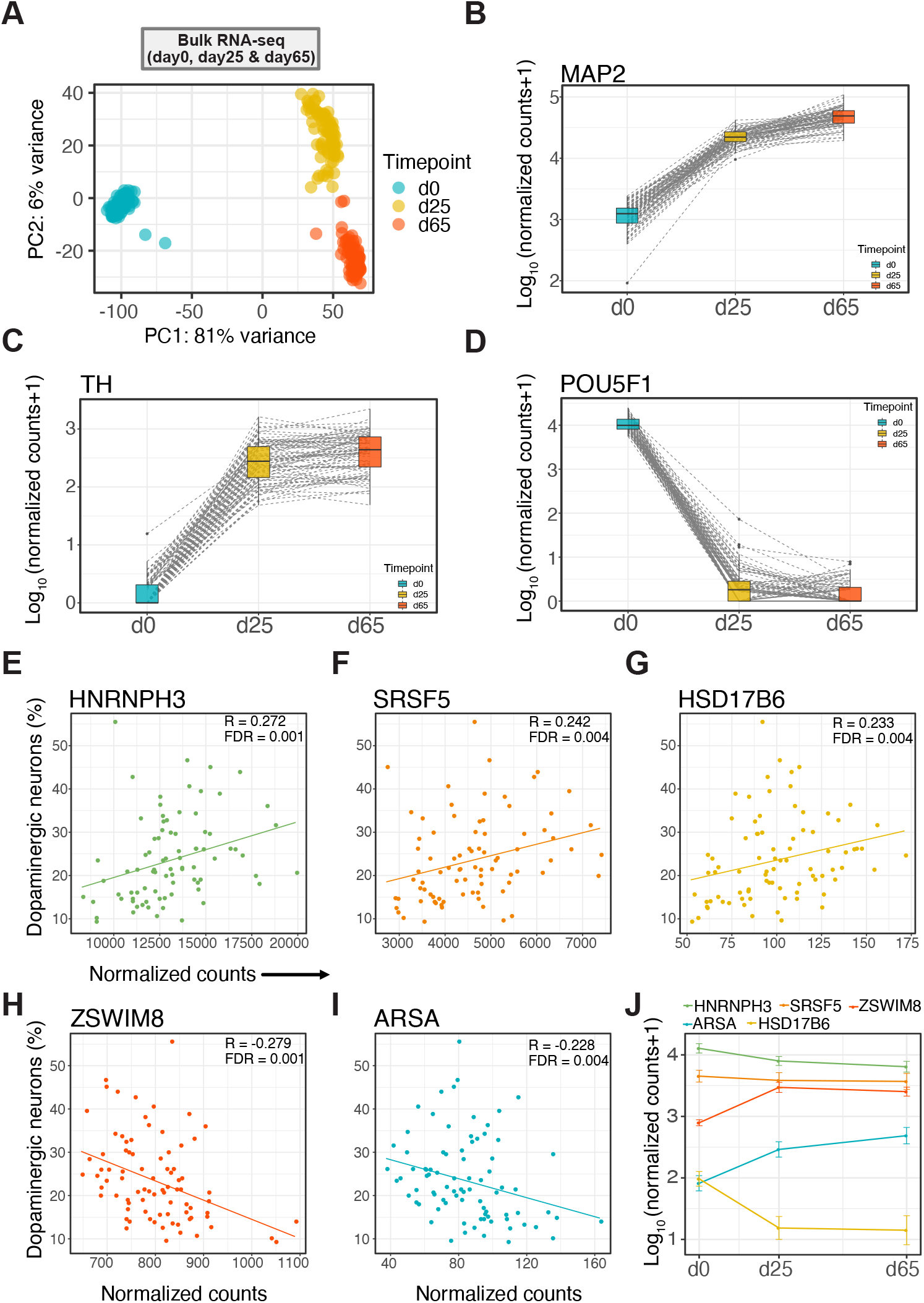
Bulk RNA-seq and neuronal differentiation efficiency prediction. A, Principal component analysis (PCA) of bulk RNA-seq showing clustering by timepoint (days 0, 25 and 65). B-D, Changes in expression of neuronal (*MAP2*), dopaminergic (*TH*) and iPSC (POU5F1) genes from day 0 to day 65. E-I, Genes significantly correlated with neuronal differentiation efficiency. *HNRNPH3, SRSF5* and *HSD17B6* show positive and ZSWIM8 and ARSA negative correlation. J, Expression levels of genes associated with neuronal differentiation efficiency. All five genes are significantly differentially expressed between day 0 and day 65 (adjusted p-value < 0.05).

Next, we used the day 0 bulk RNA-seq gene expression data to predict DA neuronal differentiation efficiency. We defined DA neuronal differentiation efficiency as the fraction of DA neurons in our scRNA-seq datasets at day 65 using a similar method described by Jerber and collaborators ^15^. Using logistic regression, ten genes were identified that have a non-zero coefficient and predicted good neuronal differentiation efficiency with an area under the curve (AUC) = 0.93 and 0.87 accuracy (95% confidence interval [0.78, 0.93]) (Supplemental Figure 12A-C). Repeated 5-fold cross-validation achieved a mean AUC of 0.85 with 0.03 standard deviation. Out of these ten genes that have a non-zero coefficient, five were significantly correlated with neuronal differentiation efficiency (FDR < 1%, Figure 3E-I and Supplemental Figure 12D). Three (*HNRNPH3, SRSF5* and *HSD17B6*) of these associated genes were positively correlated with neuronal differentiation efficiency. Moreover, the expression of these genes was significantly reduced as iPSC were differentiated to DA neurons (adjusted p-value < 0.05 from day 0 to day 65, Figure 3J), suggesting that their high expression in iPSCs may represent an increased differentiation potential. Previous studies have shown *SRSF5* is associated with neuronal differentiation efficiency (R=0.25, adjusted p-value of 0.013 ^15^ and that *SRSF5* binds to pluripotency-specific transcripts and positively correlates with the cytoplasmic mRNA levels of pluripotency-specific factors ^17^. Interestingly, *HNRNPH3* is also a known RNA binding protein, suggesting that regulation of RNA binding may be an important pathway for neuronal differentiation. The remaining two associated genes (ZSWIM8 and ARSA) were negatively correlated with neuronal differentiation efficiency and their overall expression was significantly increased during differentiation (adjusted p-value < 0.05 from day 0 to day 65, Figure 3J).

### Establishing regulatory maps of iPSC-derived dopaminergic (DA) neurons

To identify epigenetic and regulatory features of genes in iPSC and differentiated DA neurons, we generated DNA methylation, ATAC-seq (both bulk and single-cell), HiC sequencing and small RNA-seq data across several timepoints. DNA methylation data from bulk cultures were generated at day 0 (n=97 after QC, including five technical replicates) and day 65 (n=82 after QC, including three technical replicates). These data were generated to assess changes in epigenetic patterns that potentially regulate gene transcription. The methylation data showed clear separation between both timepoints (Supplemental Figure 13). Additionally, marker genes such as *MAP2* and *TH* showed a significant reduction in methylation from iPSC at day 0 to DA neurons at day 65 (Supplemental Figure 14A-E). ATAC-seq (Assay for Transposase-Accessible Chromatin using sequencing) is a commonly used technique to assess genome-wide chromatin accessibility. Bulk ATAC-seq was generated from cultures at day 0 (n=99), day 25 (n=97) and day 65 (n=94), with each timepoint including the control line with five technical replicates. As with the other assays, PCA across all samples showed clustering of samples by timepoint (Figure 4A). Peak sets merged from all samples at each timepoint showed an enrichment in open chromatin near promoters (0-3000 bp from the transcription start site) and a corresponding reduction in the proportion of peaks in distal intergenic regions by analysis with Cistrome ^18^ (Supplemental Figure 15A). Interestingly, we observed an increase in evolutionary sequence conservation at merged peak sets in more differentiated cells, where the lowest PhastCons score ^19^ across all peak sets was at day 0 and the highest at day 65 (Supplemental Figure 15B).

**Figure 4.**
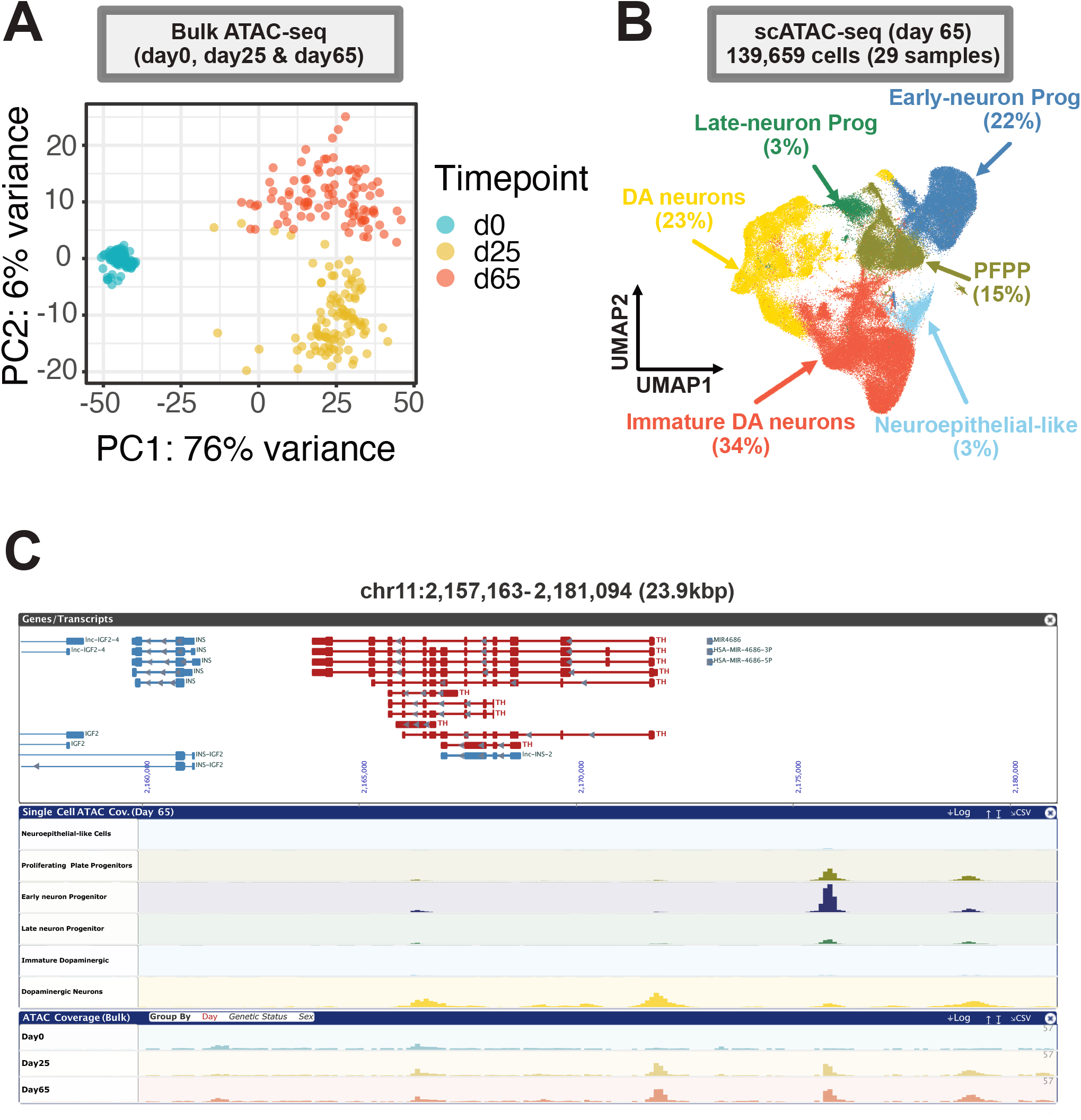
Chromatin accessibility in iPSC-derived neurons on day 65. A, Principal component analysis (PCA) across all bulk ATAC-seq samples showing clustering by timepoint. B, Uniform Manifold Approximation and Projection (UMAP) of scATAC-seq data at day 65 showing the clustering of 139,659 cells (from 29 samples) and similar broad cell types as in scRNA-seq (Figure 2C). C, chromatin accessibility data at the *TH* locus showing timepoint specific peaks identified in bulk ATAC-seq at day 25 and day 65 and cell type-specific peaks in scATAC-seq at day 65. This figure was generated using the FOUNDIN-PD browser (https://www.foundinpd.org).

To provide a cell-type-specific view of chromatin accessibility in our differentiated cells, we generated single cell ATAC-seq (scATAC-seq) at day 65 for a subset of the samples (n=27+2 replicates). Following quality control, 139,659 cells were retained, with an average of 4,816 cells per sample (range 944 to 11,649). We identified similar broad cell types, as in the scRNA-seq data (Figure 4B). However, the percentage of immature DA neurons and progenitor cell types was different between the scRNAseq and scATAC-seq datasets. Cell-type-specific chromatin accessibility was observed at particular genes of interest. For example, a distinct peak was identified at the promoter of *TH* in bulk ATAC-seq at days 25 and 65 that, when examined in scATAC-seq, only appeared in the DA neuron cluster (Figure 4C). Overall, ATAC-seq reads were enriched at the promoters of expressed genes, but it is important to note that not all genes in this region had peaks at their promoters in either bulk or scATAC-seq, reflecting cell type-specificity. ATAC-seq is also known to identify cell type-specific intergenic regulatory regions. Reflecting this, we observed peaks at putative regulatory regions upstream of *TH* that were restricted to the progenitor and DA neuron populations, suggesting that these sequences may play a role in priming *TH* expression. A peak identified at the promoter of *MAP2* in bulk ATAC-seq at days 25 and 65 also appeared as a broader neuronal marker in all cell types identified in scATAC-seq, except for the neuroepithelial-like cells, which are a non-neuronal cell type (Supplemental Figure 16).

HiC sequencing (HiC-seq) is a method used to identify three-dimensional chromosome interactions (chromosome loops). These loops are known to be involved in regulating gene transcription by enabling physical interactions of enhancers with their cognate promoters ^2021^. These data can be particularly useful for linking distal risk loci/variants with regulatory regions and genes. HiC-seq data was generated for a subset of batch 1 at day 0 (n=9) and day 65 (n=8) due to the large number of cells required as input for this assay. The HiC chromosome loops showed clear separation of both timepoints, and marker gene *MAP2* showed distinct differences in HiC loop patterns (Supplemental Figure 17 and 18).

To complement the other gene expression and regulatory data, we performed small RNA-seq to investigate various classes of small RNAs, including microRNAs (miRNAs, piRNAs, tRNA fragments) and other noncoding RNAs less than 50 basepairs (bp), which are often involved in gene silencing and post-transcriptional regulation of gene expression. Small RNA-seq was generated at day 0 (n=99), day 25 (n=98) and day 65 (n=96), with each timepoint including the shared line with five technical replicates. Separation was seen between all timepoints (Supplemental Figure 19). The miRNAs that were significantly upregulated between day 0 and day 65 are enriched for those that are highly expressed in tissues from the central nervous system when examined across 34 different tissues ^22^ (Supplemental Figure 20).

### Longitudinal imaging of iPSC-derived dopaminergic (DA) neurons

To assess the relationship between the various molecular readouts described above and neuronal phenotypes, we performed longitudinal imaging and single cell analysis. Cell-based imaging can be a valuable complementary approach to molecular analyses for characterising phenotypes. To perform longitudinal single cell analysis, 10 out of 95 iPSC lines differentiated into DA neurons (batch 1) were frozen on day 25 of differentiation. Frozen neurons were thawed, plated in 96-well dishes, matured for an additional 25 days and transduced with a lentivirus to express GFP under the control of a synapsin I (SYN1) promoter on day 50. To focus our analysis on the subpopulations of cells perceived to be most relevant to PD, we expressed GFP from a SYN1 promoter to restrict marker gene expression to relatively mature neurons. Fluorescence became visible within a day of transfection, and robotic microscopy ^23^ was used to image cells every 24 hours for approximately 10 days. Cells exhibiting GFP fluorescence had the characteristic morphological features of relatively mature DA neurons (Figure 5A). The GFP morphology signal was used to unambiguously identify individual neurons and to track each cell from one imaging timepoint until the next. Because of its ability to track individual cells, robotic microscopy can monitor whether and how phenotypes change over time and obtain a cumulative measure of phenotypic endpoints that better controls for variability and increases sensitivity of comparisons of phenotypes between cohorts. Live neurons could be followed throughout the duration of the experiment. Representative neuron survival over 6 days is shown in Figure 5A. Cell death was detected as an abrupt loss of signal, indicative of a loss of membrane integrity (Figure 5A). In total, 2,992 cells were analysed across the 10 lines. The time required for complete loss of signal (time of death) from hundreds of neurons was analysed with the Kaplan-Meier survival model ^24^, and cumulative risk of death curves were generated (Figure 5B). Comparison of lines from individuals harboring *GBA1* mutations compared to healthy control lines demonstrates a significant increased risk of death in *GBA1* lines (Fig. 5C).

**Figure 5.**
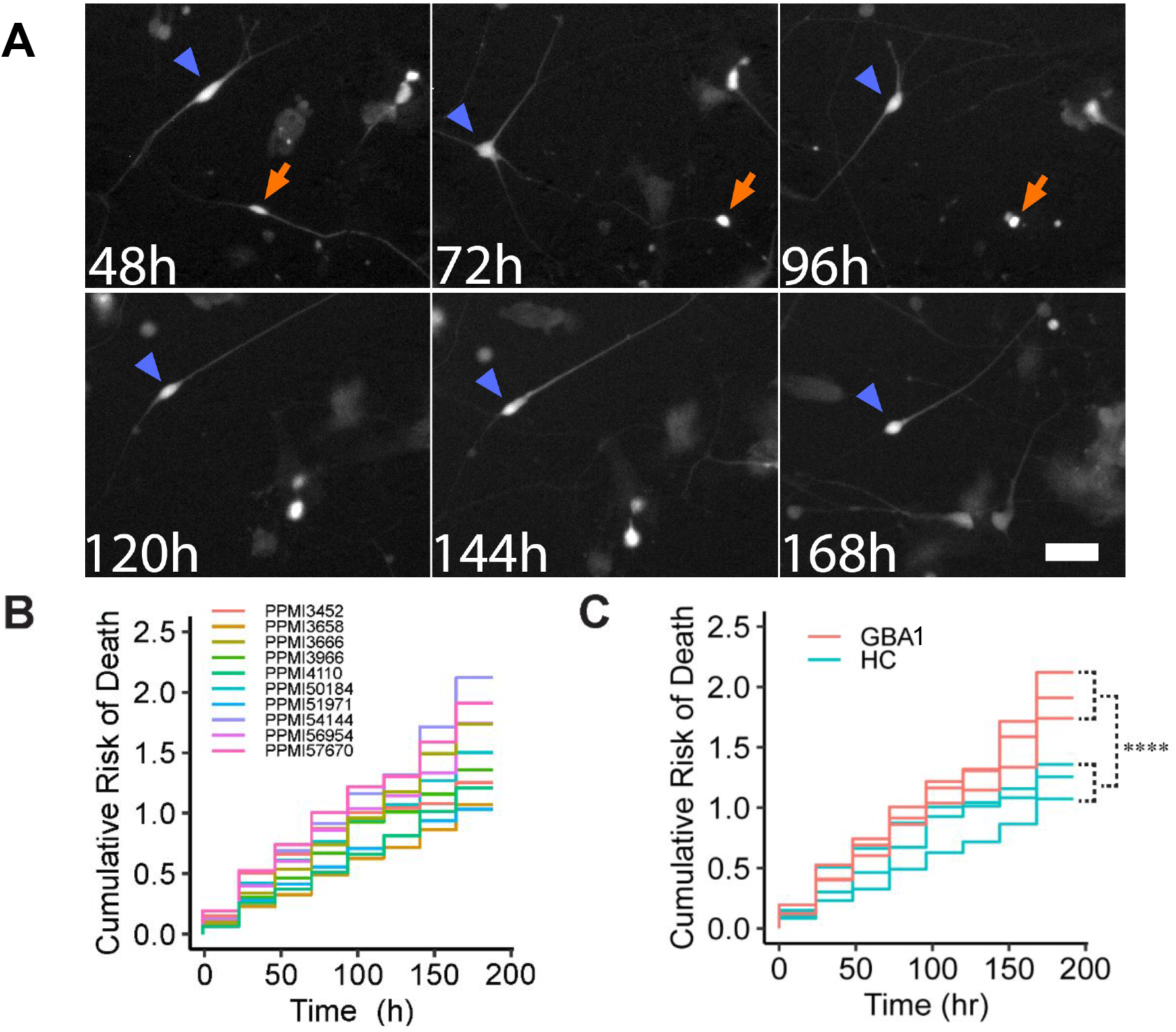
Automated longitudinal imaging of dopaminergic neurons. A, Time-lapse imaging of dopaminergic neurons (PPMI4110) expressing synapsin-I driven GFP. Analysis started on day 55-56 of differentiation. One neuron (green arrowhead) survives the entire duration of imaging. A second neuron (red arrow) dies at 96h. Scale bar: 60 µm. B, Cumulative risk of death curves showing the neuronal survival from all batch 1 lines over 8 days of automated imaging. C, Cumulative risk of death curves show increased degeneration in dopaminergic neurons differentiated from *GBA1* PD lines compared with healthy control lines over 8 days of automated imaging (****p < 0.0001; based on 891 neurons from *GBA1* lines and 647 neurons from healthy control volunteers (HC)).

### Testing the contextual fit of iPSC-derived DA neurons for modelling PD related genetic risk

We identified a wide genetic risk spectrum across the iPSC lines that we studied (Table 1, Supplemental Figure 1). In addition to the contribution of genetic risk from known damaging variants in *GBA1, LRRK2* and *SNCA*, there is a substantial common risk element that can be quantified by polygenic risk score, as previously shown using GWAS ^2^. One limitation of GWAS is that it often cannot identify the causal variants, genes or relevant cell type for each locus without additional gene expression or functional data. A method commonly used to infer cell-type-relevance based on GWAS statistics is MAGMA (Multi-marker Analysis of GenoMic Annotation). This method relies on the convergence of unbiased genetic risk maps with single cell expression data; the enrichment of expression of genes from risk loci in individual cell types acts as a powerful indicator of cell type relevance ^25^. Previous analysis using mouse and human brain expression data has shown that DA neurons are a critical cellular context for PD-related genetic risk ^2,4^. Analysis of the scRNA-seq expression data from this study revealed a dramatic enrichment of expression of genes within PD-linked risk loci in the two identified DA cell types (immature and DA neurons), relative to the other cell types (Figure 6A and Supplemental Table 6). Combined with the comparisons detailed above, these data reveal that this model resembles human brain neurons and provides a cellular context that is appropriate for modelling complex genetic risk in PD.

**Figure 6.**
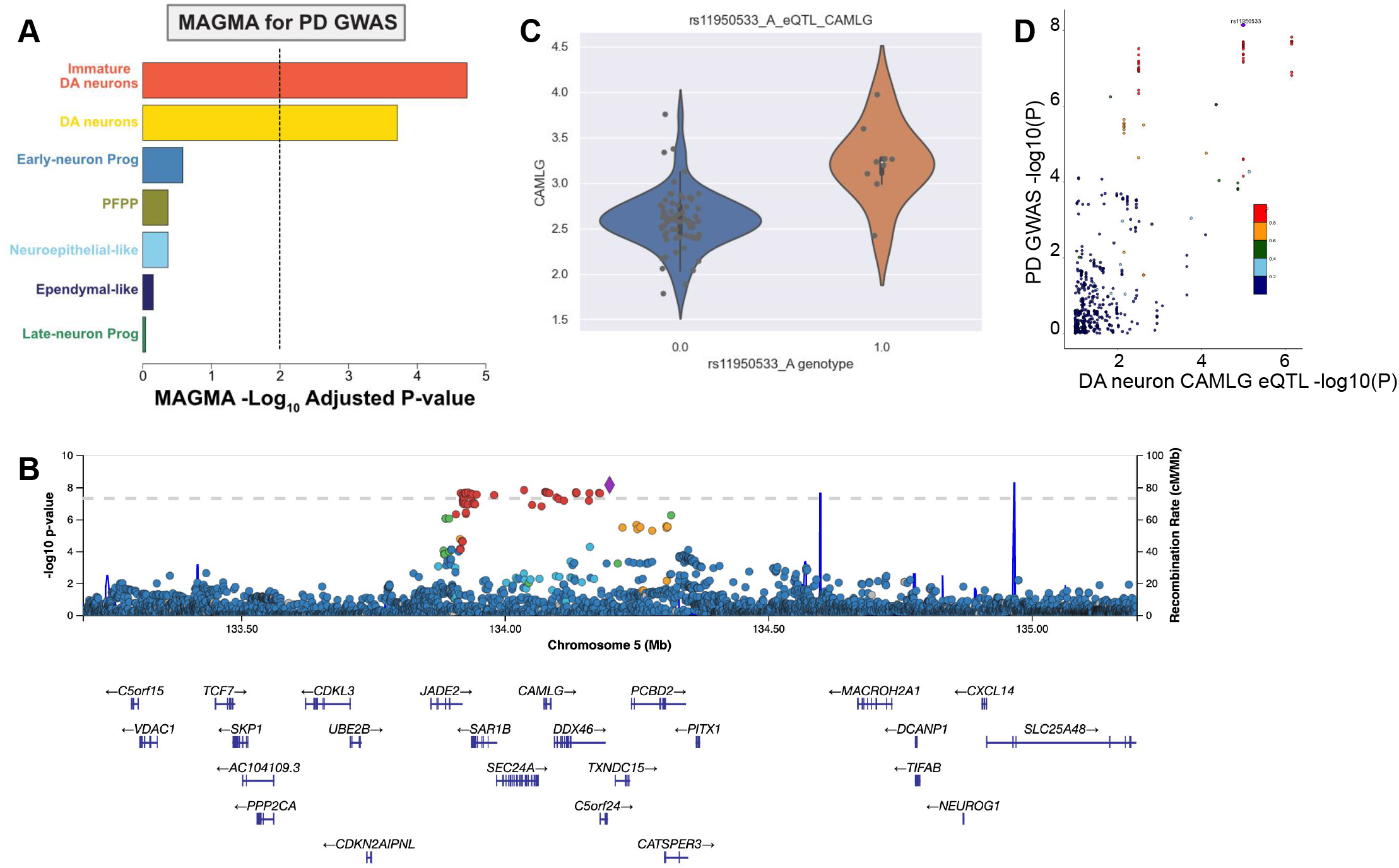
Using scRNAseq expression data to dissect genetic risk. A, Multi-marker Analysis of GenoMic Annotation (MAGMA) gene-set enrichment based on the scRNA-seq data showed significant associations with both dopaminergic neuron cell clusters. Colors represent the same cell types as in Figure 2C. B, LocusZoom plot of locus 28 with rs11950533 as the index variant. Association data is derived from the most recent PD GWAS (Nalls et al. 2019). C, Violin plot showing correlation between the genotype at rs11950533 and expression of *CAMLG* in the DA neuron cell cluster. D, LocusCompare plot of the correlation between the PD GWAS (Nalls et al. 2019) association results and the scRNA-seq expression quantitative trait (eQTL) analysis.

In an effort to nominate potential causal genes and molecular mechanisms tagged by each GWAS locus, we combined whole genome sequencing with our scRNA-seq data in differentiated cells to identify expression quantitative trait loci (eQTL) in each broadly defined cell type. Using this approach, we replicated known eQTL in the KANSL1 and LRRC37A region reflecting the H1/H2 MAPT haplotypes (Supplemental Figure 21A-D). When exploring the eQTL results further, we specifically focused on the 90 risk variants from the most recent GWAS in PD ^2^. Multiple variants in this dataset showed significant eQTL associations in at least one of the defined cell types in our DA neuron scRNA-seq (Supplemental Table 7). For example, the locus with rs11950533 as the lead variant harbours at least 25 genes (Figure 6B), and based on the PD GWAS browser prioritisation tool ^26^, four (*CAMLG, JADE2, TXNDC15* and *SAR1B*) were prioritised based on their high correlation between cortical brain eQTL data ^27^ and PD GWAS signal (Supplemental Figure 22A-D). In the current FOUNDIN-PD scRNA-seq expression data, an eQTL for *CAMLG* was identified (Figure 6C), which shows high correlation with the PD GWAS signal (Figure 6D). However, no eQTL signals were identified for *JADE2, SAR1B* or *TXNDC15* (Supplemental Figure 23A-C), despite all genes being expressed in our DA neurons (Supplemental Figure 24). Inspection of the *CAMLG* bulk RNA-seq eQTL signal and the PD risk signal intersection revealed that this eQTL was not detected at the iPSC state at day 0 but became detectable at day 65 (Supplemental Figure 25). This suggests that the regulatory effect signal or trajectory of *CAMLG* expression may correspond with differentiation to DA neurons. Therefore, based on FOUNDIN-PD data, *CAMLG* should be prioritised further as a candidate for functional follow-up to confirm the association between *CAMLG* and PD risk.

PD risk and scRNA eQTL signals for DA neurons also intersected with other independent PD risk loci including *TBC1D5, PRCP, CCAR2, ARIH2*, and *CCDC58*. In these genes, the PD risk variant appears to be more statistically significant associated with expression in DA neurons when compared to other cell types detected in the FOUNDIN-PD resource. This intersection of disease risk and DA neuron expression effect appears to be specific to DA neurons for *TBC1D5* (Supplemental Figure 26), *CCAR2* (Figure 7B) and *ARIH2* (Supplemental Figure 27) whereas *PRCP* and *CCDC58* (Supplemental Figures 28 and 29), and *CAMLG* (Supplemental Figure 30) showed signal that intersected PD risk signals in multiple cell types. Inspection of additional FOUNDIN-PD resources such as the single-cell ATAC peaks revealed instances of peaks that were elevated in DA neurons in comparison to other cell types and contained (Figure 7C, D) variants that were in high LD with the PD risk index variants at the locus. For *TBC1D5, CCAR2*, and *ARIH2*, the signal intersection was more specific to DA neurons, but, when inspecting the bulk RNA (Figure 7E) data, the signal was not present.

**Figure 7.**
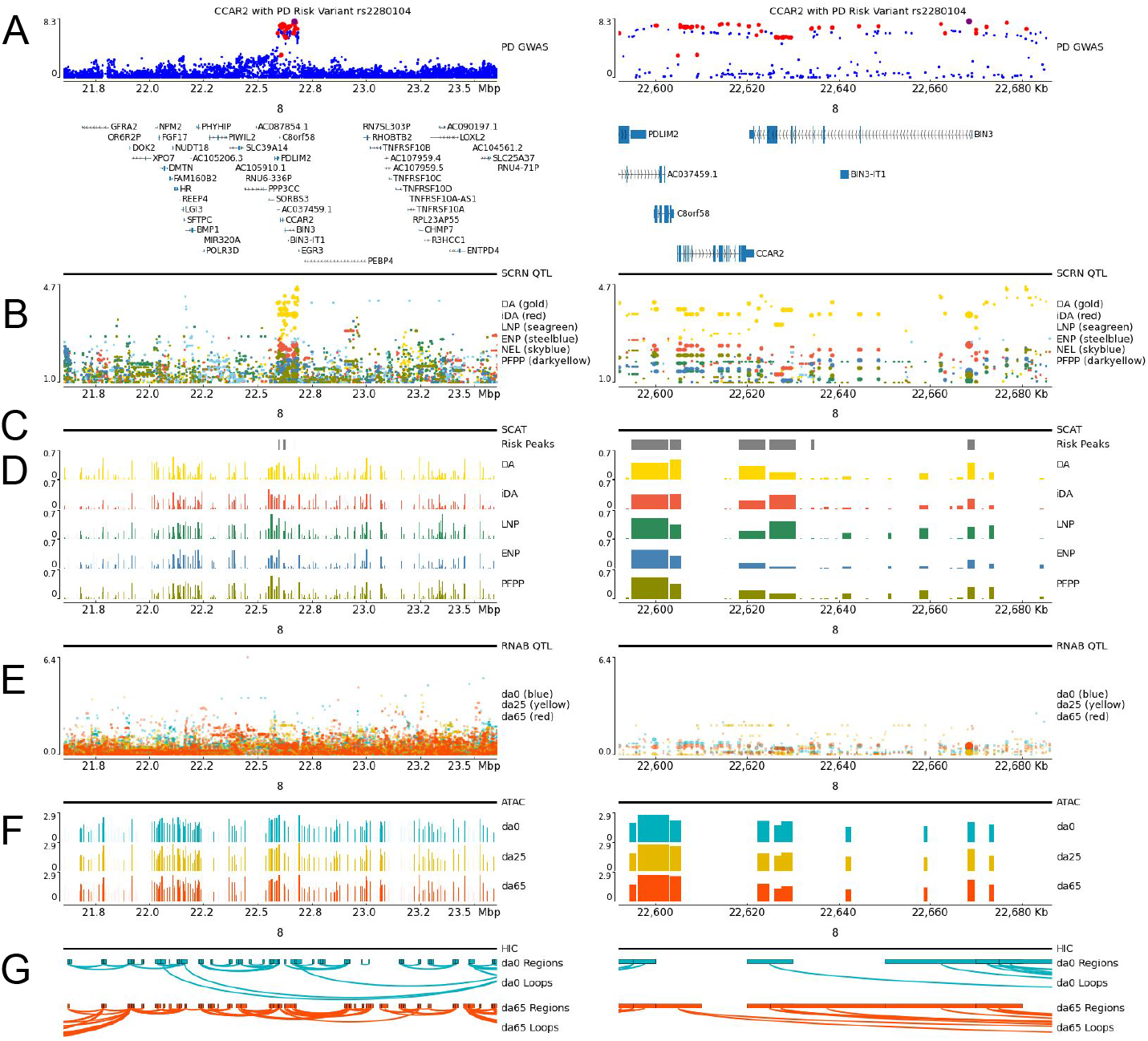
PD risk locus, FOUNDIN-PD resources, and *CCAR2* effects in dopaminergic neurons The PD risk locus near BIN3 on chromosome 8 that intersects with an eQTL for *CCAR2* in dopaminergic neurons. Tracks represent different data modalities generated or considered in FOUNDIN-PD as different data tracks; figures generated using pyGenomeTracks (Lopez-Delisle et al. 2020). The left and right panels of the figures display the same tracks where the left side is a larger region centered on the Parkinson’s disease risk locus and the ride side only includes the interval containing the index PD risk variant for this locus and variant in linkage disequilibrium with that index variant. (A) GWAS risk for PD in the region. Point size denotes r^2^ linkage disequilibrium with the PD index variant rs2280104 (large: r2=1, medium: 1 > r2 >= 0.8, small: r2 < 0.8). (B) Single-cell RNA-seq eQTL data for dopaminergic neurons (DA), immature dopaminergic neurons (iDA), late-neuron progenitors (LNP), early-neuron progenitors (ENP), neuroepithelial-like cells (NEL) and proliferating floor plate progenitors (PFPP). (C) Single-cell ATAC-seq peaks containing a variant in high linkage disequilibrium (r^2^ >= 0.8) with rs2280104. (D). Single-cell ATAC-seq peaks for different cell types. (E) Bulk RNA-seq (RNAB) eQTL results per differentiation timepoint for *CCAR2*. (F) Bulk ATAC-seq peaks separated per differentiation timepoint. (G) HiC data depicting chromatin regions connected by loops at different differentiation timepoints.

### The FOUNDIN-PD Data Portal

To allow rapid and easy data access to researchers, gene and region-level views of data are available through a web-based portal (https://www.foundinpd.org) integrating the multiple omic data types together (Figure 1). All users can access summary level data for a region (<5Mb) or gene by registering with a single sign-in Google account. A single-integrated view allows for visualisation of genomic data by genomic coordinates with tracks available for scRNA-seq, scATAC-seq, bulk RNA-seq, bulk ATAC-seq, methylation arrays, HiC-seq, small RNA-seq, among others. The portal is interactive, allowing the dynamic ability to view facets/partitions of data split by *LRRK2*/*GBA1*/*SNCA* status, sex, and diagnosis. The tracks are responsive for dynamic zooming and panning by touch or mouse, can be reordered or hidden from views. Users can download data backing the graphs via CSVs and export screen snapshots. Users who are authenticated for access to individual-level data via https://www.ppmi-info.org/, will also have the ability to visualise individual-level data. Additional phenotypic detail is available, and users can, for example, dynamically plot expression versus SNP genotype or many other variables available on subjects. The portal contains links to several additional access points, including PPMI-INFO for individual-level data and a GitHub site (https://github.com/FOUNDINPD) with analysis and standard operating procedures (SOPs). Finally, a specific single-cell view of the data is available via an embedded cellxgene instance ^28^, providing UMAP and PCA views. Through this interface, users can view identified clusters, genes, and gene families across profiled cells.

## Discussion

Genetic understanding of disease is the first step on the path from biological insight and target identification, to the development of mechanistic-based treatment. However, in order to translate genetics to biology, we require an ability to model the influence of genetic risk in a contextually appropriate system and to generate replicable disease-relevant readouts. The rapidly growing number of genetic risk variants and mutations associated with PD offers considerable challenges because modelling tens or hundreds of genetic factors cannot be sustainably achieved using traditional reductionist approaches. Moreover, this problem becomes more complex when considering risk variants in combination. However, this expanding risk landscape also offers opportunities. The more disease-linked genetic insight that can be modelled in a system, the more complete our understanding of disease biology will be and, as the molecular consequences of modelling risk coalesce, the more certainty we can have that these resulting events are disease-related. The application of large-scale iPSC models, with robust and reproducible molecular readouts, offers us the ability to assess the biological consequences of genetic risk factors in a disease appropriate cellular context.

Here we generated genetic, epigenetic, regulatory, cellular imaging, and transcriptomic data for 95 iPSC lines. These samples included healthy controls, PD patients with fully penetrant mutations in *SNCA*, mutations with reduced penetrance in *LRRK2*, and risk variants in *GBA1* as well as unaffected mutation carriers and individuals with iPD. Notably, there exists extensive biologic, clinical, and imaging data on each of the subjects from whom the lines were derived. Thus, the data described in the current study can be combined and compared with data collected on these subjects including longitudinal blood RNA-seq ^29^, CSF markers ^30^ and clinical data ^31^. Although we generated very large datasets totalling over 20 terabytes of data, we have sought to make these data available and accessible through the deposition of processed datasets, detailed experimental procedures, and data processing pipelines using the website for PPMI, https://www.ppmi-info.org/. In addition, we have created a dynamic browser (https://www.foundinpd.org) that allows users to interact with the data and to examine the features captured by FOUNDIN-PD at loci of interest and in genetic, phenotypic, and cellular subsets.

In characterising the first data release from the FOUNDIN-PD resource, we show that the large-scale differentiation process is robust and reproducible across technical replicates, but is variable between lines. Molecular characterization of the differentiation process and of the terminally differentiated cells revealed transcriptional and epigenetic changes in line with neuronal development. Further, our data reveal that, in the context of transcriptional profiles, the DA neurons created here closely model those from the adult human brain. Our work, combining previously published unbiased GWA-derived association loci with scRNA-seq data from FOUNDIN-PD, showed that the DA neurons generated here are an appropriate cellular context to model complex genetic risk. We believe that these data will also begin to offer insights into the mechanisms of disease-related loci by providing regulatory and expression information that has not been previously available.

During the course of this resource generating study, some important lessons were learned. Although the differentiation of multiple iPSC lines using a small-molecule approach produced a highly enriched neuronal culture (up to 93% *MAP2*+), there was also a variable amount of DA neurons (5-42% *TH*+) and a small percentage of non-neuronal cell types (2%). This variation was not related to batch, genetic sex, or the robustness of the differentiation protocol, as the technical replicates showed a very high correlation in subsequent rounds of differentiation. Such variability in the proportion of target cells produced by iPSC lines have been recently reported ^15^ and are mainly attributed to cell-intrinsic factors maintained over multiple freeze-thaw cycles. One of the factors driving inefficient differentiation towards specific cell types seems to be the heterogeneity of endogenous WNT signalling between iPSC lines ^32^ meaning that efficient patterning to dopaminergic neurons is dependent on the concentration of the GSK3 Inhibitor/WNT activator (CHIR99021) and would need to be optimised for each iPSC line ^33,34^. However, performing such optimization for all FOUNDIN iPSC lines would have been costly and time consuming. To minimise such variability between iPSC cell lines, researchers developed strategies such as the selection of well characterised cell lines for specific applications ^35,36^ and large-scale collaborative projects ^37^. However, such a strategy can only be applied to projects developed with a small number of iPSC lines.

The inclusion of single cell methods, which emerged into general usage during the execution of this study, has clearly been of great benefit to FOUNDIN-PD. These data help overcome the cell type heterogeneity of differentiated “mixed” cultures, provide a cellular context for genetic risk and also have the capacity to reveal transcriptomic and regulatory features specific to the disease-relevant cellular context. Therefore the role of bulk RNAseq indeed has changed. The bulk RNAseq is now complementary to the single-cell RNAseq data and provides certain benefits which the single-cell data cannot including the much higher sequencing depth which therefore allows investigating of splicing events and isoforms, which can be combined with deconvolution analysis. Overall, based on our observations thus far, the expansion of these methods to include combined single cell transcriptomics and ATAC-seq, single cell HiC, single cell chromatin immunoprecipitation methods to reveal transcriptional binding factor targets, and single cell proteomics will add more resolution to the FOUNDIN-PD study and more disease-relevant insights. Inclusion of these single cell data will be a key part of the next stage of FOUNDIN-PD.

Finally, we believe that longitudinal imaging of intact cells can complement the molecular analyses and add significantly to the characterization of patient-derived iPSC lines and to our goal to conduct functional genomics for PD ^39^. FOUNDIN-PD includes an extensive set of molecular analyses, but we recognize that some potentially important classes of bioactive molecules (e.g., lipids, metabolites) and functions (e.g., electrical activity) were not measured. For some assays, important subcellular spatial relationships of the macromolecules are necessarily lost during sample preparation. Imaging provides a method of studying cells as intact living systems, preserving critical components and their spatial relationships *in situ*, and enabling functional measurements relevant to PD that would be difficult to infer from reductionist molecular analyses. As noted above, there are inherent challenges associated with understanding how genetic variants implicated in PD contribute to disease. The effect size of individual variants is often small, making functional effects hard to detect, and it may be the case that substantial disease risk for an individual is conferred through the combined non-linear effects of multiple variants. If so, then combining imaging with molecular analyses may be particularly helpful because it provides an approach to study the integrated effect of genetic variants on specific cell functions relevant to disease. Finally, imaging data are especially amenable to powerful machine learning-types of analyses, which can be used to discover biological insights from images that elude the human eye ^40^ and provide a computational framework for integrating other data types, including types of multi-omic data produced by FOUNDIN-PD. Indeed, next steps include multi-omics data integration to systematically understand and identify PD-relevant pathways. This integration provides an opportunity to investigate PD-relevant biological pathways at multiple layers like genotype, chromatin and transcript level.

## Limitations of the Study

The efficiency and reproducibility of the DA neuron differentiation protocol was not previously explored on the large set of iPSC lines. Here we identified as expected that there is substantial line to line variation. Interestingly, we were able to identify early expression markers that correlate with the potential to generate high levels of DA neurons in the FOUNDIN-PD cell lines, therefore it is tempting to speculate that sorting iPSC based on a high expression of for example *SRSF5* may improve differentiation efficiency. These results are in line with a previous report showing that expression markers detected at the iPSC stage can robustly predict differentiation capability^15^. While the emergence of single cell molecular methods relieves some concerns regarding cellular heterogeneity, improving differentiation consistency line-to-line would be of benefit. We acknowledge that this dataset is underpowered to reveal all but the strongest of mechanisms associated with complex disease risk loci. Additionally, while iPSCs are a useful model, they have limitations, including the fact that DNA methylation signatures from the donor are not preserved upon reprogramming and therefore they lack ageing related phenotypes, which is the biggest common risk factor for PD. While the number of lines required to generate insights at the remaining loci will vary from risk allele to risk allele, we believe that the next stage of FOUNDIN-PD should include a significant increase in scale. Notably, as initiatives such as the Global Parkinson’s Genetics Program (GP2) focus on diversifying the ancestral basis of our genetic understanding in PD ^38^, efforts such as FOUNDIN-PD should also prioritise the generation of reference data in well-powered ancestrally diverse systems. We also see the value in diversifying our terminal differentiation target to include other cell types potentially relevant for PD.

## Conclusion

Overall, we present here the first data release of the FOUNDIN-PD project, which includes multi-omics and imaging data on iPSCs differentiated to DA neurons of 95 PPMI participants harbouring a range of genetic risk, from fully penetrant causal mutations to carriers of combinations of risk alleles identified by GWAS. We believe the FOUNDIN-PD data will serve as a foundational resource for PD research with easily accessible data and browsers designed for basic scientists. This dataset will help the community to better understand the mechanisms of PD, identify new disease-relevant targets and potentially impact the development of novel therapeutic strategies.

## Author contributions

Study design: All

Funding acquisition: ABS, PH, MRC, CB, JRG, SF, KVKJ, DWC

Data analysis: VB, IV, DWC, CB, AI, NS, EB, JRG, MC, SF, XR, MRC, FPG, EH, EA

Statistical analysis: VB, IV, DWC, CB, AI, NS, EB, JRG, FPG

Manuscript drafting: CB, EB, XR, VB, DWC, PH, MMC, SF, ABS, MRC

Manuscript revision: All

DA neuron culture: EB, SB, MMC

Assay preparation and processing: EB (ICC, scRNA-seq, scATAC-seq, HiC-seq), SB (ATAC-seq), NF & PR (scRNA-seq), XR (Genotyping, Methylation, ATAC-seq), CB (Genotyping, Methylation, HiC-seq), MMC (Imaging), CD (ATAC-seq, HiC-seq), JB (Genotyping, Methylation), FPG (Methylation, HiC-seq), BM, RR, AL, JA, AC (RNA isolation for small and long RNA library preparation; library preparation and sequencing)

SOPs: CB, IV, EB, PR, SB, XR, MMC, VB

Browser: DWC, MW, RS, IV

## Inclusion and diversity statement

We support inclusive, diverse, and equitable conduct of research.

## Acknowledgements

We would like to thank all of the subjects who donated their time and biological samples to PPMI, without whom we could not have done this study. This work is supported by the Michael J. Fox Foundation for Parkinson’s Disease Research and is part of the PD Pathogenesis consortium. Cell lines used in the analyses presented in this article were obtained from the Golub Capital iPSC Parkinson’s Progression Markers Initiative (PPMI) Sub-study (www.ppmi-info.org/cell-lines). Data used in the preparation of this article were obtained from the PPMI database (www.ppmi-info.org/data). As such, the investigators within PPMI contributed to the design and implementation of PPMI and/or provided data and collected samples but did not participate in the analysis or writing of this report. For up-to-date information on the study, visit www.ppmi-info.org. PPMI – a public-private partnership – is funded by The Michael J. Fox Foundation for Parkinson’s Research and corporate sponsors, including: Abbvie, AcureX Therapeutics, Allergan, Amathus therapeutics, Avid Radiopharmaceuticals, BIAL Biotech, Biogen, Biolegend, Briston-Myers Squibb, Calico, Celgene, Denali, 4D Pharma, GE Healthcare, Genentech, GlaxoSmithKline, Golub Capital, Handl Therapeutics, Insitro, Janssen neuroscience, Lilly, Lundbeck, Merck, Meso Scale Discovery, Neurocrine Biosciences, Pfizer, Piramal, Prevail Therapeutics, Roche, Sanofi Genzyme, Servier, Takeda, Teva, UCB, Verily and Voyager Therapeutics. An up to date list of all PPMI Industry Partners can be found at http://www.ppmi-info.org/about-ppmi/who-we-are/study-sponsors/. This work is supported in part by the Intramural Research Program of the National Institute on Aging, National Institutes of Health, part of the Department of Health and Human Services; project ZO1 AG000949. This study utilized the high-performance computational capabilities of the Biowulf Linux cluster at the National Institutes of Health, Bethesda, MD, USA. (http://biowulf.nih.gov). Additional support include ERACoSysMed2; PD-Strat. Multi-dimensional stratification of Parkinson’s disease patients for personal interventions (FKZ 031L0137A). VB is supported by a Career Development Fellowship at DZNE Tuebingen. The authors would like to thank the NIH Intramural Sequencing Center (NISC), the American Genome Center and the Platform foR SinglE Cell GenomIcS and Epigenomics (PRECISE) for sequencing services and Phase Genomics for HiC library construction. Additional support for this work came from RF1 AG1058476, U54 NS191046 and R37 NS101966 to SF.

## STAR Methods

### RESOURCE AVAILABILITY

#### Lead contact

⍰ Further information and requests for resources and reagents should be directed to and will be fulfilled by Cornelis Blauwendraat.

#### Materials Availability

⍰ All iPSC lines used in this study are available upon request at https://www.ppmi-info.org/access-data-specimens/request-cell-lines/.
⍰ Extensive protocols and all data generated is available at https://www.ppmi-info.org/=>Access-data-specimens/Download-data/Geneticdata/FOUNDIN-PD.

#### Data and code availability

⍰ All code from this study is publicly accessible and available at https://github.com/FOUNDINPD.

⍰ All data from this study are publicly accessible and available at https://www.ppmi-info.org/.

### EXPERIMENTAL MODEL AND SUBJECT DETAILS

The induced pluripotent stem cell (iPSC) lines (n=134) used were obtained from the Parkinson’s Progression Marker Initiative (PPMI; https://www.ppmi-info.org/). Each cell line vial was identified with an unique barcode and accompanied by a quality control certificate for showing normal karyotype, pluripotency and a negative test for mycoplasma. Frozen cell line stocks were quickly thawed at 37 °C, washed once with DMEM/F-12 (Gibco) to remove cryopreservation medium, resuspended in Essential 8 Flex (E8) or Essential 6 (E6) media (both Gibco) supplemented with 10 µM Y-27632 and plated on matrigel (Corning)-coated plates. E8 and E6 media were supplemented with growth factors to become equivalent in composition. Cells were kept in culture for about one month (5 passages) to allow recovery from thawing and to obtain enough cells for differentiation and assays on day 0 (iPSC state).

### Overview of PPMI iPSC lines included in FOUNDIN-PD

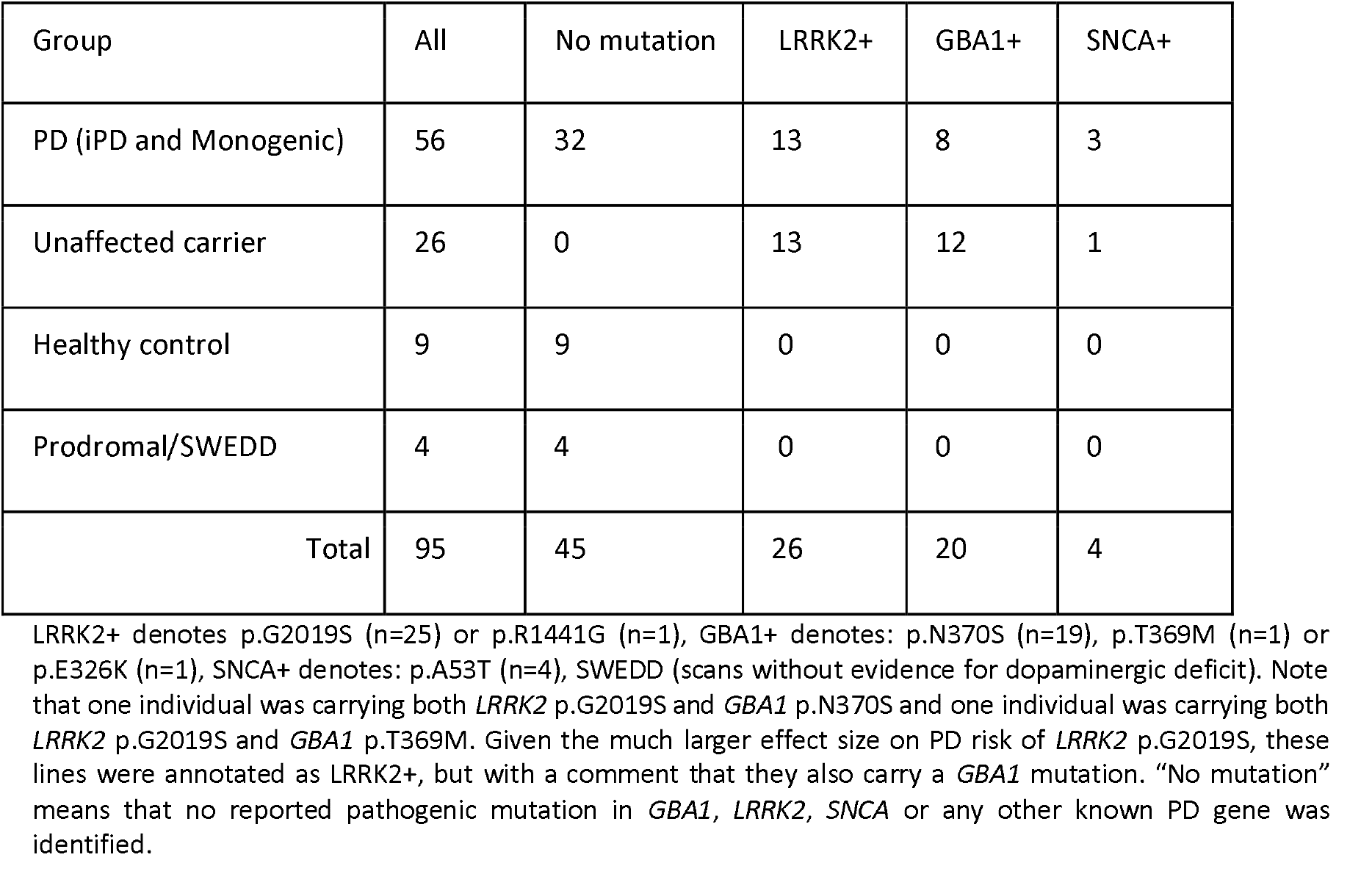

Overview of PPMI iPSC lines included in FOUNDIN-PD.
| Group | All | No mutation | LRRK2+ | GBA1+ | SNCA+ |
| --- | --- | --- | --- | --- | --- |
| PD (iPD and Monogenic) | 56 | 32 | 13 | 8 | 3 |
| Unaffected carrier | 26 | 0 | 13 | 12 | 1 |
| Healthy control | 9 | 9 | 0 | 0 | 0 |
| Prodromal/SWEDD | 4 | 4 | 0 | 0 | 0 |
| Total | 95 | 45 | 26 | 20 | 4 |

LRRK2+ denotes p.G2019S (n=25) or p.R1441G (n=1), GBA1+ denotes: p.N370S (n=19), p.T369M (n=1) or p.E326K (n=1), SNCA+ denotes: p.A53T (n=4), SWEDD (scans without evidence for dopaminergic deficit). Note that one individual was carrying both *LRRK2* p.G2019S and *GBA1* p.N370S and one individual was carrying both *LRRK2* p.G2019S and *GBA1* p.T369M. Given the much larger effect size on PD risk of *LRRK2* p.G2019S, these lines were annotated as LRRK2+, but with a comment that they also carry a *GBA1* mutation. "No mutation" means that no reported pathogenic mutation in *GBA1*, *LRRK2*, *SNCA* or any other known PD gene was identified.

## METHOD DETAILS

### Sample selection and batching

Upon receiving, all cell lines were NeuroChip array genotyped ^41^ to confirm sample origin and to assess if large genomic events occurred during reprogramming, iPSC culture and differentiation. The data were compared to donor (blood derived) whole-genome sequencing (WGS) to identify large genomic abnormalities. Of 134 subjects, 80 are males and 54 females. The cell line collection included healthy controls, PD cases without mutations in PD related genes and affected and unaffected individuals harbouring damaging point mutations including *SNCA* p.A53T, *LRRK2* p.G2019S, *LRRK2* p.R1441G, *GBA1* p.E326K, *GBA1* p.T369M and *GBA1* p.N370S. Note that one iPSC line carries both *LRRK2* p.G2019S and *GBA1* p.N370S and another iPSC line carries both *LRRK2* p.G2019S and *GBA1* p.T369M. Given the much larger effect size on PD risk of *LRRK2* p.G2019S, these lines were annotated as *LRRK2*+, but with a comment that they also carry a *GBA1* mutation. From the 134 cell lines, 95 passed QC and were selected for DA neuron differentiation and split into five batches (Table 1). One control cell line was included in each batch as a technical replicate (n=5) totaling 99 samples (Supplementary Table 1).

### Differentiation of iPSC into dopaminergic (DA) neurons

The PPMI iPSC lines were thawed and grown on matrigel (Corning)-coated plates with Essential 8 Flex (E8, Batches 1, 2 and 3) or Essential 6 (E6, Batches 4 and 5) media (both Gibco) for about one month (5 passages). Essential 6 medium was supplemented with growth factors to become equivalent in composition to Essential 8. Upon reaching 70-80% confluency, iPSC lines were dissociated into a single cell suspension with Accutase (Gibco) and plated at 200,000 cells/cm^2^ on matrigel-coated one-well plates (barcoded, Greiner) suitable for automated cell culture. Cells were grown until they covered the plate surface, usually 24-48 hours after single cell plating. The time required to reach confluence was variable and dependent on the growth rate of each iPSC line. The DA differentiation protocol was adapted from Kriks and collaborators ^9^ with minor modifications ^10^. Differentiations were carried out in an automated cell culture system ^11^ with manual replatings on day 25 and 32 for final differentiation and immunocytochemistry (ICC), respectively ^11^. Samples for assays were collected on days 0 (iPSC), 25 (mid-point) and 65 (DA neurons). For DNA assays, cells were dissociated with Accutase, washed once with PBS and spun down at 200x g. The cell pellet was snap frozen or processed according to assay protocols. Most of the RNA assays required snap frozen cells collected by scraping the plate surface with PBS or lysis buffer. Single-cell (sc) RNA-seq and scATAC-seq assays required a single cell suspension prepared in 0.04% human serum albumin (HSA)/PBS. All samples were stored at -80°C until further processing. For cryopreservation, day 25 DA neuron precursors were dissociated with Accutase, washed once with neurobasal medium (Gibco), resuspended in cold Synth-a-Freeze cryopreservation medium (Gibco) supplemented with 10µM Y-27632 and aliquoted into barcoded cryovials (NovaStora) at 10×10^6 cells/ml/vial (on ice). The cryovials were placed in CoolCell cell freezing containers (Biocision), kept overnight at -80°C and transferred to liquid nitrogen for long term storage.

### Immunocytochemistry (ICC) and image analysis

Cells were differentiated until day 65, fixed in 4% PFA, washed 3×5 min in PBS and blocked in 5% goat serum/1% bovine serum albumin (BSA)/0.1% Triton-X100/PBS for 1 h at room temperature (RT). Primary antibodies were applied overnight at 4°C and included *TH* (Pel-Freez Biologicals #P40101 and Merck Millipore #AB9702, both at 1:750 dilution), *MAP2* (Santa Cruz #sc-74421, 1:100) and TUJ1 (R&D #MAB1195, 1:500). After incubation with primary antibodies, cells were washed 3×5 min in PBS. Cells were incubated with second antibodies (AF488 and AF594, Invitrogen, 1:1000) for 2 h at RT followed by nuclear counterstaining with Hoechst 33342 (Invitrogen, 1:8000) for 30 min at RT. Finally, cells were washed 3×5 min in PBS and imaged with a CellVoyager 7000 (Yokogawa) confocal microscope and 20x objective. Images were analysed on Columbus (PerkinElmer) as described previously ^11^. The total number of *TH* (DA neuron) and *MAP2* or TUJ1 (neuron) positive cells was estimated and normalized to the total number of nuclei. Data is represented as the percentage of positive cells per 30 fields.

### Longitudinal image analysis of iPSC DA neurons

Frozen day 25 DA neuron precursors were thawed and replated at a density of approximately 450,000 cells/cm^2^ onto dishes coated with 0.1mg/ml poly-l-ornithine (PLO), 5µg/ml laminin, and 5µg/ml fibronectin in NB/B27 medium prepared as described ^10^ with the addition of 10µM ROCKi and 100µg/ml matrigel (Corning). The media was changed 4 h later to remove ROCKi. DA neurons were matured in NB/B27 medium, then replated into 96-well plates on day 49. On day 50, cells were transduced with synapsin-driven GFP via lentivirus (SignaGen), followed by a media change the next day. Cells were imaged daily from approximately day 54 through 66 using robotic microscopy, a previously described automated imaging platform ^23,24^. Images obtained from 8 consecutive days were processed using custom programs in Galaxy ^42^; ^43^ to assemble arrays of images into montages representing each well, and to stack montages across timepoints. Neuron survival was analyzed using a custom program written in MATLAB. Live neurons expressing GFP were selected for analysis only if they had extended processes at the first timepoint. Neurons were tracked longitudinally across timepoints until death, and survival time was defined as the last timepoint a neuron was seen alive. The survival package for R statistical software was used to construct Kaplan-Meier curves from the survival data^24^. Survival functions were fitted to these curves to derive cumulative risk-of-death curves. Statistical differences between groups were analyzed using the Cox mixed-effects model.

### Methylation library preparation and data-processing

DNA was extracted from each timepoint using standard phenol:chloroform extraction. DNA from day 0 and day 65 underwent Bi-sulfite conversion using the EZ-96 DNA Methylation Kit (Zymo Research). Bisulfite converted DNA was then put through the standard Infinium HD array based methylation assay (Illumina) with Illumina Infinium HumanMethylation EPIC BeadChips. Raw signal intensity data were processed from raw idat files through a standard pipeline using Meffil ^44^. A number of standard quality control steps were performed to these data prior to normalization including: sample origin confirmation based on SNP presence on array, sex concordance check, methylated versus unmethylated ratio, low bead numbers, control probes quality and, finally, general outlier samples were identified using principal component analysis and excluded. Subsequently, the quality controlled data was normalised using quantile normalisation. The analysis pipeline can be found here: https://github.com/FOUNDINPD/METH.

### Bulk ATAC sequencing library preparation, sequencing and data-processing

Bulk ATAC-seq data was generated from all batches at all timepoints. Cells at each timepoint were collected using Accutase (Gibco) to make a single cell suspension and 75,000 cells per sample were aliquoted for bulk ATAC-seq. Standard procedures with slight modifications were used ^45^. In brief, cells were lysed (10mM Tris-HCl, 10mM NaCl, 3mM MgCl2, 0.1% (v/v) NP-40), nuclei were then spun down, resuspended in transposition buffer (TD buffer, Tn5 Transposase from the Illumina Tagment DNA Enzyme and Buffer kit) and incubated for 30 min at 37°C. After incubation, DNA was isolated using Qiagen MinElute Reaction Cleanup Kit (Qiagen) according to manufacturer’s recommendations. DNA was eluted in 10µl of EB buffer (10mM Tris-Cl, pH 8.5) and then frozen at -80°C.

Libraries were prepared by combining transposed DNA with NEBNext High-Fidelity 2X PCR Master Mix (New England Biolabs) and 1.25µM indexing primers (Ad1_noMX primer and Ad2.x indexing primer ^45^ or IDT for Illumina dual index primers (Illumina, Nextera DNA UD Indexes Set A Ref 20025019). Standard PCR conditions for NEBNext High-Fidelity 2X PCR Master Mix were used with 10-12 cycles completed on each library. Libraries were purified using AMPure XP (Beckman Coulter) beads with the manufacturer’s protocol for double-sided purification. Quality assessment of libraries was measured on an Agilent High Sensitivity DNA analysis chip (Agilent) to determine average library size and concentration. The concentration of each library was verified by Qubit Fluorometric Quantification (Thermo Scientific) before sequencing. Batch 1 libraries were sequenced at the NIH Intramural Sequencing Center (NISC) on an Illumina NovaSeq, with 50bp paired-end (PE) reads. Batches 2-5 were sequenced at The American Genome Center (TAGC) at the Uniformed Services University on an Illumina NovaSeq with 100bp PE reads. Fastq files for each sample were assessed using FastQC (v0.11.9) and reads were aligned to GRCh38 using Bowtie2 (v2.4.1; Langmead and Salzberg, 2012) in local mode. Reads mapping to ChrM and ChrUn were filtered out and samples with less than 20 million PE reads remaining were removed from analysis. MACS2 was used to call peaks ^46^. The full analysis pipeline can be found here: https://github.com/FOUNDINPD/ATACseq_bulk.

### HiC sequencing library preparation, sequencing and data-processing

HiC sequencing data were generated from batch 1 day 0 and day 65 samples. Library preparation was performed by Phase Genomics (https://phasegenomics.com/) using their standard protocol. Fastq files from each lane were merged to give each sample two read fastqs. Fastqc was run on all sample fastq files before further analysis. The Juicer pipeline was used to obtain high resolution contact maps and loop regions for each sample ^47^. Preliminary testing indicated excessive mitochondrial data in samples, so the pipeline was altered to remove mitochondrial reads after mapping. The Juicer pipeline incorporates the Burrows-Wheeler Aligner (BWA) to map fastqs to a reference genome ^48^. Loop regions in samples were detected using the HiCCUPs algorithm included in the Juicer pipeline. These regions were saved in .bedpe files and used for further analysis. Loop region overlap was calculated between samples and with public PsychENCODE data ^49^. The HiCCUPSDiff tool was used to detect differential loops between day 0 and day 65. Heatmaps were generated for each sample and each chromosome to visualise chromatin interactions using the HiCExplorer tool ^50^. The analysis pipeline can be found here: https://github.com/FOUNDINPD/HiC_Pipelines.

### Bulk RNA sequencing library preparation, sequencing and data-processing

Bulk RNA sequencing data was generated from all batches and all timepoints. RNA was isolated using Qiagen’s “Purification of miRNA from animal cells using the RNeasy Plus Mini Kit and RNeasy MinElute Cleanup Kit” using protocol 1 to purify total RNA containing miRNA. Briefly, cells were lysed with Guanidine-isothiocyanate and homogenised with QIAshredder, then passed through a gDNA Eliminator spin column. The lysate was combined with ethanol to bind RNA to the spin column while contaminants are washed away. Samples were separated into 4 different RNA isolation protocols dependent on the sample’s cell counts (target of 1.3-4 million cells per column). Samples with 1.33-4 million cells/vial were isolated using 1 column. Samples with 4.61-7.86 million cells/vial were isolated on 2 columns with 2.3-3.93 million cells/column. Samples with 8.17-12 million cells/vial were isolated on 3 columns with 2.72-4.0 million cells/column. Samples with 12.75-52 million cells/vial were isolated on 3 columns with 4 million cells/column and the leftover lysate was stored. High quality total RNA (containing miRNA) was then eluted and used for either bulk RNA sequencing or small RNA sequencing library preparation. Libraries were prepared using the SMARTer Stranded Total RNA Sample Prep Kit - HI Mammalian (Takara Bio USA, Inc.), which incorporates both RiboGone and SMART (Switching Mechanism At 5’ end of RNA Template) technologies to deplete nuclear rRNA and synthesise first-strand cDNA. This along with PCR amplification and AMPure Bead Purification generates Illumina-compatible libraries. Using the total RNA stock concentration, we determined the volume needed for 1ug RNA input. Samples were concentrated by SpeedVac or diluted with nuclease-free water to obtain a volume of 9µl per sample. Addition of buffers and enzymes including RNase H, DNase I, and 10X RNase H Buffer along with three PCR reactions and a 1.8X bead purification removed specific rRNA sequences (5S, 5.8S, 12S, 18S, and 28S). rRNA depleted RNA was fragmented at 94°C for 3 min and immediately placed on ice. Master mix containing reverse transcriptase and an oligonucleotide added to samples and incubated in a preheated thermal cycler to convert RNA to single-stranded cDNA. cDNA was purified at 1X ratio with AMPure beads. Unique dual-indexed PCR primers (allowing for multiplexing) combined with SeqAmp DNA Polymerase were added to each first-strand cDNA. Using 12 cycles on a preheated thermal cycler, cDNA was amplified into RNAseq libraries. AMPure Bead purification (1X), 80% ethanol wash, and elution of 34µl with nuclease-free water generated final libraries ready for Illumina sequencing. 2µl of cDNA library placed on a well plate with 2µl Sample Buffer and analysed on 4200 TapeStation to determine peak range (bp). Concentration of libraries determined by 40K, 80K dilutions on Kapa SYBR Fast qPCR (Roche). Libraries were pooled together into 2 pools with a concentration of 60pM and volume of 100µl and sequenced on an iSeq 100 300-cycle flow cell. Libraries were normalised based on these results. Libraries were re-pooled together with a final concentration of 5000pM and final volume of 220µl, concentration obtained by QuantStudio. Pool was run on a NovaSeq 6000 S1 200-cycle flow cell with a loading concentration of 1,500pM and volume of 100µl with a 20% PHiX spike-in, with the following parameters: 100 × 9 × 9 (+7 dark cycles) x 100. The sequencing depth was a minimum of 30M read pairs per sample. The bcl files were demultiplexed using bcl2fastq v2.19.1.403 (Illumina) using default parameters. Reads were trimmed with cutadapt v2.7 ^51^ to remove the first three nucleotides of the first sequencing read (Read 1), which are derived from the template-switching oligo. Trimmed reads were aligned to the GRCh38 genome using STAR v2.6.1d ^52^. Following genome alignment, reads were counted with featureCounts v1.6.4 ^53^, (part of the subread package) using a non-redundant genome annotation combined from GENCODE 29 and LNCipedia5.2 ^54^ (https://github.com/FOUNDINPD/annotation-RNA). Count data was loaded into R v3.6.3 for analysis. Normalised counts, variance stabilising transformation, and differential expression analysis were performed using DESeq2 v1.26.0 ^55^, and CPM values were generated using edgeR v3.28.1 ^56^. Heatmaps were created using the pheatmap v1.0.12 package in R. Trimmed fastq files were also quasi-mapped to the same annotation using salmon quant v1.2.2 ^57^. In order to identify upregulated and downregulated genes from day 0 to day 65, differentially expressed genes (defined as baseMean > 100, adjusted p < 0.01, and absolute value of the log2 fold change > 1) were further filtered using a general linearized model, retaining genes that have a slope > 0.05 for upregulated genes and a slope < -0.05 for downregulated genes. Gene ontology analysis was performed on these upregulated and downregulated genes with GOfuncR 1.6.1, using the refine function with a FWER = 0.1, and GOxploreR 1.1.0 ^58^was used to remove redundant GO terms Parameters used for genome alignment, annotation, and quasi-mapping are described on GitHub. The analysis pipeline can be found here: https://github.com/FOUNDINPD/bulk_RNASeq.

### Small RNA sequencing library preparation, sequencing and data-processing

Small RNA sequencing data were generated from all batches and all timepoints. RNA was isolated in the same manner as for bulk RNA sequencing, using Qiagen’s “Purification of miRNA from animal cells using the RNeasy Plus Mini Kit and RNeasy MinElute Cleanup Kit” using protocol 1. Small RNA libraries were made using the NEXTFLEX Small RNA v3 kit (PerkinElmer), followed by 3’ adapter ligation and excess 3’ adapter removal according to manufacturer’s protocol. Excess adapter inactivation was not performed. 2µl of the inactivation ligation buffer were used without enzyme in lieu of the inactivation step. 5’ adapter ligation and reverse transcription was performed per manufacturer’s protocol. 62.5µl of cDNA, beads, and isopropanol solution was transferred instead of 70µl to help reduce adapter dimer moving forward to PCR. Libraries of appropriate size were collected using gel purification. Purified libraries were quantified using the high sensitivity DNA kit on the Bioanalyzer (Agilent). Equimolar pools were made and sequenced on a Hiseq 2500 at 8pM. The bcl files were demultiplexing using bcl2fastq. Small RNA sequencing reads (fastq files) were processed using the exceRpt pipeline. The pipeline was run using the RANDOM_BARCODE_LENG*TH*=4 parameter to trim off the random 4-bp ends in NEXTFLEX sequencing data along with the Illumina (TruSeq) smallRNA adapters. All other parameters were set to defaults. Pipeline was run using a custom transcriptome database composed of human sequences from mirBase 22, gencode 28, piRBase and tRNAscan-SE. Following the pipeline run on each sample an R summary script (mergePipelineRuns.R) was run which generates raw read alignment counts, RPMs and QC metrics for all small RNA species across all samples. Expression of small RNAs that were consistently increasing over timepoints were investigated for their expression patterns using data from a small RNA tissue atlas (Aslop et al, in preparation). The analysis pipeline can be found here: https://github.com/FOUNDINPD/exceRpt_smallRNAseq.

#### Single-cell (ATAC and RNA) library preparation, sequencing and data-processing

Cells harvested on day 65 of differentiation were processed following the 10x Genomics single-cell (sc) RNA and ATAC sequencing protocols to generate DNA libraries. To note, batch 1 cells processed for scRNA-seq were generated from a second run of differentiation, since this assay was included later in the study. Additionally, scATAC-seq was performed only for cells from batches 4 and 5. For scRNA-seq, the libraries comprised standard Illumina paired-end constructs which begin with P5 and end with P7. The 16bp 10X barcodes are encoded at the start of TruSeq Read 1, while 8bp sample index sequences are incorporated as the i7 index read. TruSeq Read 1 and Read 2 are standard Illumina sequencing primer sites used in paired-end sequencing. TruSeq Read 1 is used to sequence 16bp 10x barcodes (cell identifier) and 12bp UMI (transcript identifier). scATAC-seq libraries compatible with Illumina sequencing were generated by adding a P7 and a sample index via PCR. Sequencing was performed on Illumina NovaSeq. Libraries were sequenced at a minimum depth of 20,000 read pairs per cell for scRNA-seq and 25,000 read pairs per nucleus for scATAC-seq.

#### scRNA-seq

The BCL files obtained after sequencing were demultiplexed into FASTQ files using the cellranger *“mkfastq”* software and unique molecular identifier (UMI) gene counts were calculated by cellranger *“count”* software (v3.1.0) ^59^. UMI gene counts for each sample were merged into a table and imported into R (v3.6.0). We used Seurat (v3.1.1) ^60^within the R environment for filtering, normalisation, integration of multiple single-cell samples, unsupervised clustering, visualisation and differential expression analyses. The following data processing was done: (1) Filtering. Cells with less than 1,000 and more than 9,000 genes expressed (≥1 count) were filtered out, and only genes that were expressed in at least 100 cells were kept. Moreover, cells with more than 20% of counts in mitochondrial genes were filtered out. After filtering, there were 34,960 genes in 416,216 cells; (2) Data normalisation and integration. Gene UMI counts for each cell were normalised using the *“SCTransform”* function in Seurat. Integration of scRNA-seq data from multiple samples was performed using top 3000 variable features and top 3 samples as reference with the highest number of cells; (3) Clustering and visualisation. Clustering was performed using *“FindClusters”* function with default parameters except resolution was set to 0.5 and first 30 PCA dimensions were used in the construction of the shared-nearest neighbour (SNN) graph and to generate 2-dimensional embeddings for data visualisation using UMAP; (4) Differential expression analyses: We used *“FindAllMarkers”* function with default parameters and only tested genes that are detected in a minimum of 40% of cells in either of the two clusters. Genes with an adjusted p value <0.05 were considered to be differentially expressed. The pipelines used in this study are available at https://github.com/FOUNDINPD/FOUNDIN_scRNA.

#### scATAC-seq

The BCL files obtained after sequencing were demultiplexed into FASTQ files using the cellranger-atac “mkfastq” software and unique molecular identifier (UMI) counts were calculated by cellranger-atac “count” software (v1.2.0) ^61^. Peaks for each sample were merged into a table and imported into R (v3.6.0). We used Seurat (v3.2.0), Signac (v1.1.0) ^62^ and Harmony (v1.0) ^63^ within the R environment for filtering, normalisation, integration of multiple single-cell samples, unsupervised clustering, visualisation and predicting the cell types. The following data processing was done: (1) Filtering. We kept the cells with minimum 1,000 peaks (≥ 1 count), respectively and the peaks that were called in at least 100 cells. Moreover, cells with more than 20% of counts in mitochondrial DNA were filtered out. After filtering, there were 459,495 peaks in 139,659 cells; (2) Data normalisation and integration. Peak counts for each cell were normalised using the *“RUNTFIDF”* function in Signac that performs term frequency-inverse document frequency normalisation followed by SVD decomposition to generate latent semantic indexing (LSI). Integration of scATAC-seq data from multiple samples was performed using the *“RUNHarmony”* function with LSI reduction; (3) Clustering and visualisation. Clustering was performed using the *“FindClusters”* function with default parameters except resolution was set to 0.1 or 0.2. First 30 harmony dimensions were used to generate 2-dimensional embeddings for data visualisation using UMAP; (4) Predicting cell types: Fragments in the genes (extended 2kb upstream) were calculated for each cell to generate a gene activity matrix and normalised the data using the *“LogNormalize”* method. Cell types were predicted using scRNA-seq data as a reference and scATAC-seq data as a query for *“FindTransferAnchors”* and *“TransferData”* functions. Prediction often results in heterogeneous cell type annotation with-in the same cluster. We assigned the cell type to a cluster with the maximum occurrence. The neuroepithelial-like cluster was separated using 0.2 resolution. The pipelines used in this study are available at https://github.com/FOUNDINPD/FOUNDIN_scATAC.

### Prediction of neuronal differentiation efficiency using bulk RNA-seq data at day 0

To test the predictive value of the genic expression profile in iPSC for neuronal differentiation efficiency, we performed supervised machine learning (logistic regression) on the DESeq2 v1.26.0 ^55^ normalised count expression values for genes at day 0 and estimated DA neurons fractions from the differentiated cell lines at day 65. DA neuron fractions were calculated from scRNA-seq data, based on the total number of cells and the number of cells in the ‘Dopaminergic Neurons’ cluster (see the Method section for scRNA-seq). Cell lines were classified into high (n=62) and low (n=21) differentiation efficiency classes based on the relative abundance of the DA neurons at day 65; as a threshold for classification, we used first quartile value of cell percentages (Q25=15.7%), as it was best separating the two observed distribution peaks of DA neuron counts across the cell lines.

To reduce possible bias in the predictive model, we used a full set of reliably expressed genes (threshold of inclusion mean normalised count >=50). As we did expect a significant number of genes to be highly correlated to one another in their expression, with multitude of them being possibly relevant for prediction, and the total number of relevant features for our model is unknown, we resolved to using elastic net regularisation approach, which combines both lasso regression (shrinking less important features and pruning some) and ridge regression (assigns proportional coefficients to highly correlated possibly relevant features and prevents model overfitting) to equal degree (alpha=0.5) with a penalty lambda equal to 0.22. To further control for possible overfitting, repeated (100 times) 5-fold cross-validation was performed using the *“cv*.*glmnet”* function. Data preprocessing and logistic regression was executed in R (v3.6.3) ^64^, using packages caret (v6.0-86) ^65^ for model training and glmnet (v4.0) ^66^ for elastic net regularisation of the model and repeated cross-validation. As the sample size is small and imbalanced, we directly tested the relation of the resulting predictive candidate genes’ expression to the percentage of DA neurons in each cell line. We performed Spearman’s rank correlation test, using R package stats (v3.6.3) ^64^. Benjamini & Hochberg procedure was used for multiple testing corrections of p-value.

### MAGMA to identify causative cell types

Expression gene profiles obtained from the scRNA-seq dataset were used to test for a cell type association with PD. We used the R package MAGMA_Celltyping (v1.0.0, https://github.com/NathanSkene/MAGMA_Celltyping), which utilises MAGMA ^25^ software package as a backend, to identify cell types positively associated with the common-variant genetic hits from the most recent PD GWAS ^2^. LD regions were calculated using the European panel of 1000 Genomes Project Phase 3 ^67^. The cell type enrichment analysis was performed on 5000 subsampled cells from each scRNA-seq cluster.

### Single cell expression quantitative trait loci analysis

Variants from the whole genome sequencing data were correlated with normalised average gene expression levels per cell cluster by performing single cell expression quantitative trait loci analysis. After quality control, 77 samples were included for analysis and expression data were filtered for 0.025 average expression in all samples. Then genes were removed with zero expression in 15 or more samples resulting in expression of 1256 genes across 90 risk variants. eQTL analysis was performed using MatrixEQTL ^68^ including variants with minor allele frequency >5% and using the following covariates: batch, sex, age of donor, *GBA1, SNCA, LRRK2*, phenotype, *TH*+ levels, *MAP2*+ levels, number of cells, reads per cell, total genes detected, and median UMI counts per cell. Overlap between eQTL variants and GWAS was determined using the most recent PD GWAS ^2^. For GWAS loci of interest, violin plots were generated to visualise the correlation between genotype and gene expression. Additionally, LocusZoom ^69^ and LocusCompare ^70^ plots were generated to visualise correlations between GWAS signal and eQTL signal and the PD GWAS locus browser was used for loci numbering and prioritisation ^26^. The analysis pipeline can be found here: https://github.com/FOUNDINPD/SCRN_EQTL_v2. Bulk eQTL analysis was performed separately on day 0, 25, and 65 data using tensorQTL ^71^ and included estimated cell fractions as covariates. The estimated cell fractions were generated using the Scaden ^72^ deconvolution tool trained on the day 65 single-cell data.

### FOUNDIN-PD browser

Architecturally, the FOUNDIN-PD portal is a single-page application (SPA) framework where a public javascript application interacts with a secured JSON API to build the user DOM within the user browser. The client-side nature of the application allows for dynamic interactions with the user with low latency and high scalability, leveraging the fact that many users will leverage modern computing and browsing capabilities. At a granular level, the FOUNDIN-PD application is based on JavaScript ECMAScript 2016 and builds upon Vega.js (vega.github.io; version 5.22) visualisation grammar and D3.js (https://d3js.org; version 6) for dynamic responsive graphing. The API is within a sharded MongoDB 4.2 (https://mongodb.com) framework on a CentOS8 cloud server using an NGINX (https://nginx.org; 1.18) proxy, NodeJS 12 middleware (https://nodejs.org), to provide a protected JSON API. API data is secured using JSON/JWT authentication via Auth0 (https://auth0.com) and Google OAU*TH* 2.0 (https://oauth.net/2/) for the identification of users.

### Quantification and Statistical Analysis

All analysis details pipelines for data in these studies can be found in the Methods sections and on the FOUNDIN-PD GitHub (https://github.com/FOUNDINPD). Additionally detailed SOPs for each assay can be found at the PPMI portal (https://www.ppmi-info.org/).

## Supplemental Table Legends

**Supplemental Table 1. Included iPSC overview with meta-data. Related to Figure 1**.

**Supplemental Table 2. Single cell RNA-seq cell type cell marker gene overview. Related to Figure 2**.

**Supplemental Table 3. Single cell RNA-seq cell type percentage per iPSC line. Related to Figure 2**.

**Supplemental Table 4. Bulk RNA-seq gene-set enrichment across timepoints (upregulated). Related to Figure 3**.

**Supplemental Table 5. Bulk RNA-seq gene-set enrichment across timepoints (downregulated). Related to Figure 3**.

**Supplemental Table 6. MAGMA cell type enrichment results using scRNA-seq cell types. Related to Figure 6**.

**Supplemental Table 7. FOUNDIN-PD expression quantitative trait loci results using most recent PD GWAS index variants. Related to** Figure **6**.

**Supplemental Table 8. The American Genome Center group authors**

