## Supplemental data for "The Foundational data initiative for Parkinson’s disease (FOUNDIN-PD): enabling efficient translation from genetic maps to mechanism"

### Supplemental Information

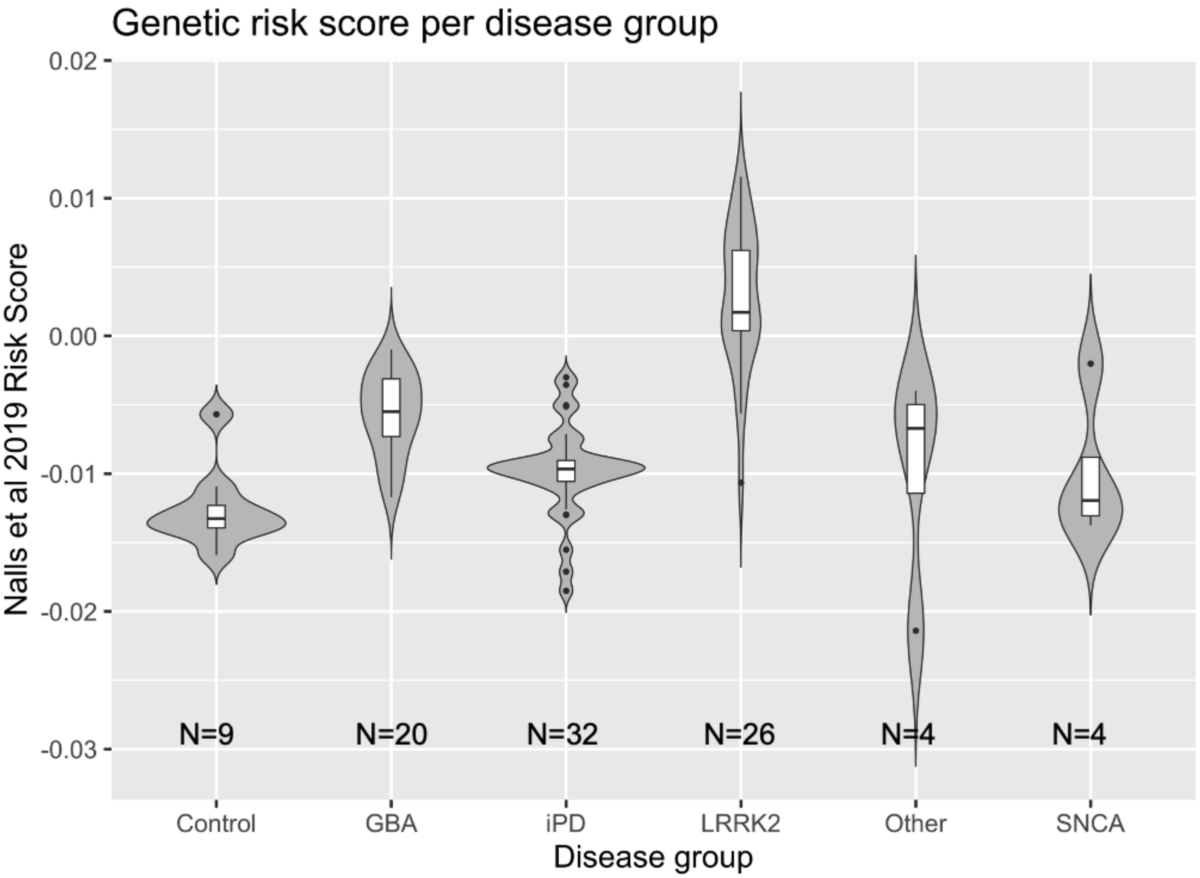


**Supplemental Figure 1.** Parkinson's disease (PD) genetic risk score based on the effect sizes of the most recent PD GWAS (Nalls et al. 2019). Groups are split by genetic status. “Other” includes prodromal (n=3) and SWEDD (n=1). SWEDD = subject with scans without evidence for dopaminergic deficit. Related to Figure 2.

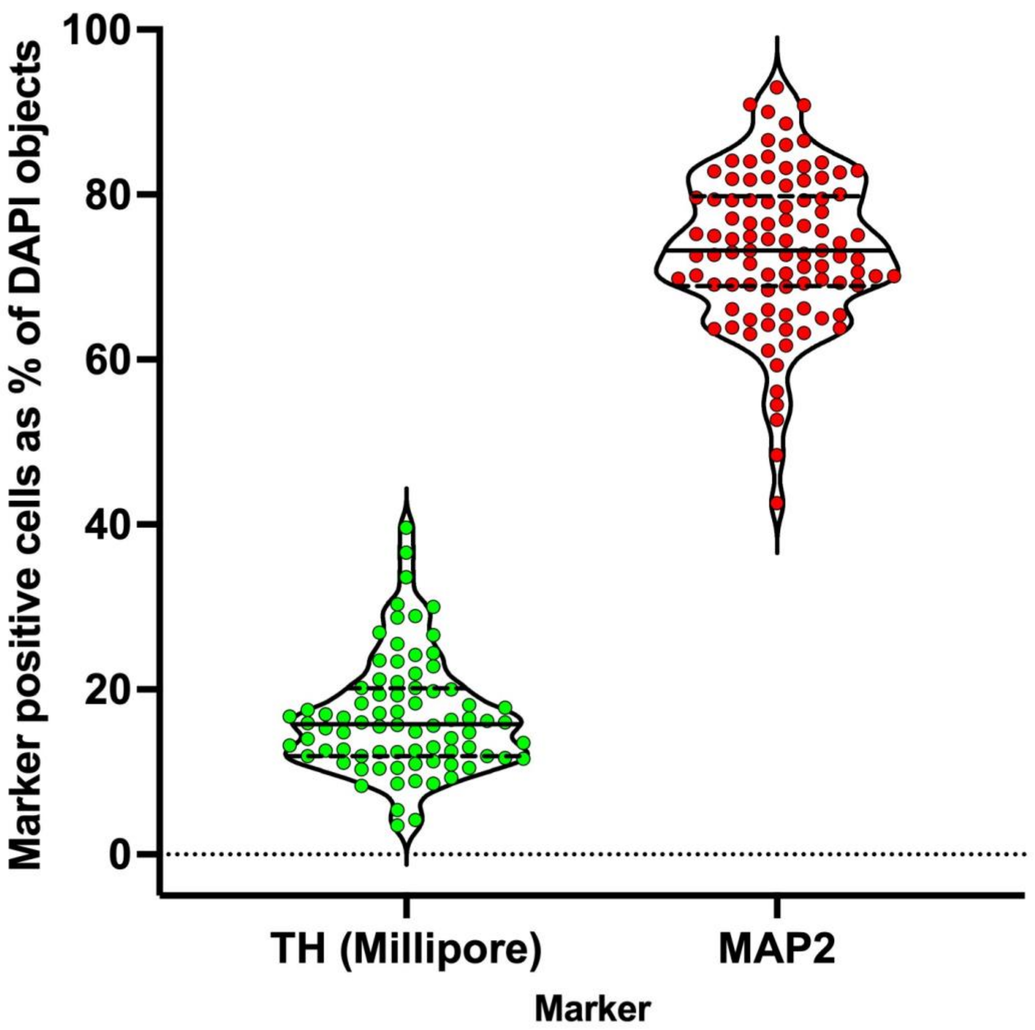


**Supplemental Figure 2.** Percentage of TH (Millipore antibody, green) or MAP2 positive cells (red) at day 65 relative to the total number of cells counted by DAPI positive objects.. Each dot represents the average percentage across 30 fields for one cell line (n=95). Related to Figure 2.

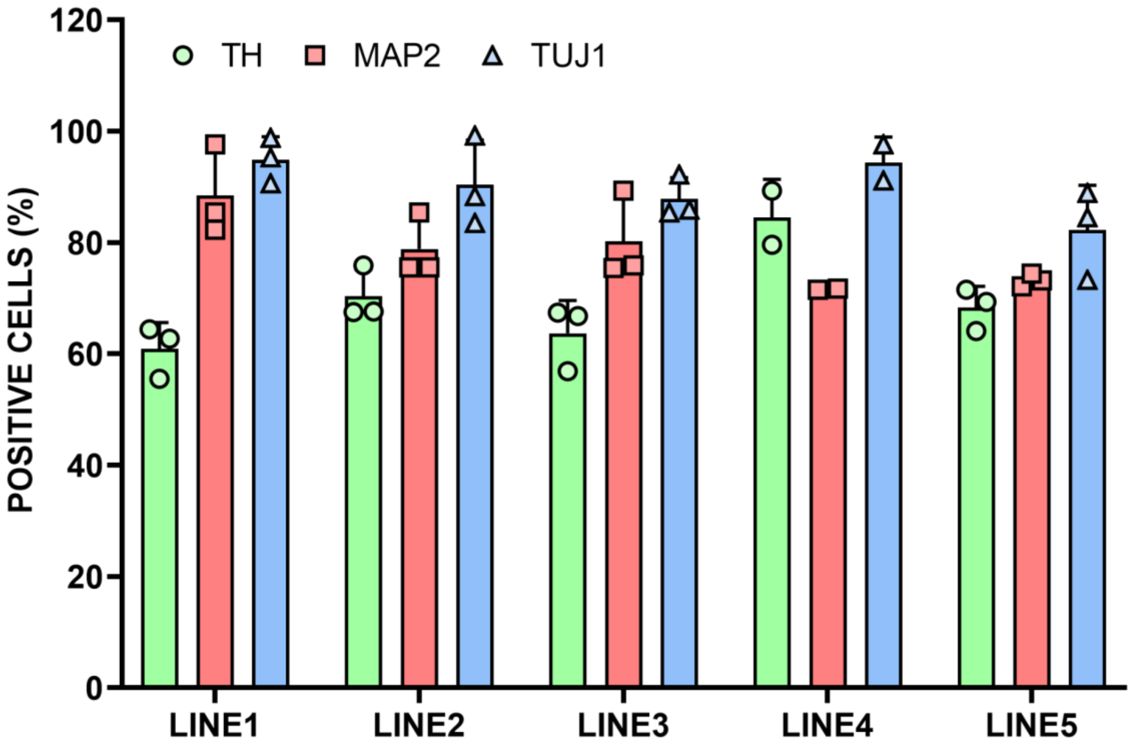


**Supplemental Figure 3.** Percentages of positive cells for TH (Pel-Freeze), MAP2 (Millipore) and TUJ1 (R&D) in five in-house lines on day 65 of differentiation. Each symbol represents one independent differentiation. The barcharts represent the mean±SD of 2-3 differentiations. Denominator is the total number of cells counted. Related to Figure 2.

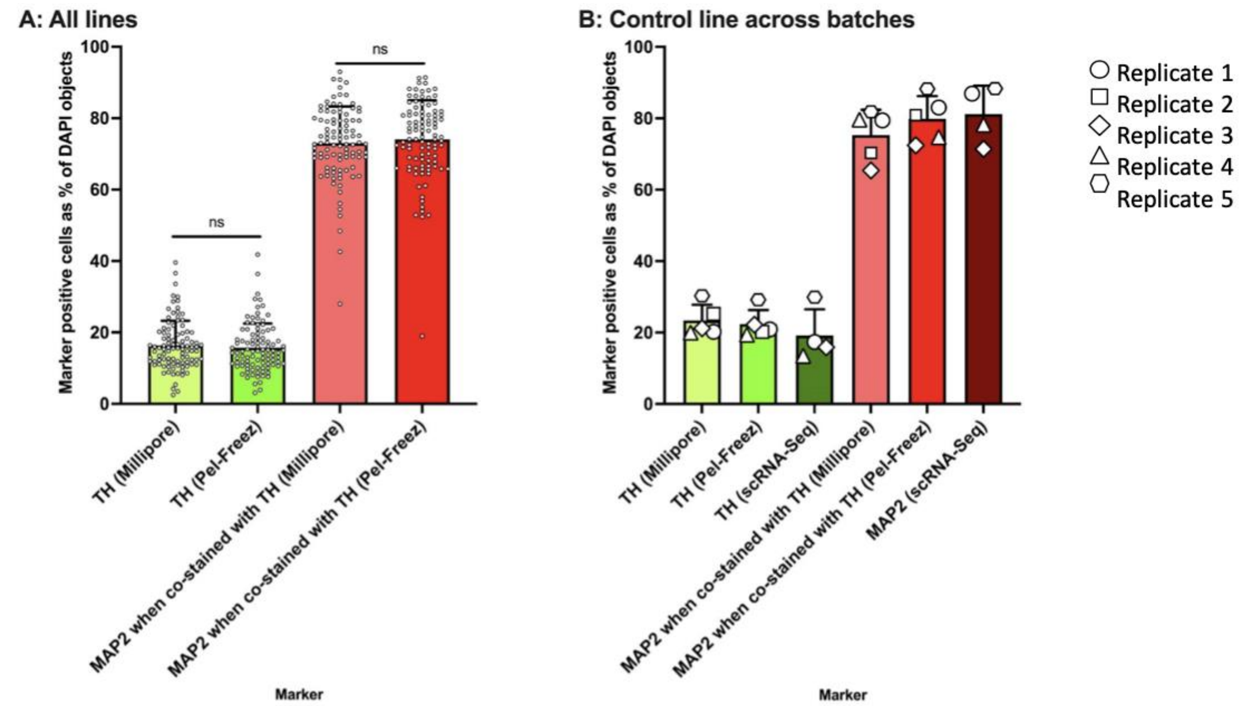


**Supplemental Figure 4.** A, Comparison of the percentages of positive cells stained with MAP2 (Santa Cruz) in combination with either of two independent TH antibodies (M, Millipore; PF, Pel-Freeze) at day 65 of differentiation. TH(M) and TH(PF) represent the percentage of cells showing immunoreactivity to TH relative to the total number cells counted in the cultures, as estimated by DAPI objects. Similarly, the two bars for MAP2 show the percentage of MAP2 positive cells, relative to total cells in the culture, when stained with either TH antibody. The barcharts represent the mean ± SD of 95 lines and individual cell lines are shown with individual dots. ns; not-significant by two-sample unpaired t-tests. B, Percentages of positive cells identified in control line that was used in each batch of automated differentiation for two independent TH antibodies for ICC as well as scRNA-seq data ) as well as MAP2 positive cells by ICC (with each TH antibody) and RNA-Seq. Each symbol represents one replicate of the control cell line. Related to Figure 2.

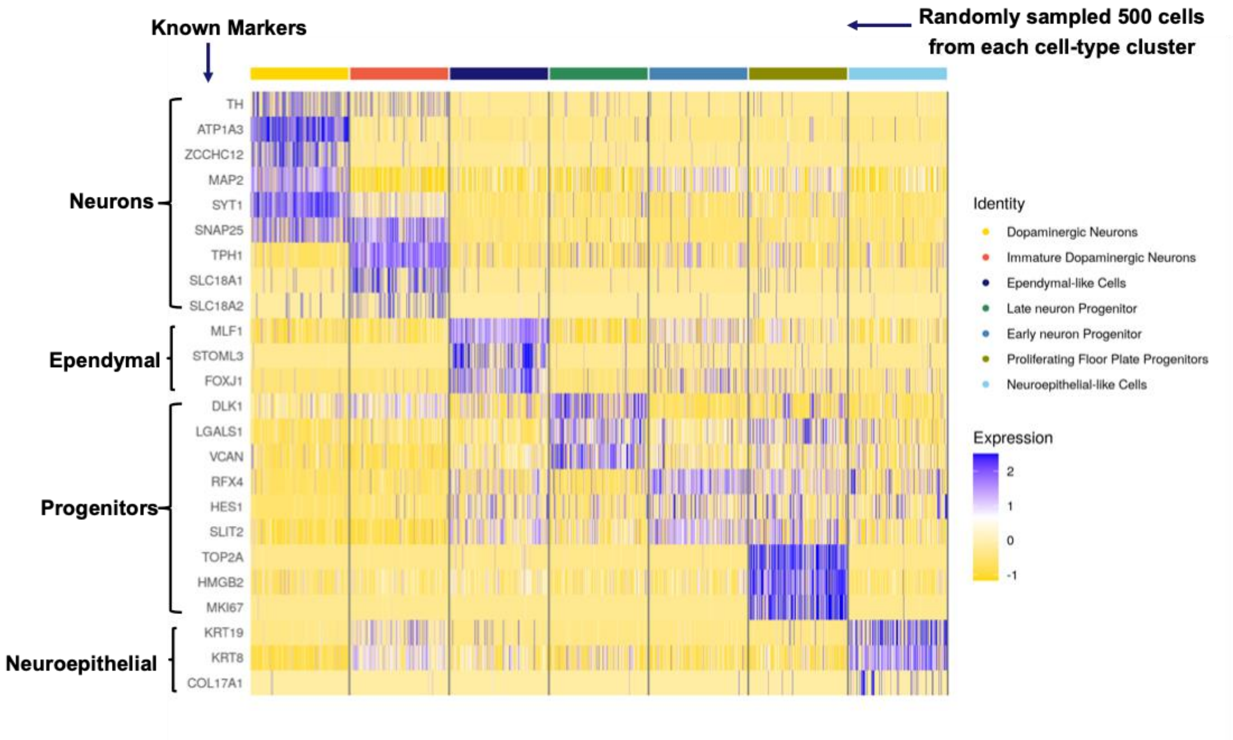


**Supplemental Figure 5.** Heatmap showing expression of marker genes in each single cell RNA-seq cluster that were used for in the cell type assignment of the single cell RNA-seq data. Related to Figure 1 and STAR Methods.

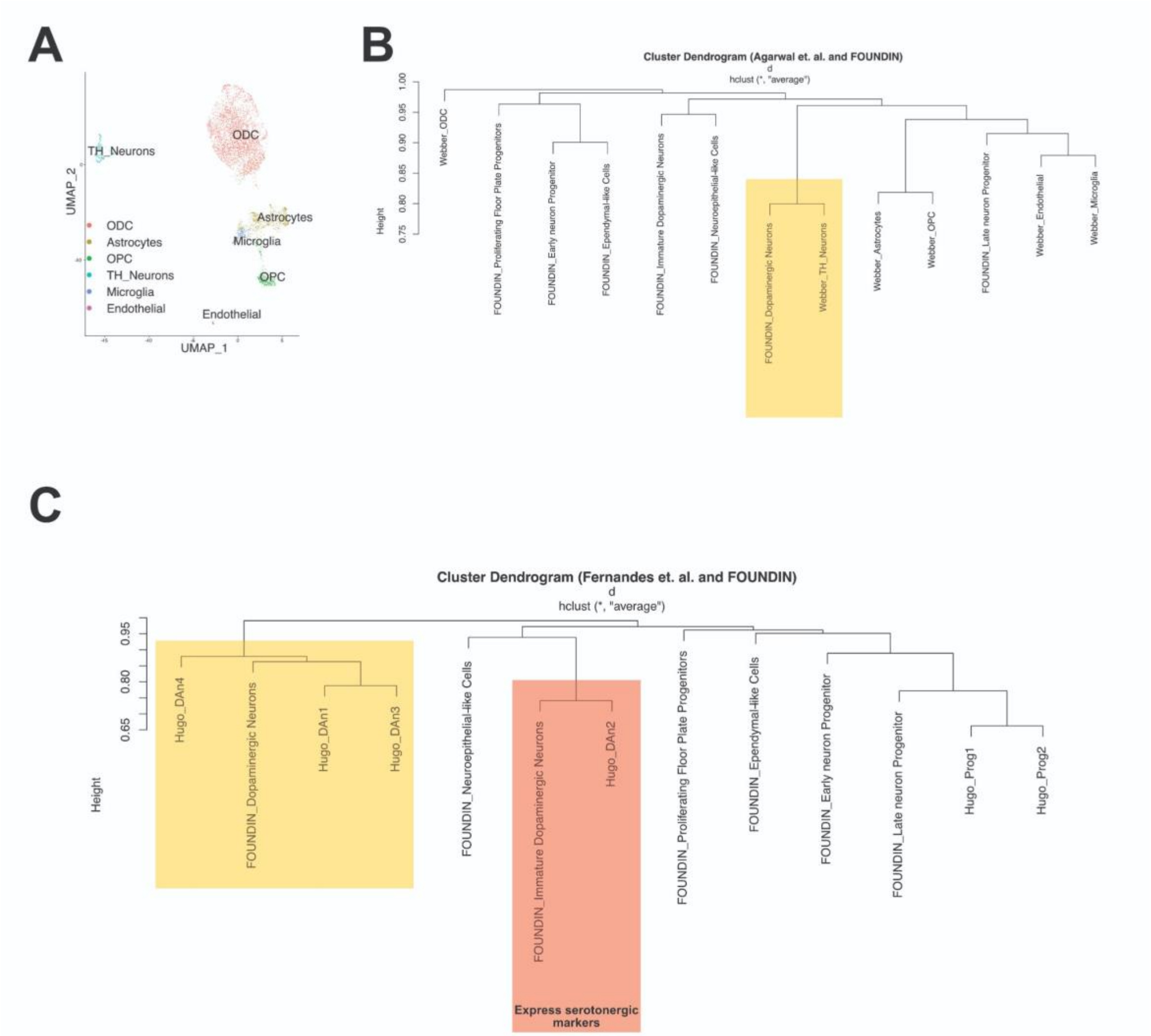


**Supplemental Figure 6.** A, Uniform Manifold Approximation and Projection (UMAP) of cell types identified by Agarwal and collaborators in the post mortem substantia nigra human brain (n=4,781 single cells, n=5+2 replicates). The raw data were analyzed using the FOUNDIN-PD GTF annotation file. 'TH_Neurons' were annotated because this cluster showed expressionup-regulation of *TH*. ODCs, oligodendrocytes; OPCs, oligodendrocyte precursor cells. B, Dendrogram showing clustering of FOUNDIN-PD cell types with Agarwal and collaborators' dataset (Agarwal et al. 2020) using ClusterMap (Gao et al. 2019). C, Dendrogram showing clustering of FOUNDIN-PD cell types with Fernandes and collaborators dataset (Fernandes et al. 2020) using the ClusterMap tool. Related to Figure 2.

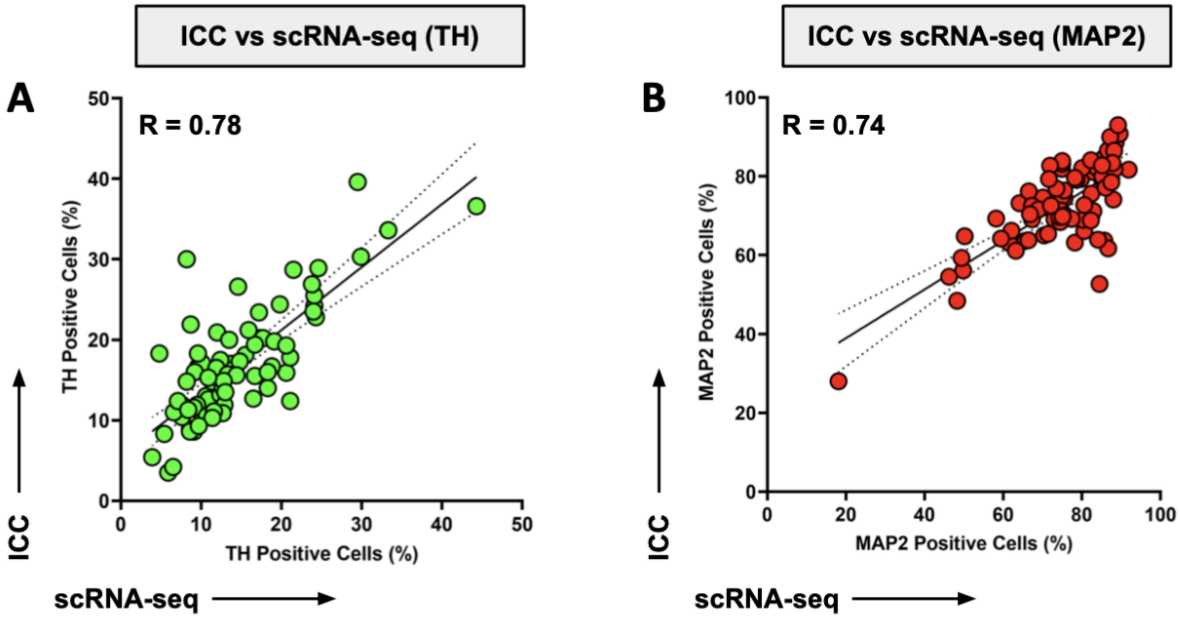


**Supplemental Figure 7.** A, Correlation between percentages of TH (Millipore) and B, MAP2 positive cells in ICC and scRNA-seq (R, Pearson correlation coefficient; p<0.0001). Each dot represents one cell line (n=83). Related to Figure 2.

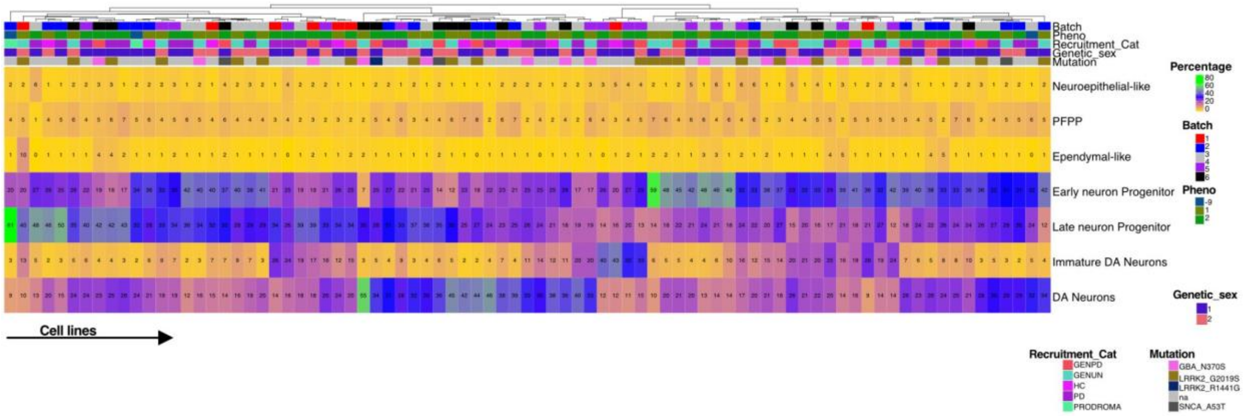


**Supplemental Figure 8.** Variability in differentiation efficiency across the iPSC lines depicted as Heatmap plot. The number shows the percentage of cell type for a particular sample and clustering was performed using default settings in ComplexHeatmap (Gu, Eils, and Schlesner 2016). For each sample, annoations are shown at the top depicting batch, phenotype, recruitment category, genetic sex and PD-linked genotype (GBA1+, LRRK2+, SNCA+) information. Recruitment_Cat = is the recruitment strategy used by PPMI, GENPD = genetic PD affected, GENUN = genetic PD unaffected, HC = healthy control, PD = Parkinson's disease case, PRODROMA = prodromal case. Since PRODROMA are not real PD cases phenotype -9 is used for this group. Related to Figure 2.

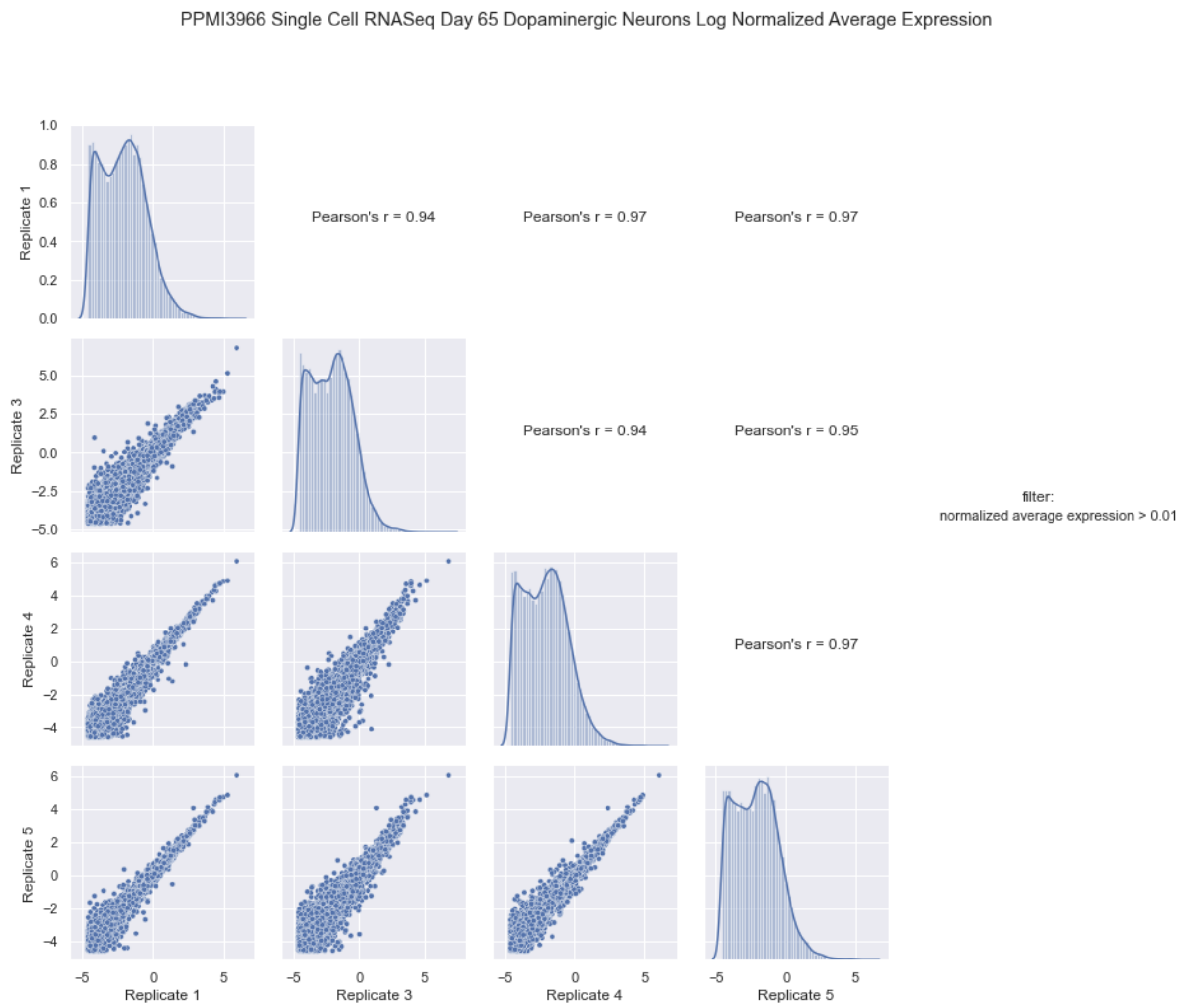


**Supplemental Figure 9.** Gene level expression correlation between four technical replicates of the control cell line using scRNA-seq dopaminergic neuron data. Only genes with average expression greater than 0.01 transcript per million were included. Average expression values were log normalized and compared with other replicates in scatter plots in the lower triangle. Distribution of log normalized values is shown on the diagonal. Pearson's r correlation is displayed in the upper triangle. Related to Figure 2.

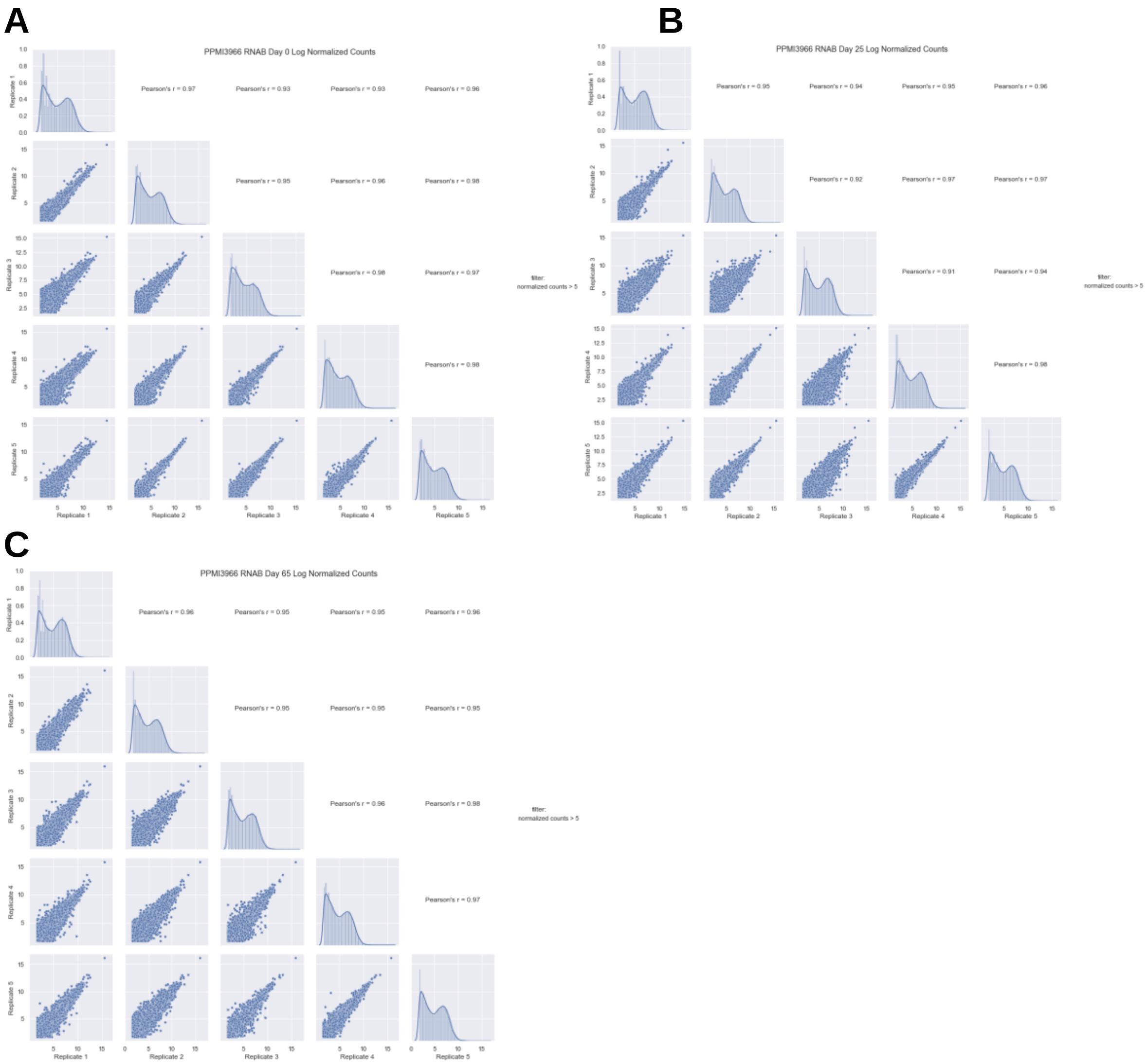


**Supplemental Figure 10.** Gene level expression correlation between five technical replicates of the control cell line using bulk RNA-seq data. A, timepoint day 0; B, timepoint day 25; C, timepoint day 65. Only genes with normalized counts greater than five were included. Counts were log normalized and compared with other replicates in scatter plots in the lower triangle. Distribution of log normalized values is shown on the diagonal. Pearson’s r correlation is displayed in the upper triangle. Related to Figure 3.

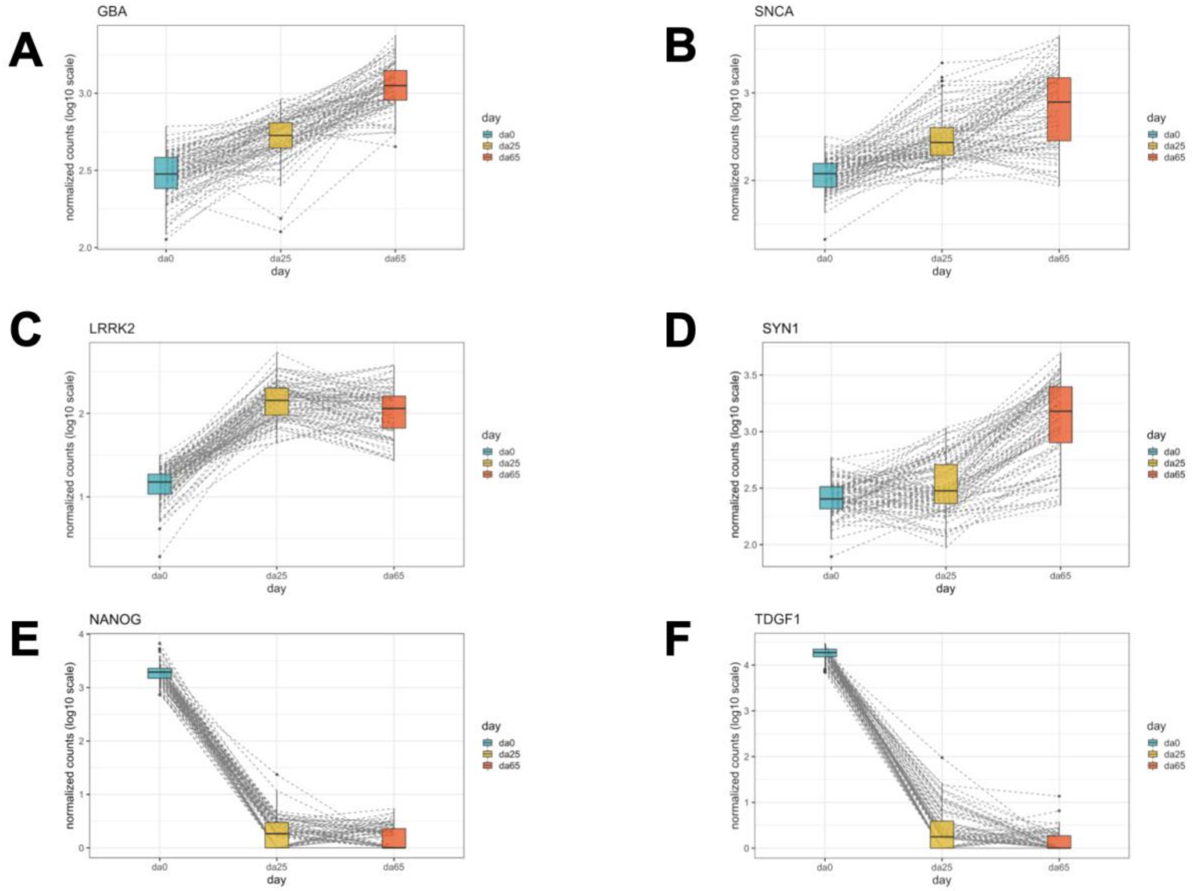


**Supplemental Figure 11.** A-D, Expression of monogenic PD genes (*GBA1*, *SNCA*, *LRRK2)* and neuronal markers (SYN1) goes up across timepoints in bulk RNA-seq. E-F, Stem cell marker gene (*NANOG* and *TDGF1)* expression goes down across timepoints. Related to Figure 3.

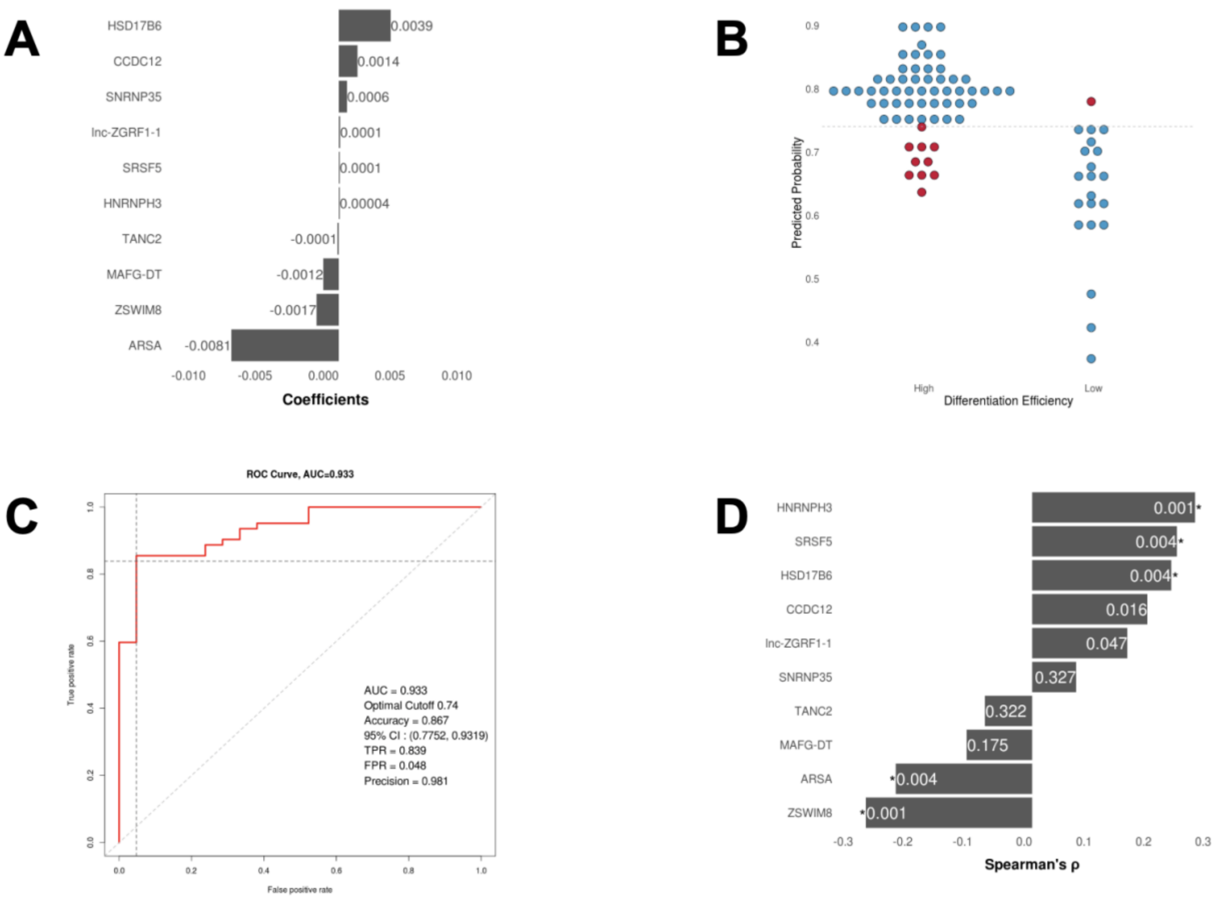


**Supplemental Figure 12.** A, Ten genes identified with non-zero coefficients using elastic net regularization approach (see Methods) to predict neuronal differentiation efficiency. B, Confusion matrix results are shown as binary scatter plots. Predicted probability is on the y-axis and initially assigned labels are on the x-axis (see Methods). Each dot represents a cell line. Blue and red color denotes truly and falsely predicted labels by the trained model. Dashed line represents the optimal probability cut-off. C, Area under the curve (AUC) using ten genes and all samples for training. D, Correlation analysis of these ten genes with neuronal differentiation efficiency. Numbers in the bar are adjusted p-values. * represents significant associations (FDR < 1%). Related to Figure 3.

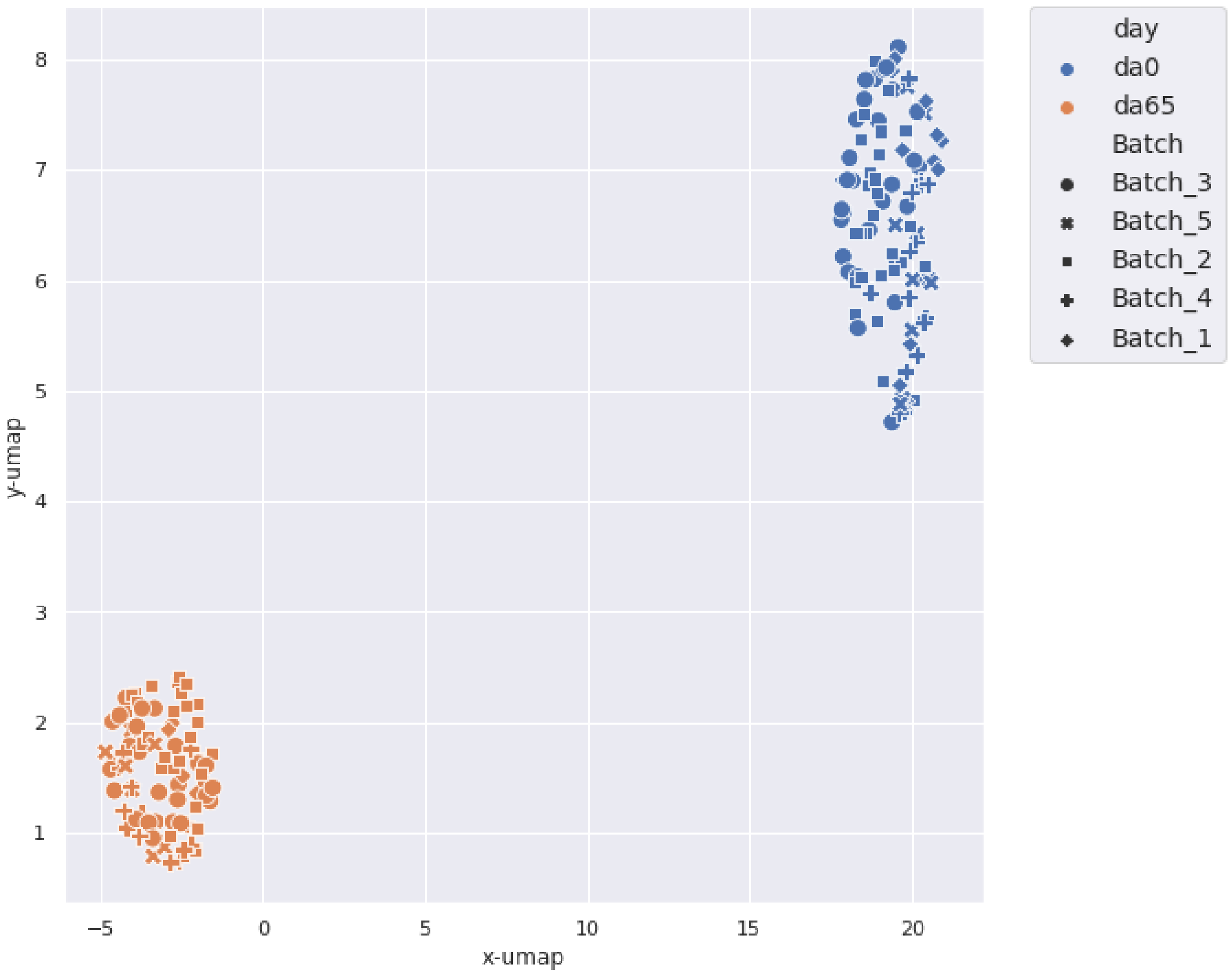


**Supplemental Figure 13.** Uniform Manifold Approximation and Projection (UMAP) analysis of methylation data shows clear clustering by timepoints (day 0 vs day 65). Related to Figure 4.

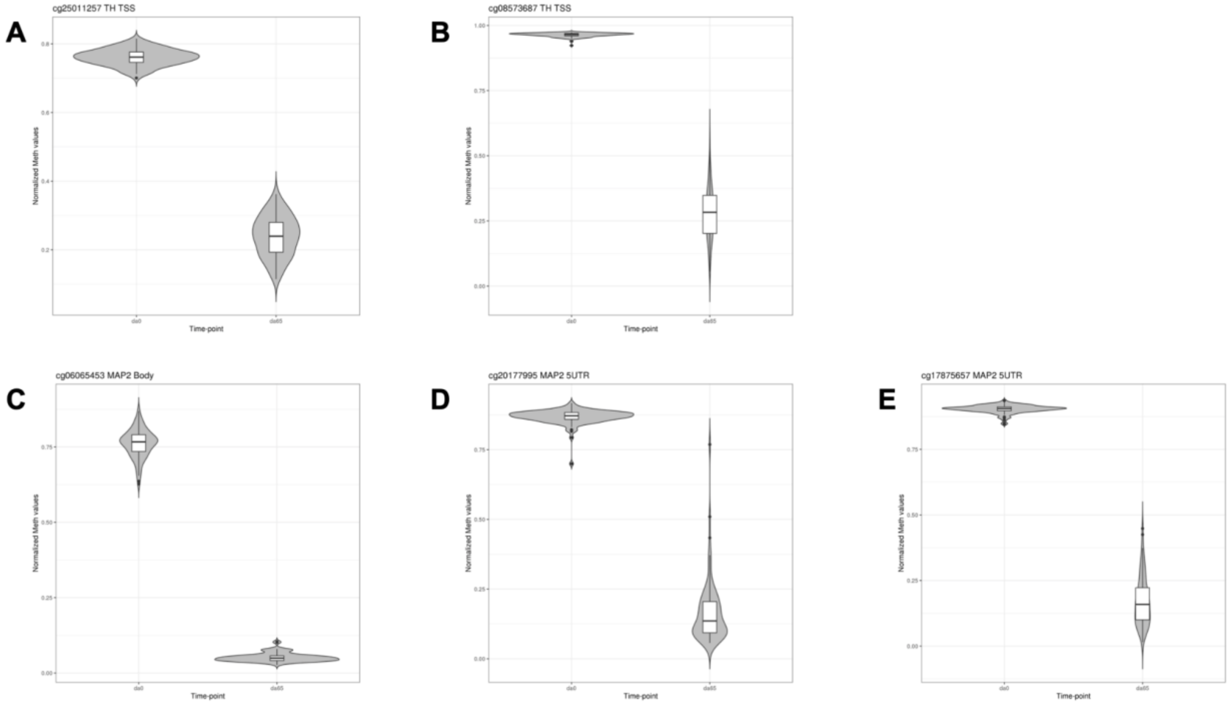


**Supplemental Figure 14.** A-B, Methylation probes in close proximity to the *TH* transcription start sites show a clear decrease in methylation levels between timepoints (day 0 vs day 65). C-E, Methylation probes across the *MAP2* (gene body, 5' UTR and 3' UTR) locus show a decrease in methylation levels between timepoints. Related to Figure 4.

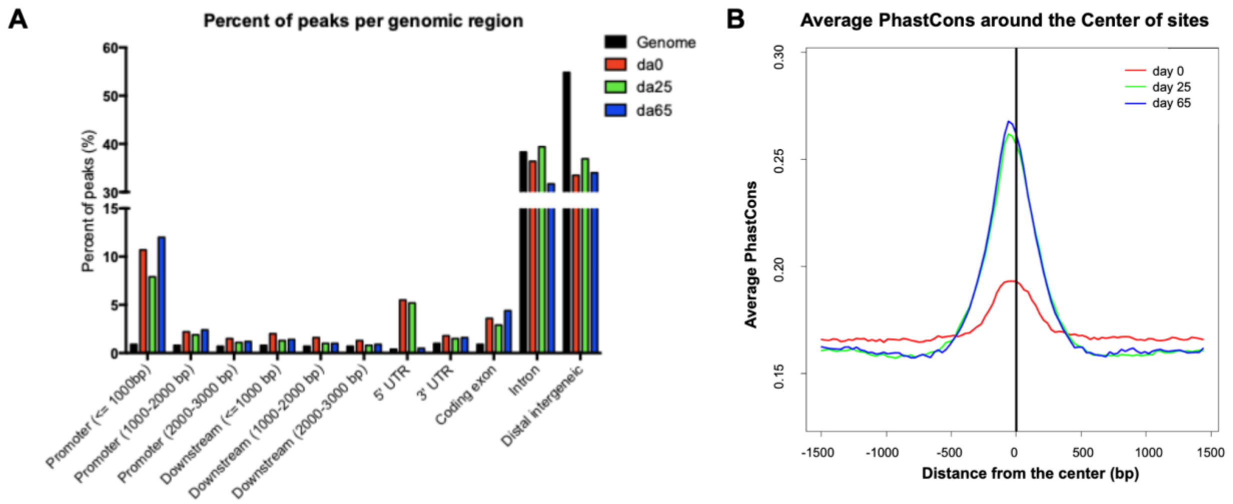


**Supplemental Figure 15.** A, Cistrome analysis showing the percent of bulk ATAC-seq peaks across genomic regions in a merged peak set at each timepoint (day 0 = red, day 25 = green, day 65 = blue) relative to the background genomic space (black). B, The average genetic conservation across species, PhastCons score, across merged ATAC-seq peak sets as cells are differentiated (day 0 = red, day 25 = green, day 65 = blue). Related to Figure 4.

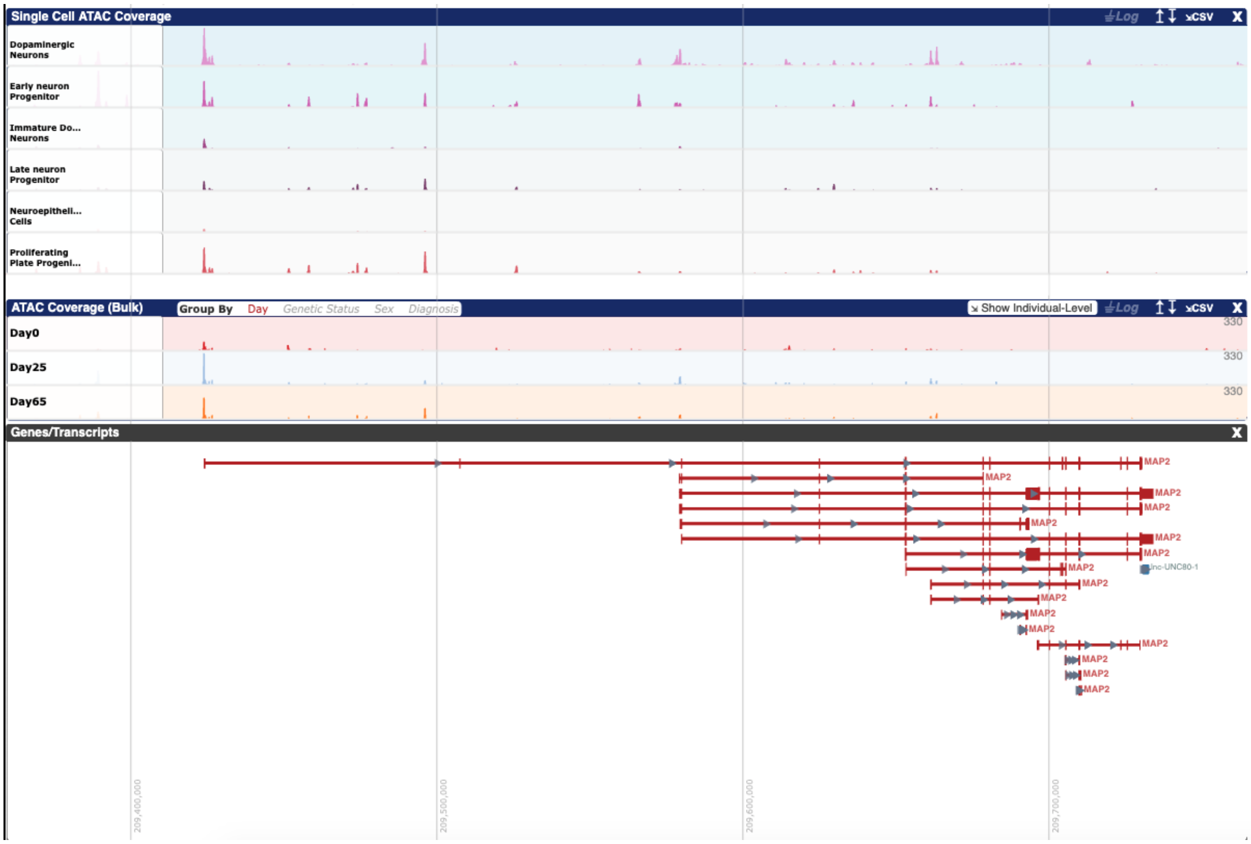


**Supplemental Figure 16.** Open chromatin at the *MAP2* locus shows differential peaks across timepoints (bulk ATAC-seq) and cell types (scATAC-seq). Related to Figure 4.

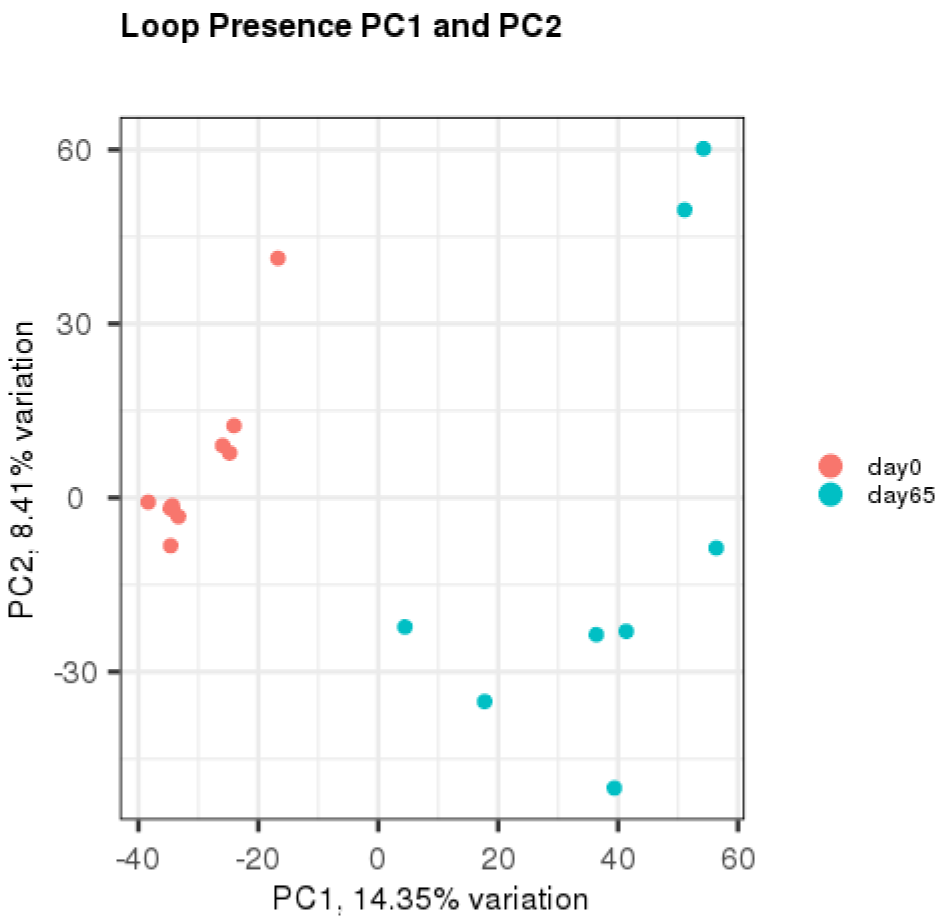


**Supplemental Figure 17.** Principal component analysis (PCA) of HiC-seq data shows clustering by timepoints (day 0 vs day 65). Related to Figure 4.

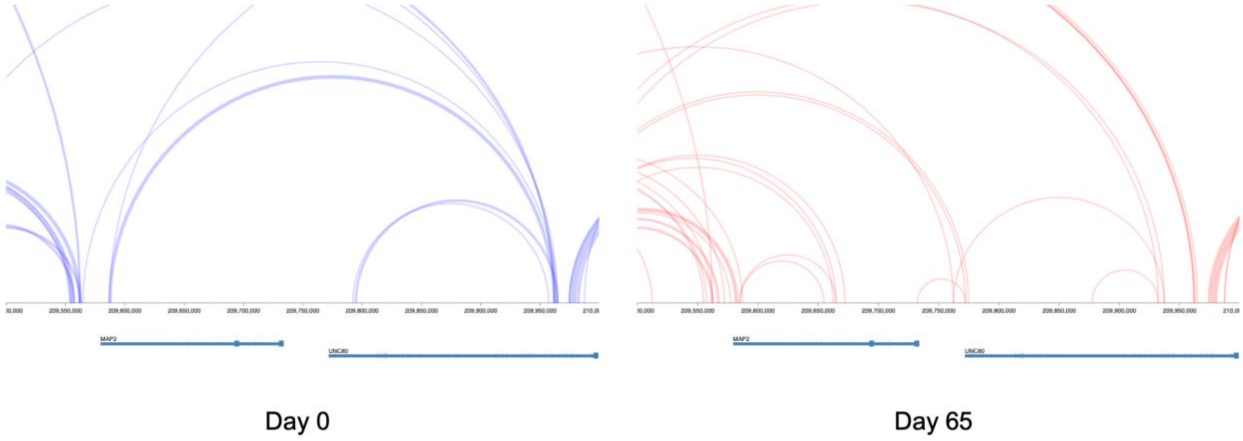


**Supplemental Figure 18.** HiC loop structure at the *MAP2* locus shows clear differences between timepoints (day 0 vs day 65). Related to Figure 4.

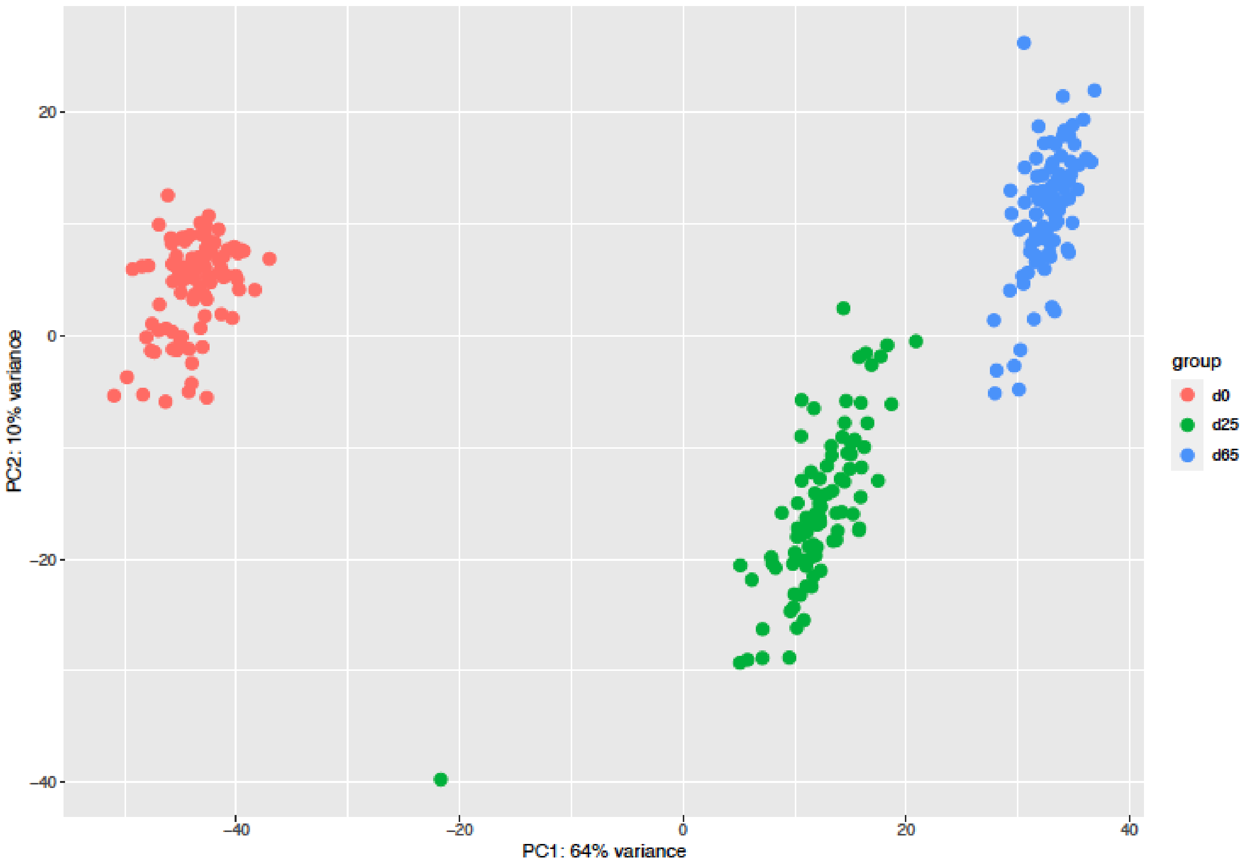


**Supplemental Figure 19.** Principal component analysis (PCA) of small RNA-seq data shows clear clustering of expression by timepoint (day 0 vs day 25 vs day 65). Related to Figure 4.

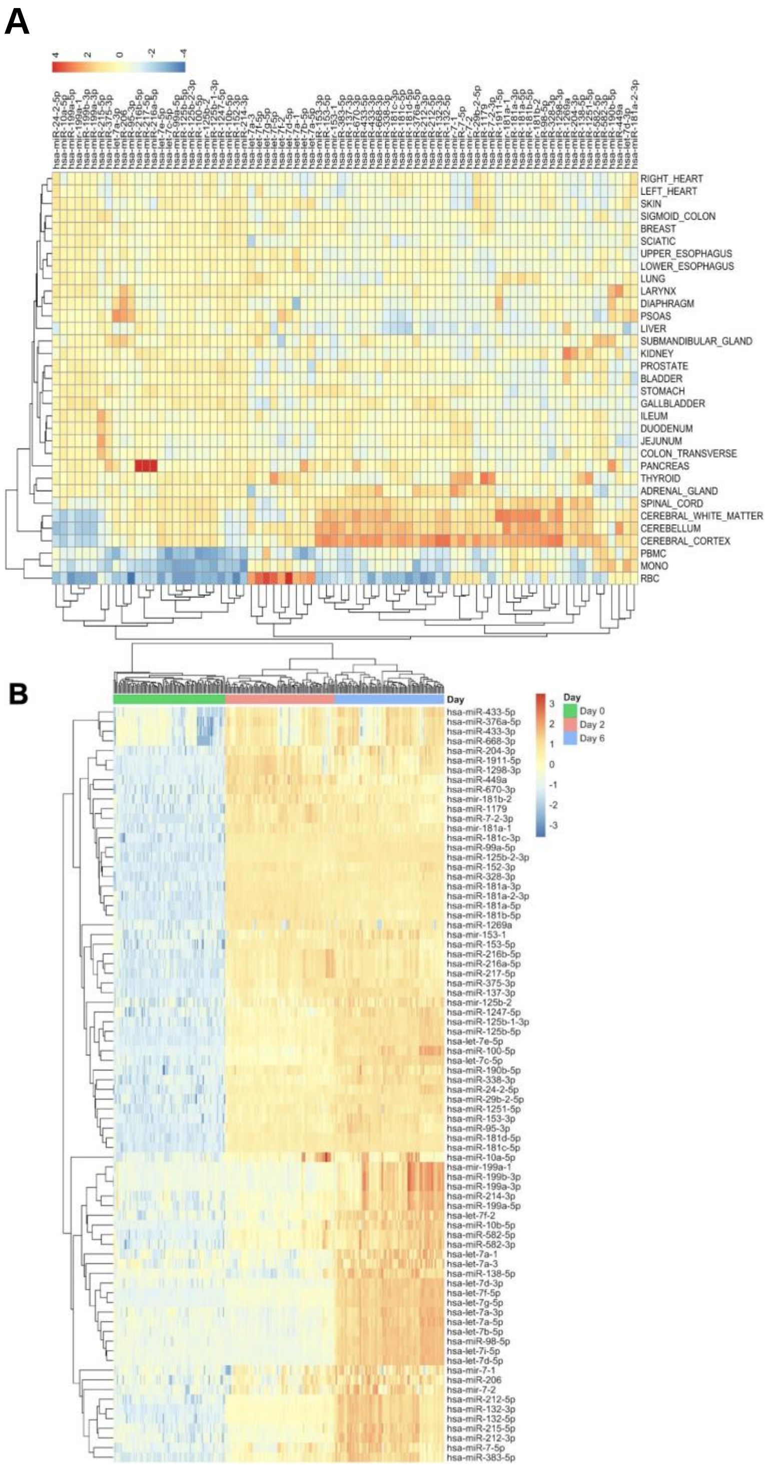


**Supplemental Figure 20.** Heatmap of upregulated miRNA expression between Day 0 and day 65. The expression of the miRNAs that were upregulated are displayed across 34 different tissues. It can be noted that several miRNAs are well-expressed in CNS tissues. Color scale is the Z score of the expression. Related to Figure 4.

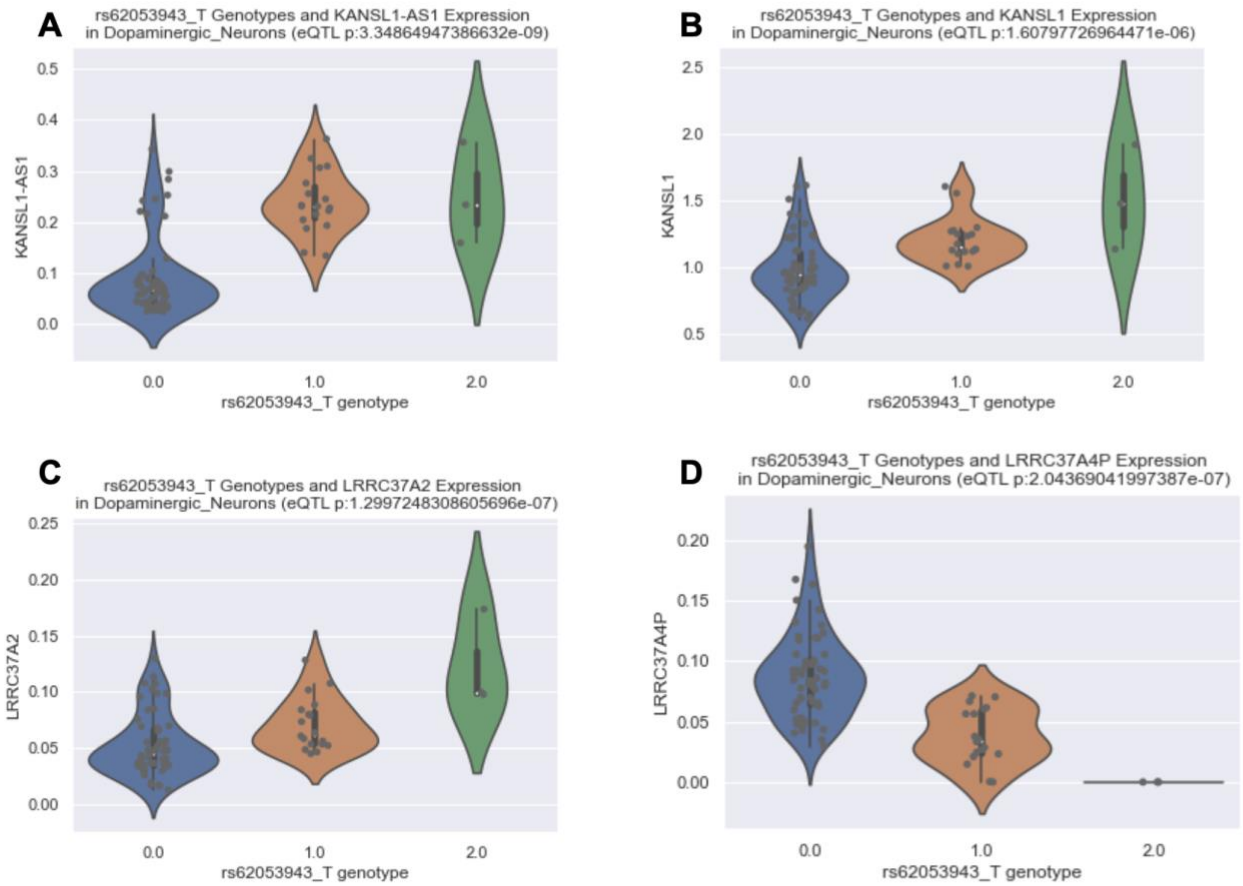


**Supplemental Figure 21.** Expression quantitative trait loci (eQTL) results of rs62053943 and genes expressed at the MAPT region (17q21-31) showing clear eQTLs likely representing the H1/H2 MAPT haplotype status. Related to Figure 6.

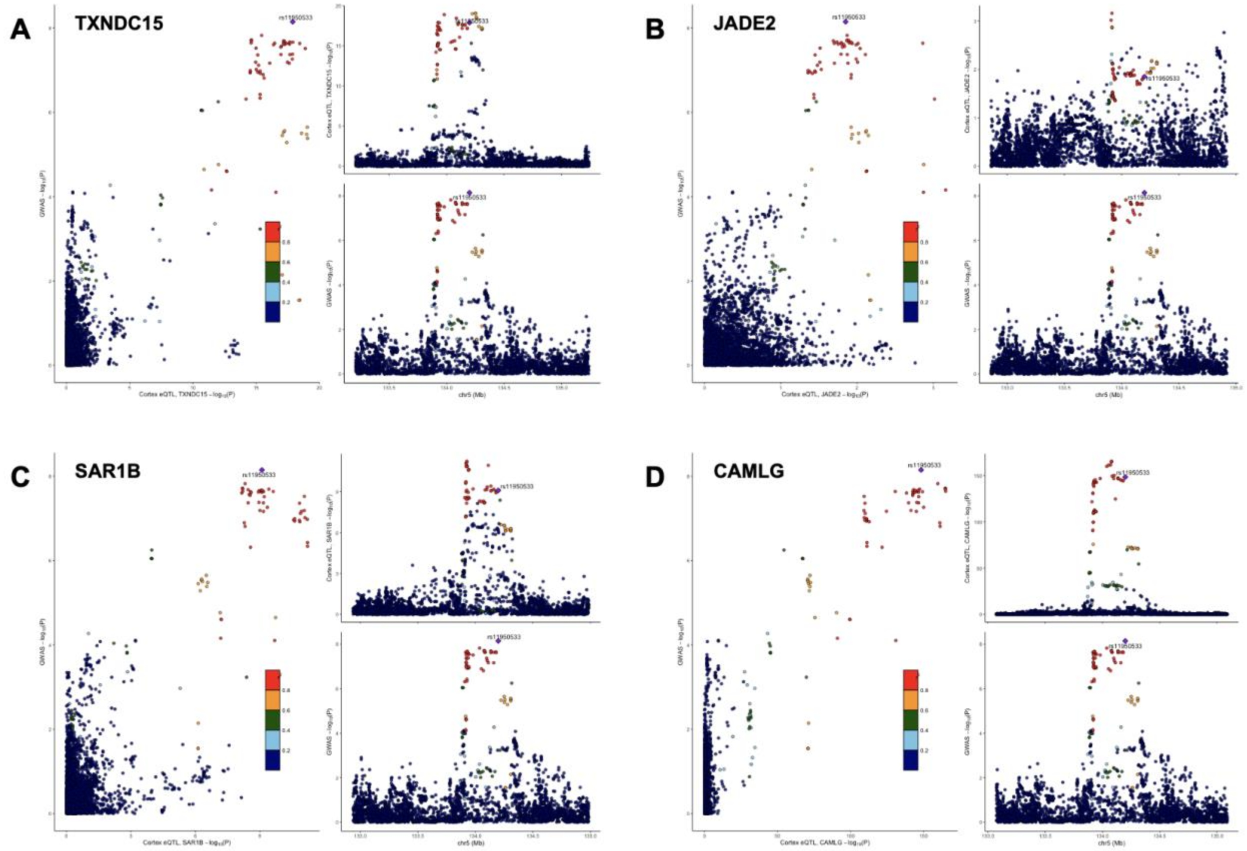


**Supplemental Figure 22.** LocusCompare plots of the correlation between the most recent PD GWAS association results (Nalls et al. 2019) and cortical brain eQTL data (Sieberts et al. 2020). A-D, TXNDC15, JADE2, SAR1B and CAMLG. Related to Figure 6.

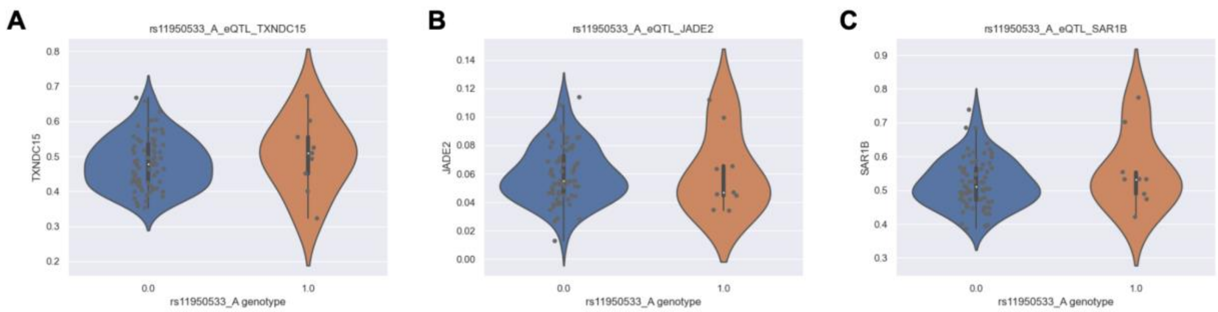


**Supplemental Figure 23.** Absence of significant expression quantitative trait loci for A, TXNDC15; B, JADE2 and C, SAR1B in DA neuron scRNA-seq data. Related to Figure 6.

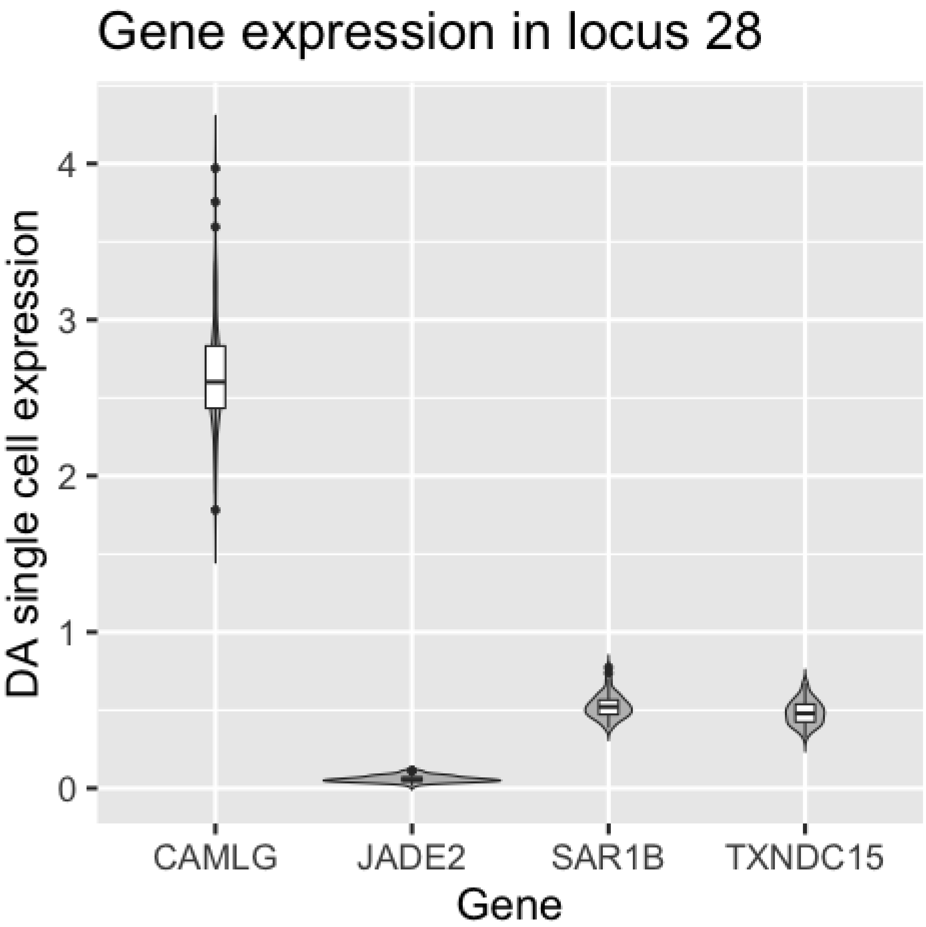


**Supplemental Figure 24.** Overview of scRNA-seq data gene level expression in the DA neuron cell cluster at the selected PD GWAS locus. Related to Figure 6.

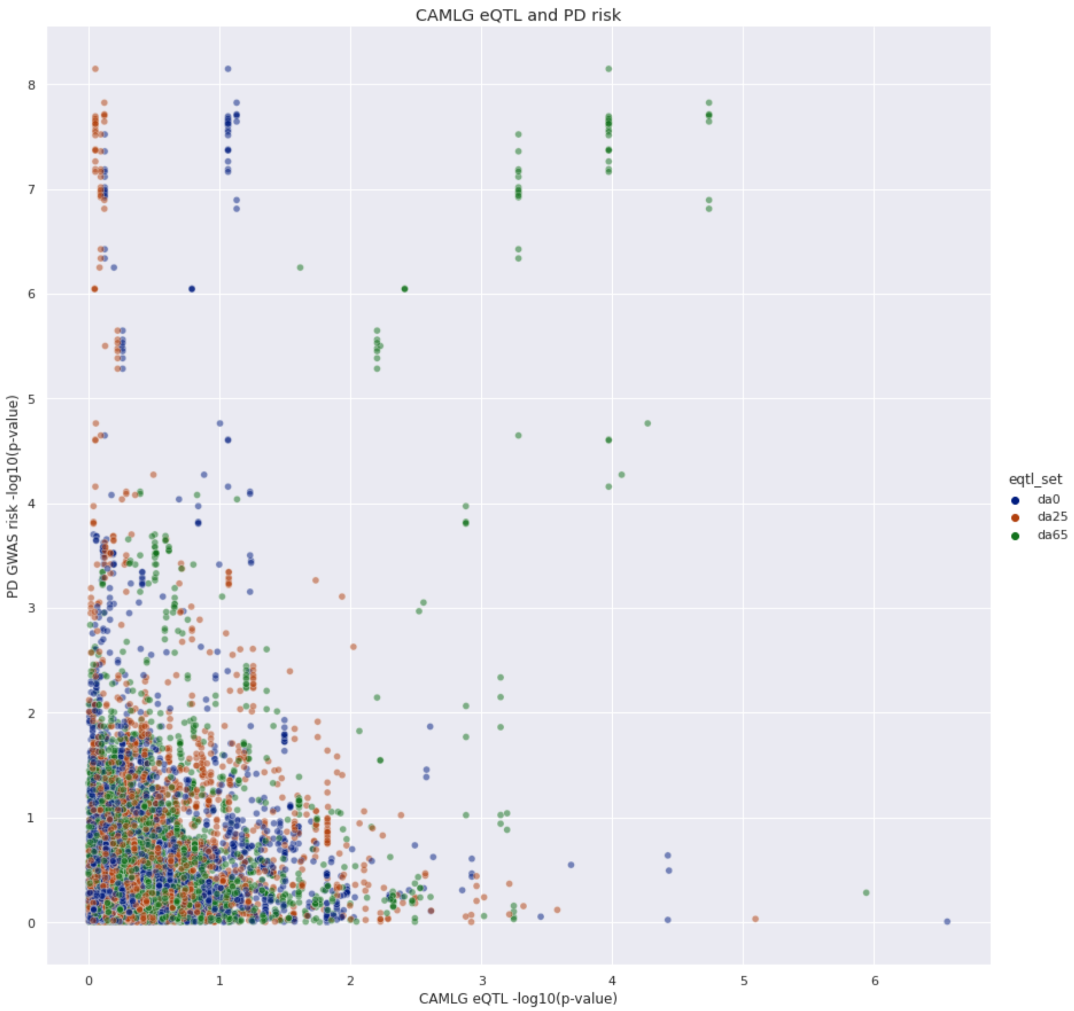


**Supplemental Figure 25.** Intersection of the CAMLG bulk RNA-seq eQTL signal and the PD risk signal at day 0 (blue), day 25 (red) and day 65 (green). Related to Figure 6.

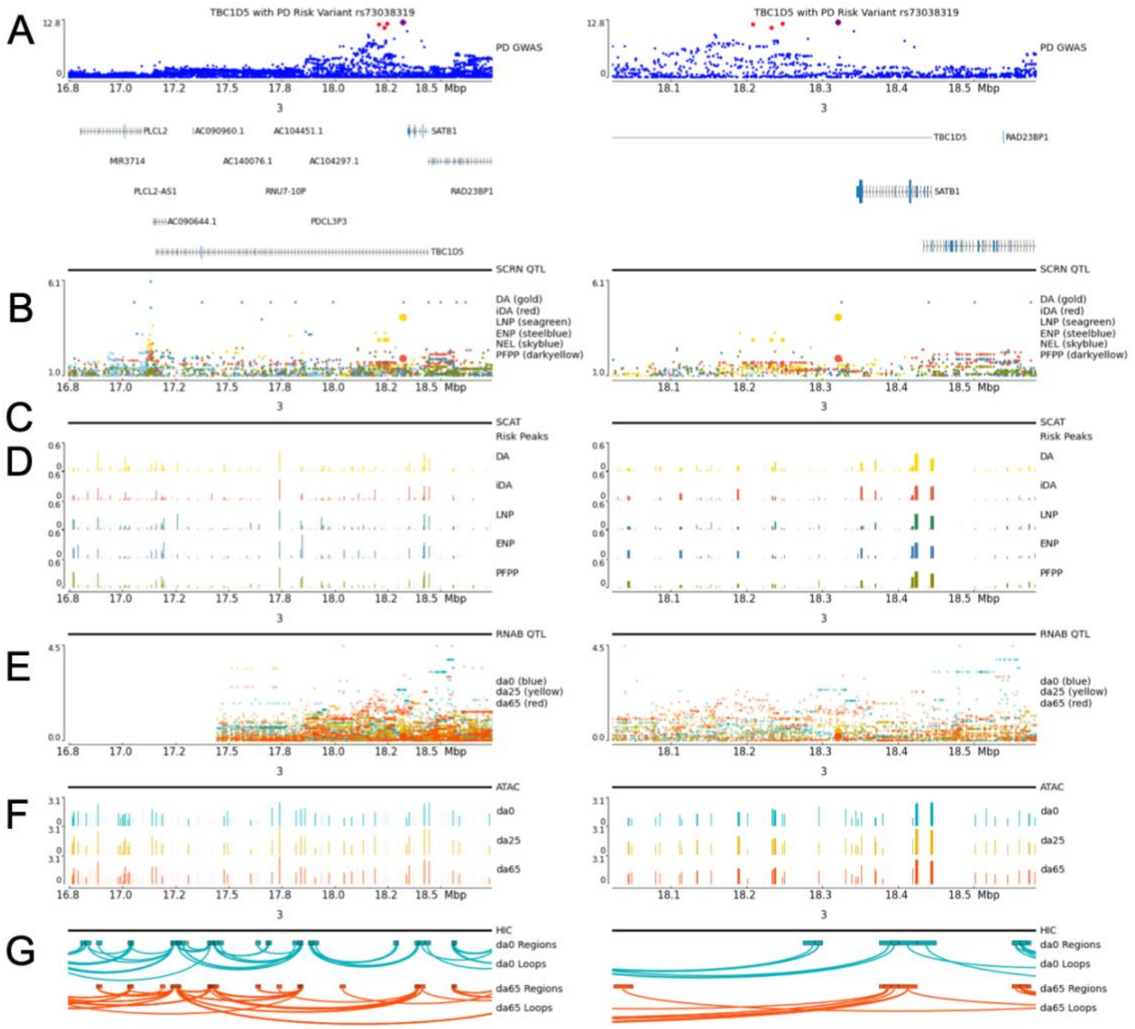


**Supplemental Figure 26.** PD risk locus multi omics figure near TBC1D5 on chromosome 3. PD risk locus multi omics figure near TBC1D5 on chromosome 3. Tracks represent different data modalities generated or considered in FOUNDIN-PD as different data tracks; figures generated using pyGenomeTracks. The left and right panels of the figures display the same tracks where the left side is a larger region centered on the PD risk locus and the ride side only includes the interval containing the index PD risk variant for this locus and variants in LD with that index variant. (A) GWAS risk for PD in the region. Point size denotes $r^2$ linkage disequilibrium with PD index variant rs73038319 (large: $r^2$=1, medium: 1 > r2 >= 0.8, small: $r^2$ < 0.8). (B) Single-cell RNA-seq (SCRN) eQTL data for dopaminergic neurons (DA), immature dopaminergic neurons (iDA), late neuron progenitors (LNP), early neuron progenitors (ENP), neuroepithelial-like cells (NEL) and proliferating floor plate progenitors (PFPP). (C) Single-cell ATAC-seq (SCAT) peaks containing a variant in high ld ($r^2$ >= 0.8) with rs73038319. (D). Single-cell ATAC-seq peaks for different cell types. (E) Bulk RNA-seq (RNAB) eQTL results per differentiation timepoint for *TBC1D5*. (F) Bulk ATAC-seq peaks separated per differentiation timepoint. (G) HiC data depicting chromatin regions connected by loops at different differentiation timepoints. Related to Figure 7.

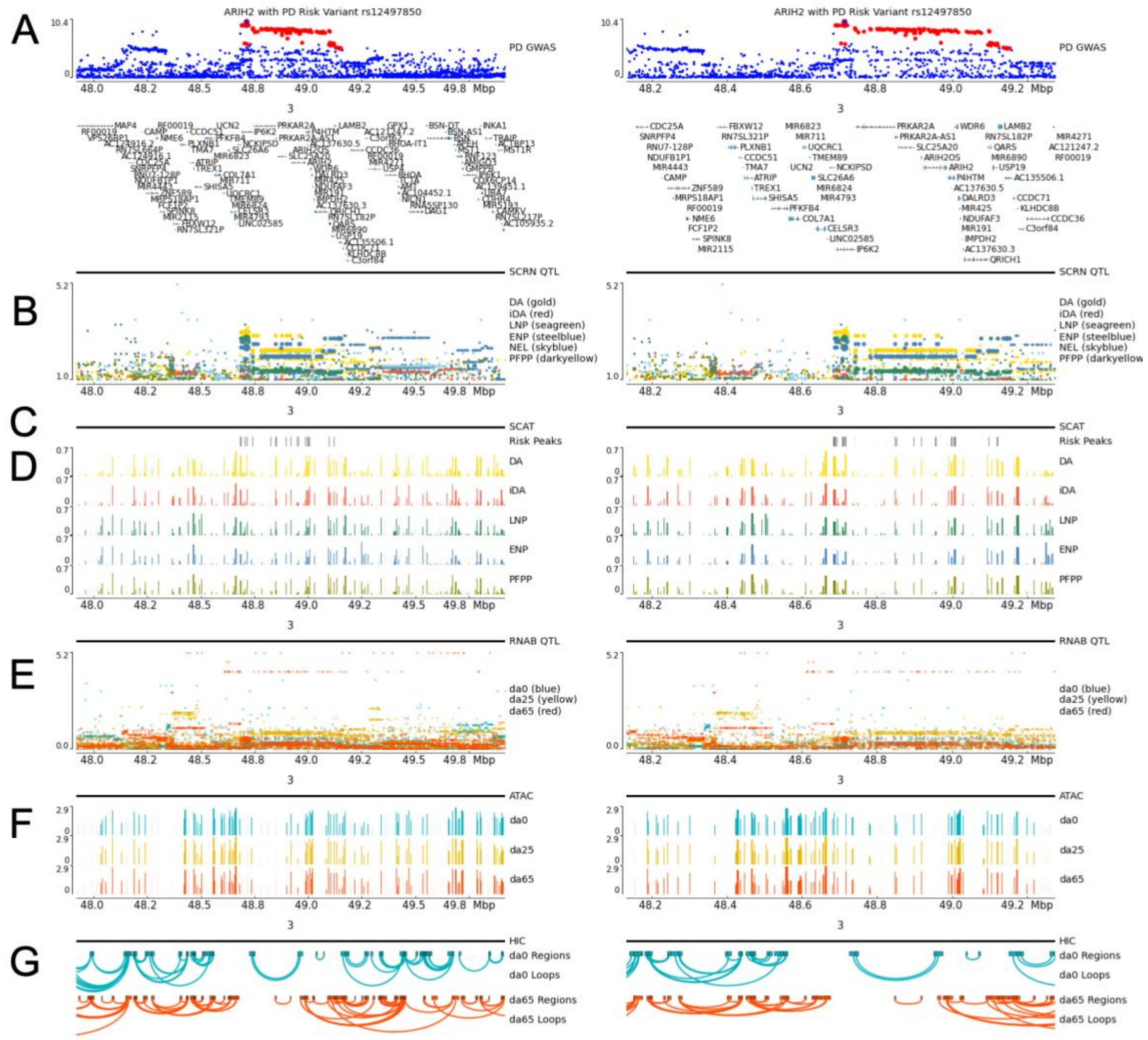


**Supplemental Figure 27.** PD risk locus multi omics figure near ARIH2 on chromosome 3. Tracks represent different data modalities generated or considered in FOUNDIN-PD as different data tracks; figures generated using pyGenomeTracks. The left and right panels of the figures display the same tracks where the left side is a larger region centered on the PD risk locus and the ride side only includes the interval containing the index PD risk variant for this locus and variants in LD with that index variant. (A) GWAS risk for PD in the region. Point size denotes r2 linkage disequilibrium with PD index variant rs12497850 (large: r2=1, medium: 1 > r2 >= 0.8, small: r2 < 0.8). (B) Single-cell RNA-seq (SCRN) eQTL data for dopaminergic neurons (DA), immature dopaminergic neurons (iDA), late neuron progenitors (LNP), early neuron progenitors (ENP), neuroepithelial-like cells (NEL) and proliferating floor plate progenitors (PFPP). (C) Single-cell ATAC-seq (SCAT) peaks containing a variant in high ld (r2 >= 0.8) with rs12497850. (D). Single-cell ATAC-seq (SCAT) peaks for different cell types. (E) Bulk RNA-seq (RNAB) eQTL results per differentiation timepoint for ARIH2. (F) Bulk ATAC-seq peaks separated per differentiation timepoint. (G) HiC data depicting chromatin regions connected by loops at different differentiation timepoints. Related to Figure 7.

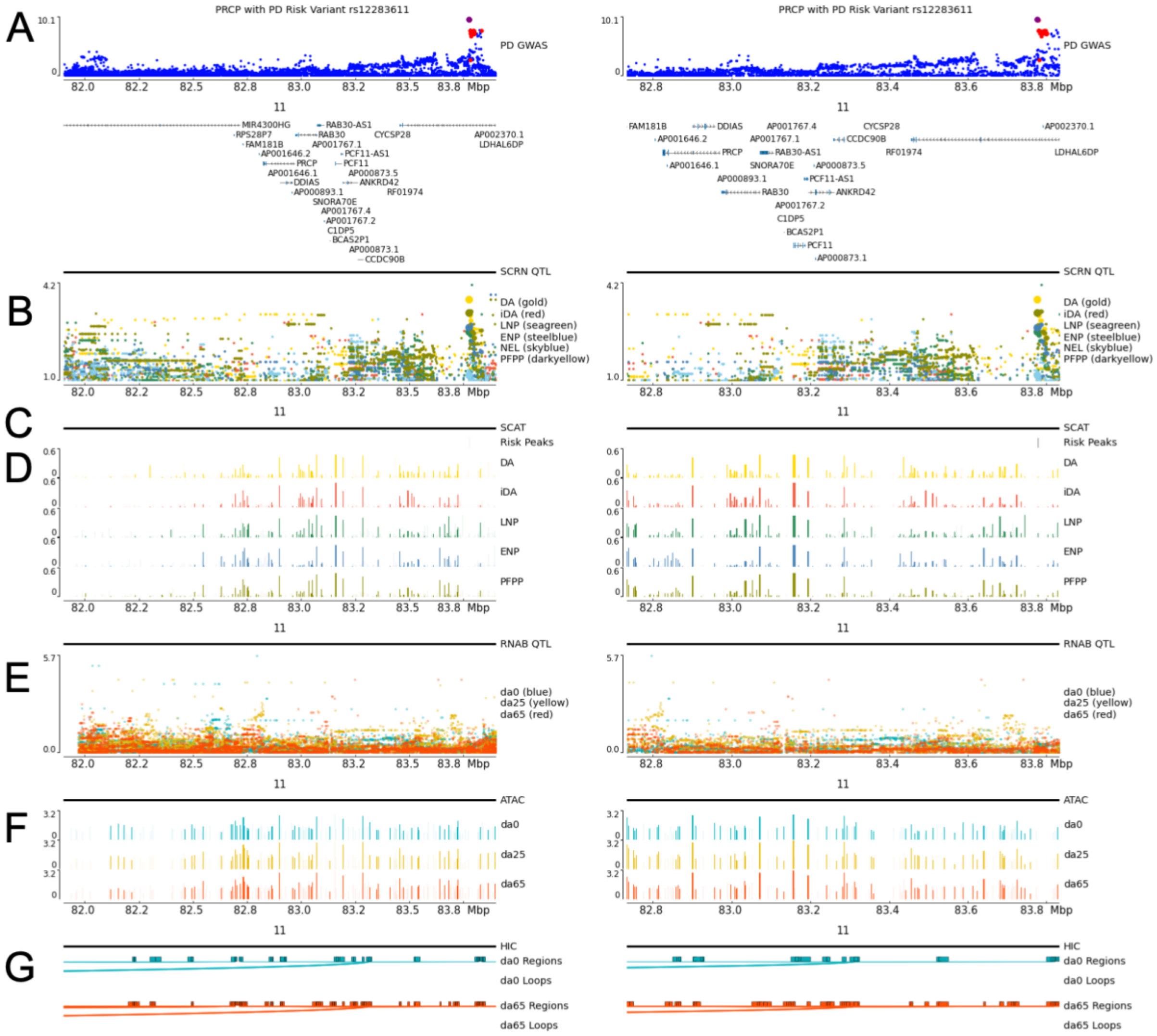


**Supplemental Figure 28.** PD risk locus multi omics figure near PRCP on chromosome 11. Tracks represent different data modalities generated or considered in FOUNDIN-PD as different data tracks; figures generated using pyGenomeTracks. The left and right panels of the figures display the same tracks where the left side is a larger region centered on the PD risk locus and the ride side only includes the interval containing the index PD risk variant for this locus and variants in LD with that index variant. (A) GWAS risk for PD in the region. Point size denotes r2 linkage disequilibrium with PD index variant rs12283611 (large: r2=1, medium: 1 > r2 >= 0.8, small: r2 < 0.8). (B) Single-cell RNA-seq eQTL data for dopaminergic neurons (DA), immature dopaminergic neurons (iDA), late neuron progenitors (LNP), early neuron progenitors (ENP), neuroepithelial-like cells (NEL) and proliferating floor plate progenitors (PFPP). (C) Single-cell ATAC-seq peaks containing a variant in high ld (r2 >= 0.8) with rs12283611. (D). Single-cell ATAC-seq peaks for different cell types. (E) Bulk RNA-seq eQTL results per differentiation timepoint for PRCP. (F) Bulk ATAC-seq peaks separated per differentiation timepoint. (G) HiC data depicting chromatin regions connected by loops at different differentiation timepoints. Related to Figure 7.

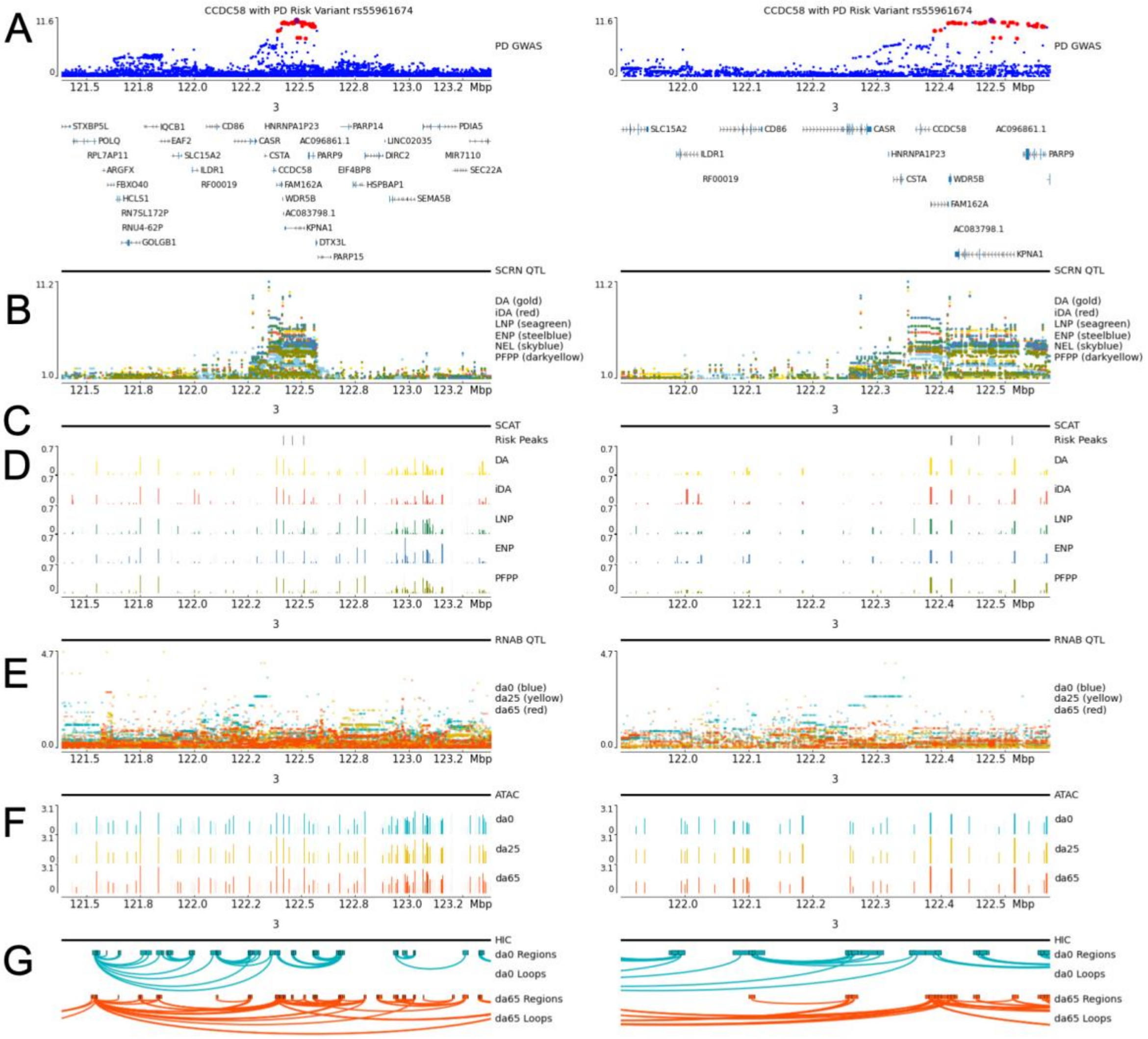


**Supplemental Figure 29.** PD risk locus multi omics figure near CCDC58 on chromosome 3. Tracks represent different data modalities generated or considered in FOUNDIN-PD as different data tracks; figures generated using pyGenomeTracks. The left and right panels of the figures display the same tracks where the left side is a larger region centered on the PD risk locus and the ride side only includes the interval containing the index PD risk variant for this locus and variants in LD with that index variant. (A) GWAS risk for PD in the region. Point size denotes r2 linkage disequilibrium with PD index variant rs55961674 (large: r2=1, medium: 1 > r2 >= 0.8, small: r2 < 0.8). (B) Single-cell RNA-seq eQTL data for dopaminergic neurons (DA), immature dopaminergic neurons (iDA), late neuron progenitors (LNP), early neuron progenitors (ENP), neuroepithelial-like cells (NEL) and proliferating floor plate progenitors (PFPP). (C) Single-cell ATAC-seq peaks containing a variant in high ld (r2 >= 0.8) with rs55961674. (D). Single-cell ATAC-seq peaks for different cell types. (E) Bulk RNA-seq eQTL results per differentiation timepoint for CCDC58. (F) Bulk ATAC-seq peaks separated per differentiation timepoint. (G) HiC data depicting chromatin regions connected by loops at different differentiation timepoints. Related to Figure 7.

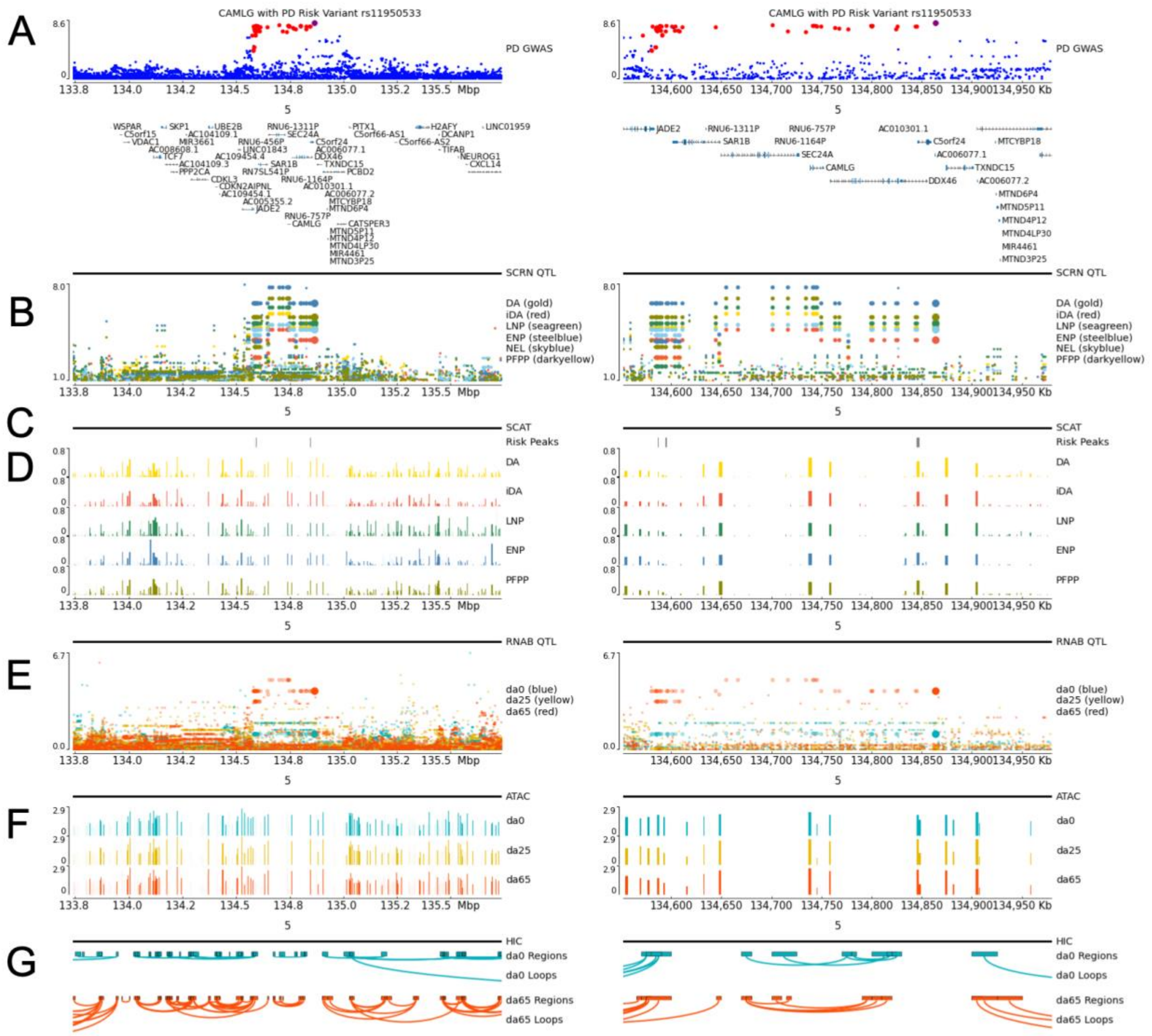

**Supplemental Figure 30.** PD risk locus multi omics figure near CAMLG on chromosome 5. Tracks represent different data modalities generated or considered in FOUNDIN-PD as different data tracks; figures generated using pyGenomeTracks. The left and right panels of the figures display the same tracks where the left side is a larger region centered on the PD risk locus and the ride side only includes the interval containing the index PD risk variant for this locus and variants in LD with that index variant. (A) GWAS risk for PD in the region. Point size denotes r2 linkage disequilibrium with PD index variant rs11950533 (large: r2=1, medium: 1 > r2 >= 0.8, small: r2 < 0.8). (B) Single-cell RNA-seq eQTL data for dopaminergic neurons (DA), immature dopaminergic neurons (iDA), late neuron progenitors (LNP), early neuron progenitors (ENP), neuroepithelial-like cells (NEL) and proliferating floor plate progenitors (PFPP). (C) Single-cell ATAC-seq peaks containing a variant in high ld (r2 >= 0.8) with rs11950533. (D). Single-cell ATAC-seq peaks for different cell types. (E) Bulk RNA-seq eQTL results per differentiation timepoint for CAMLG. (F) Bulk ATAC-seq peaks separated per differentiation timepoint. (G) HiC data depicting chromatin regions connected by loops at different differentiation timepoints. Related to Figure 7.
